# ENNGene: an Easy Neural Network model building tool for Genomics

**DOI:** 10.1101/2021.11.26.424041

**Authors:** Eliška Chalupová, Ondřej Vaculík, Jakub Poláček, Filip Jozefov, Tomáš Majtner, Panagiotis Alexiou

**Affiliations:** Faculty of Science, National Centre for Biomolecular Research, Masaryk University, Brno, Czechia; Central European Institute of Technology (CEITEC), Masaryk University, Brno, Czechia; Faculty of Informatics, Masaryk University, Brno, Czechia

**Keywords:** Deep Learning, Convolutional Neural Network, Recurrent Neural Network, Evolutionary Conservation Score, RNA Secondary Structure, GUI

## Abstract

**Background:** The recent big data revolution in Genomics, coupled with the emergence of Deep Learning as a set of powerful machine learning methods, has shifted the standard practices of machine learning for Genomics. Even though Deep Learning methods such as Convolutional Neural Networks (CNNs) and Recurrent Neural Networks (RNNs) are becoming widespread in Genomics, developing and training such models is outside the ability of most researchers in the field.

**Results:** Here we present ENNGene - Easy Neural Network model building tool for Genomics. This tool simplifies training of custom CNN or hybrid CNN-RNN models on genomic data via an easy-to-use Graphical User Interface. ENNGene allows multiple input branches, including sequence, evolutionary conservation, and secondary structure, and performs all the necessary preprocessing steps, allowing simple input such as genomic coordinates. The network architecture is selected and fully customized by the user, from the number and types of the layers to each layer's precise set-up. ENNGene then deals with all steps of training and evaluation of the model, exporting valuable metrics such as multi-class ROC and precision-recall curve plots or TensorBoard log files. To facilitate interpretation of the predicted results, we deploy Integrated Gradients, providing the user with a graphical representation of an attribution level of each input position. To showcase the usage of ENNGene, we train multiple models on the RBP24 dataset, quickly reaching the state of the art while improving the performance on more than half of the proteins by including the evolutionary conservation score and tuning the network per protein.

**Conclusions:** As the role of DL in big data analysis in the near future is indisputable, it is important to make it available for a broader range of researchers. We believe that an easy-to-use tool such as ENNGene can allow Genomics researchers without a background in Computational Sciences to harness the power of DL to gain better insights into and extract important information from the large amounts of data available in the field.

## Background

Artificial Neural Networks (ANNs) are a family of Machine Learning (ML) algorithms, which learn complex tasks via the connection of simple artificial neurons. ANNs have been used for a multitude of tasks since their inception more than 50 years ago [1], such as image and signal processing, natural language processing and translation, and many more. In recent years, the field of ANNs has undergone a revolutionary change, with the adoption of Deep Neural Networks, or Deep Learning (DL) [2]. These types of ANNs utilize a large number of stacked artificial layers of neurons in order to learn increasingly complex representations of input data. DL consistently outperforms other ML methods in cases where moderately large training sets are available.

Some of the commonly used networks for Deep Learning are Convolutional Neural Networks (CNNs) and Recurrent Neural Networks (RNNs). CNNs make use of Convolutional neural layers to effectively model specific aspects of input samples while reducing the number of parameters that need to be trained. That allows training with fewer input data, together with the development of deeper architectures. RNNs utilize different architectures such as Long-Short Term Memory (LSTM) and Gated Recurrent Units (GRU) to introduce a temporal memory capability. In Genomics, DL models were quickly applied to problems such as the prediction of biomolecule binding specificities [3], prediction of the effect of non-coding variants [4], learning the functional activity of DNA sequences [5], and many more [6].

Training DL models from scratch requires high-level programming skills and an understanding of DL programming libraries. Recently, projects such as pysster [7], Selene [8], or Janggu [9] have been developed to standardize the usage of Deep Learning in Genomics and lower the prerequisite for training DL models for genomics data. Pysster [7] is a Python package focusing on CNNs for biological sequence data. Pysster enables learning sequence and structure motifs and visualizing them along with information about their positional and class enrichment. Selene [8] is a PyTorch-based library for working with sequence-based DL models. It offers workflows for applying an existing model, retraining it on new data, or developing new model architectures. Though it is meant to be run by simple scripts, the input is provided via extensive configuration files. When creating a new architecture, users must be able to define it in a separate Python file properly. Together, this is a rather intricate process that can discourage researchers not versed in programming or scripting. Finally, Janggu [9] is a library offering support for both Keras and PyTorch [10] backend, together with unified dataset objects simplifying data acquisition and preprocessing. It supports visualization options, including genomic tracks and input feature importance via Integrated Gradients. While these tools offer a measure of facilitation for developing DL models, they still expect users to have at least intermediate programming skills.

We have experienced the need for an accessible and user-friendly framework that could allow Genomics researchers with minimal programming skills to access the power of applying DL on their preferred dataset. With this motivation, we have developed ENNGene, a tool utilizing a user-friendly Graphical User Interface (GUI) to handle data preprocessing, model building, training, and evaluation, as well as interpretation of the predicted results (Fig.1). ENNGene allows users to select among up to three input branches - genomic sequence, predicted RNA secondary structure, and, unlike any other DL implementation for Genomics, phylogenetic conservation scores. In the last decade, most of the progress in the DL field has been concentrated on finding better performing, easier-to-train deep neural networks. In practice, it is also essential to be able to verify the results and validate the process that led to them. That is especially important in applications such as the Genomic and Biomedical fields, where the model's reliability must be guaranteed. ENNGene offers an interpretation of trained models based on a novel implementation of the Integrated Gradients method [11] that can be applied on multi-branch networks.

**Fig. 1:**
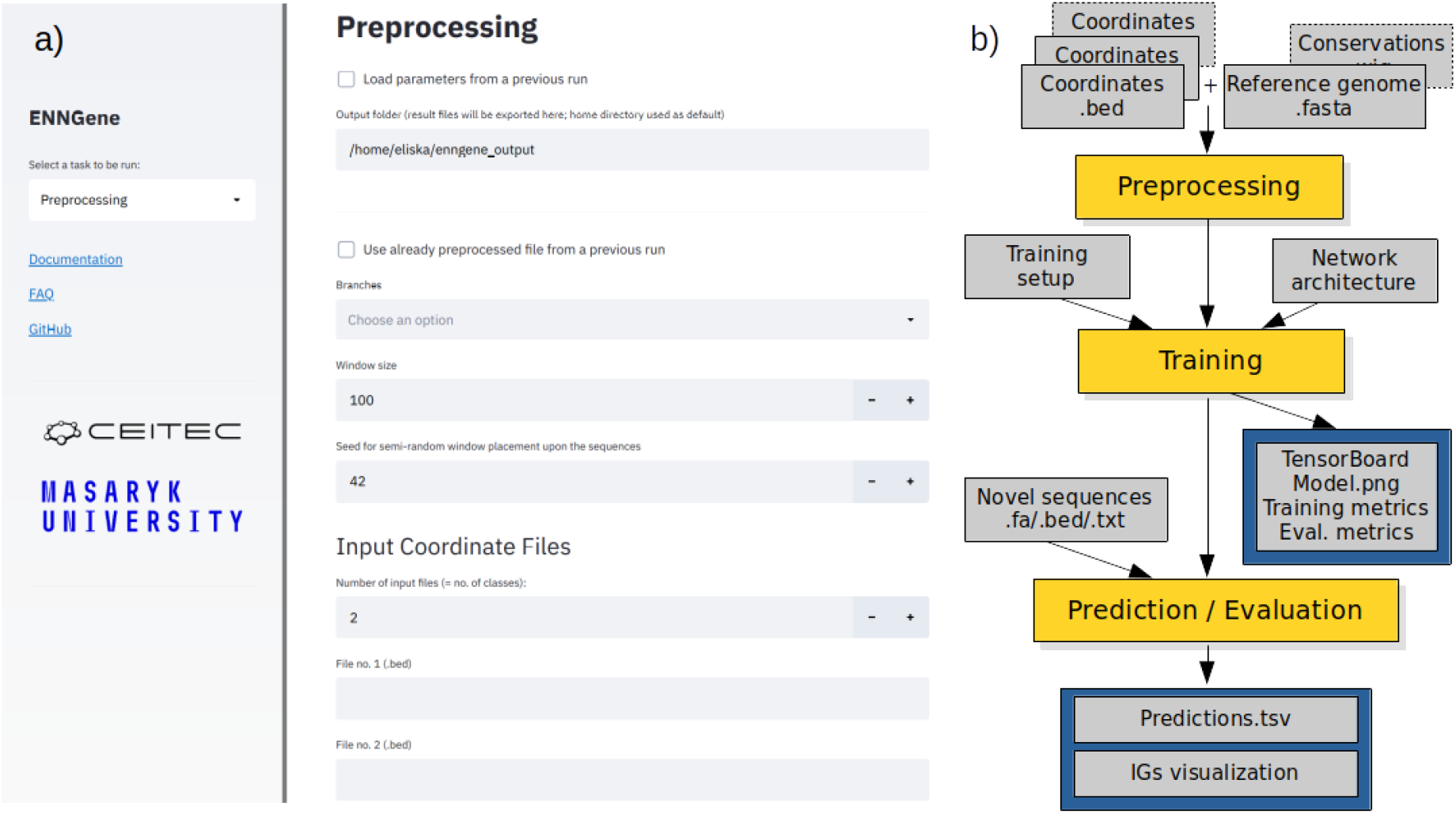
a) The Graphical User Interface (GUI) of ENNGene - ENNGene is fully operated via the GUI. Users define the input parameters using simple interactive elements, such as dropdown menus or checkboxes. Warnings and hints are displayed via the GUI in a user-friendly way directly as the user interacts with it. Web browser being at the basis of the GUI, interactive plots or results are visualized immediately throughout or after the calculations. b) Simplified data flow - ENNGene comprises multiple subsequent modules with separate functionality, covering the whole process from input preparation and network architecture definition to model evaluation and interpretation.

We demonstrate the functionality of ENNGene on the use case of a well-known benchmark dataset based on the classification of RNA Binding Protein (RBP) binding sites [12]. On this benchmark, we show that models developed using ENNGene, without specific hand-crafted features based on domain knowledge, match or outperform state-of-the-art methods created explicitly for this task.

## Implementation

ENNGene uses the open-source Streamlit framework (https://streamlit.io/) and is built atop TensorFlow [13], one of the most popular DL backends. It runs locally on either Central Processing Units (CPUs) or Graphics Processing Units (GPUs), which offer a considerable speed-up when training larger networks [14]. Our major considerations in the implementation of ENNGene were ease of use, generic functionality on various genomic problems, and reproducibility of results. The software consists of four linearly connected modules: (a) preprocessing module, (b) training module, (c) prediction module, and (d) evaluation module.

### User Input

The minimal user input is a Browser Extensible Data (BED) file containing DNA/RNA sequences and a genome or transcriptome (fasta) reference file. An additional PhyloP [15] reference file must be provided if the user wishes to utilize the evolutionary conservation. Such reference files are readily available for most organisms from public repositories such as Ensembl [16] or UCSC Genome Tables Browser [17]. Optionally, the user may also input a log file (Yaml) from a previous ENNGene run, instantly reproducing the input configuration. This way, users can guarantee full reproducibility across different instances of ENNGene, provided the same input files are available. To avoid re-mapping the same inputs to the same reference multiple times, ENNGene provides an option to save and reuse mapped files between runs.

### Module 1: Preprocessing

In the first module, data is preprocessed into a format convenient for CNN input. The user may select one or more input types engineered from the given interval files. (a) Sequence – one-hot encoded RNA or DNA sequence, obtained by mapping given intervals to the reference genome or transcriptome fasta file. (b) RNA secondary structure is based on the mapped sequences and computed by the ViennaRNA2 package [18]. When calculating the secondary structure, users can choose the number of CPU cores dedicated to the computation. (c) The evolutionary conservation score is mapped similarly to the sequence, with the user-provided phylogenetic conservation file as a reference.

As required by CNNs, all input sequences must be of the same length. One strategy for ensuring same length inputs is N or 0 padding [19, 20]. However, such an approach runs the danger of introducing a blind spot or even learning the padding artifact above the legitimate features when sequences of different classes do not have the same length distribution [21]. In the case of ENNGene, we avoid this by extending or trimming the provided genomic coordinates to the same length, thus ‘padding’ our intervals with actual neighboring sequences and other features.

The training of CNNs can be affected by an imbalance in class sizes [22]. It is common practice to artificially balance the various classes in training sets by downsampling the more populous classes. ENNGene makes this optional preprocessing step easy by allowing users to define a reduction by an absolute or relative dataset size (e.g., 5,000 out of 10,000 sequences, or 0.5 ratio).

Following this step, datasets are split into ‘training’, ‘validation’, ‘evaluation’ (testing), and ‘black-box’ (left-out testing) randomly at a user-defined ratio or based on reference chromosome number. The ‘training’ and ‘validation’ datasets are used throughout the training process. The ‘evaluation’ dataset is used for model evaluation directly after the training. The last one, the ‘black-box’ dataset, is optional and is never seen by any part of the training or evaluation modules of ENNGene. It is a truly left-out dataset that can be used for final evaluation after multiple rounds of hyperparameters tuning and architecture exploration in order to avoid overfitting the evaluation dataset. Splitting the sequences based on the chromosome they are found on also ensures that each of these datasets comes from entirely unrelated genomic regions and that there is no possibility for one part of the process encountering a left-out locus, for example, when using input files with nearly overlapping genomic loci.

### Module 2: Training

In the second module, the user can use the GUI to define the neural network architecture and training hyperparameters: (a) batch size, (b) learning rate, and (c) choose one of three available optimizers: stochastic gradient descent (SGD) - preset with Nesterov’s momentum to 0.9 [23], RMSprop [24], or Adam [25]. If SGD is selected, two additional learning rate options become available, using a learning rate scheduler or applying one cycle policy [26]. Users may also define a specific number of training epochs or apply the early stopping callback, which stops the training when there is no more improvement on the validation dataset. We have opted to simplify hyperparameter selection as much as possible while not compromising the power of the produced models.

ENNGene enables the building of CNNs consisting of convolutional layers followed by fully connected layers, or hybrid CNN-RNNs. In the latter case, the convolutional layers are first followed by recurrent layers - either GRU [27] or LSTM [28], finally followed by dense layers. The details of the network architecture are defined per each section separately, having one section for each selected branch corresponding to the preprocessed input types and one section for the common part of the network following the concatenation of the branches.

After setting up the number and types of the layers, the layer parameters can be customized. For each layer, users can pick a dropout rate [29], choose to apply the batch normalization [30], and set the layer-specific options - the number and size of the convolutional filters, the number of dense or recurrent units, and the application of a bidirectional wrapper on the recurrent layers. In accordance with the focus of the ENNGene on classification problems for two or more classes, the loss function is preset to categorical cross-entropy, together with the last layer set as the softmax activation function with the number of units corresponding to the number of classes.

Models created by ENNGene are trained using the TensorFlow framework with the support of GPU acceleration. The GPU is automatically detected and used when available on the machine, providing considerable speed up during the training phase. Throughout the training, users can monitor the progress, together with the development of the accuracy and loss metrics on an interactive plot. These metrics are also plotted and exported for later use.

After the training, the resulting model is directly evaluated on the user-defined evaluation dataset. Above the basic metrics, the receiver operating characteristic (ROC) and precision-recall curves, both adjusted for the multi-class classification problems, are calculated and exported in the form of plots and tables. Furthermore, TensorBoard [13] files allowing visualization and browsing of the model architecture, as well as metrics comparison between multiple runs, can be produced.

### Modules 3&4: Prediction and Evaluation

The final two modules allow the user to evaluate a previously trained model on a different dataset (Evaluation) or apply the model to classify novel data samples (Prediction). The two modules share most of the functionality; therefore, they will be addressed together in this last section. At this point, the user may upload any pre-trained model, either produced by ENNGene or not, and a set of genomic loci to be evaluated either with known class labels (Evaluation) or not (Prediction).

Loci to be evaluated are preprocessed in the same way described in the first module. For evaluation purposes, the black-box dataset prepared during the preprocessing of the original data can be used. If the trained model does not use a conservation branch, then even simple fasta files containing sequences of the correct length may be used for prediction.

Calculated predictions are exported as probability scores for each class, with the highest-scoring class highlighted. ENNGene does not return a single ‘predicted class’, as that can vary based on the subjective choice of a threshold for each class. Optionally, Integrated Gradients (IG) [11] are calculated for each sample and exported as a list of scores for custom visualization. At the same time, the top ten predictions per class are directly displayed in the browser (Fig.2 (c)). The IG technique is based on calculating the difference between the baseline, a vector of zeros in our case, and an input sequence. That dependency is the core of the attribution and is expressed as the color adjustment of each nucleobase across all the input types. The higher the attribution of the sequence to the prediction, the more pronounced red is its color. On the other hand, blue means a low level of attribution. The attribution visualization can be used for auxiliary evaluation and debugging of the model.

**Fig.2:**
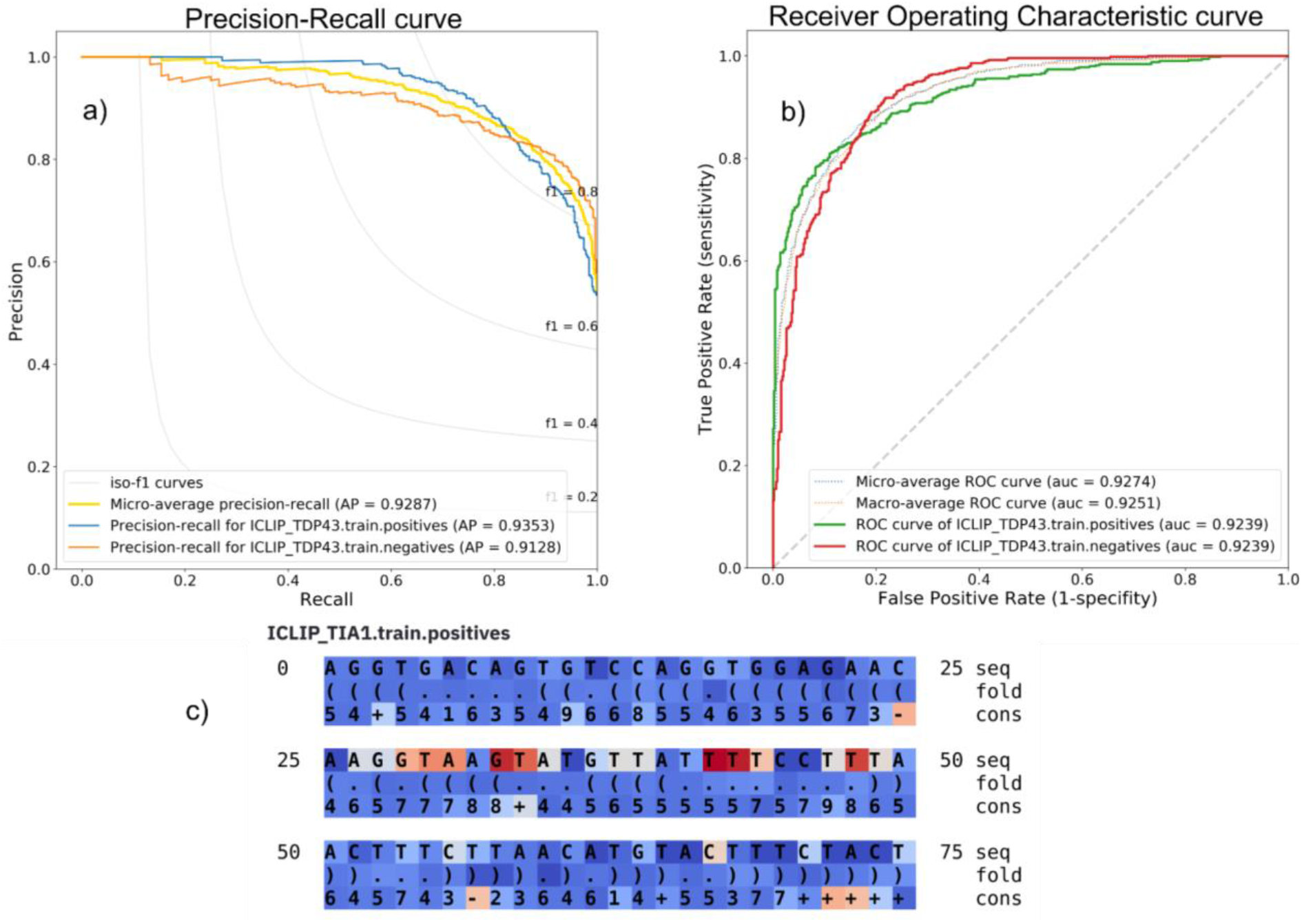
a) Precision-recall curve - the precision-recall metric indicates the relationship between the model’s positive predictive value (precision) and sensitivity (recall) at various thresholds. b) Receiver Operating Characteristic (ROC) curve - the ROC metric is calculated as a ratio between the true positive rate and the false positive rate at various thresholds. Both the metrics, precision-recall and ROC calculated by ENNGene, are adjusted for multi-class classification problems and thus can be applied to models with any number of classes. Both curves and other metrics (accuracy, loss, AUROC) are a standard part of exported results after a model evaluation, optionally with Integrated Gradients’ scores. c) Integrated Gradients visualization - IG scores of ten sequences with the highest predicted score per class are directly visualized in the browser. Scores are displayed in separate rows for each input type used - sequence, secondary structure, and conservation score. The higher the nucleotide’s attribution to the prediction of a given class, the more pronounced is its red color. On the other hand, the blue color means a low level of attribution.

**Fig. 3:**
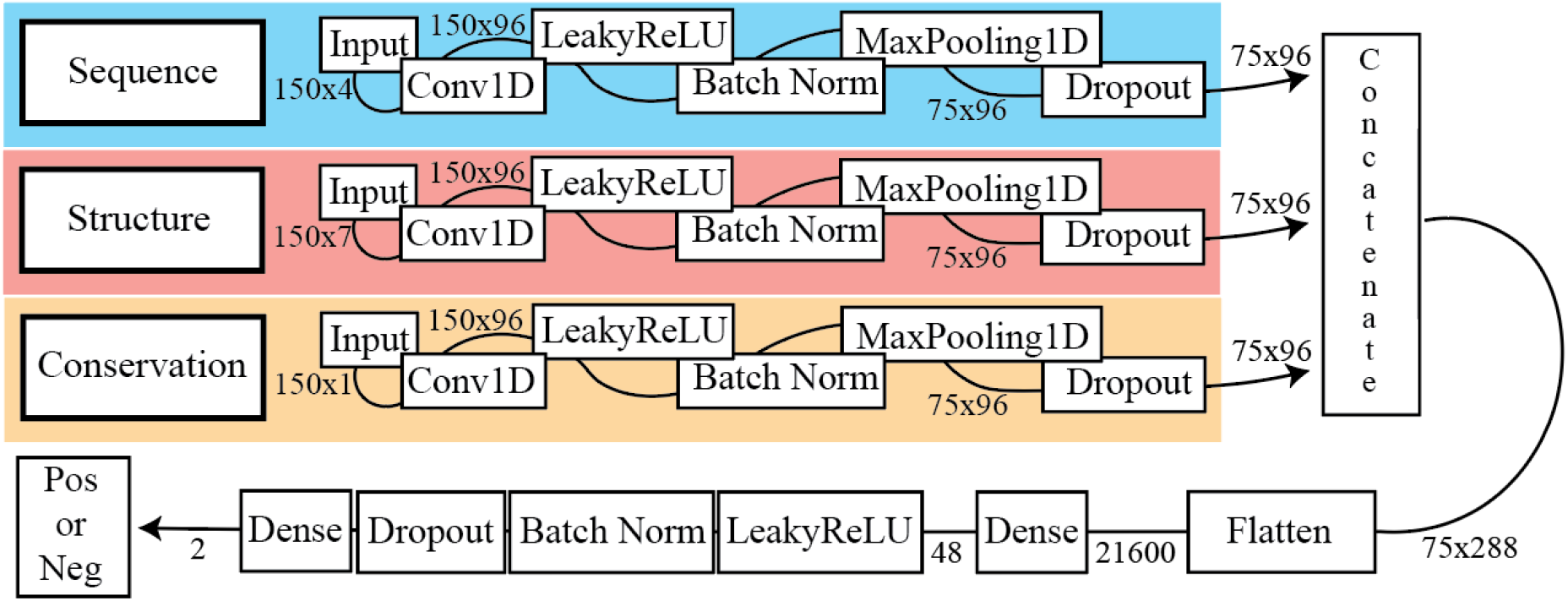
Simplified representation of model architecture. The model in this example was trained on 150 nt long sequences using all three available input types - sequence, secondary structure, and conservation score - each represented by a separate model branch. After the network extracts information from the separate inputs, the branches are concatenated, and the network continues learning interdependencies by looking at the combined information via dense or recurrent layers. Boxes represent individual layers, while the adjacent numbers indicate the data dimensionality. A plain graphical representation of the network architecture is produced and exported by ENNGene for every trained model.

## Results and Discussion

ENNGene is a generic tool created to enable researchers to produce state-of-the-art DL models with minimal programming knowledge. It was developed with a focus on ease of use, reproducibility, and interpretability while avoiding compromising the power of the produced models.

### Ease of use and reproducibility

The entirety of ENNGene’s code is freely and openly available at GitHub at https://github.com/ML-Bioinfo-CEITEC/ENNGene (also see Additional file 1) under an open-source GNU Affero General Public License. ENNGene runs on a Linux system and was extensively tested on the Ubuntu distribution. Using the Anaconda package manager, we ensure containerization and safe un/installation of the application. A step-by-step guided installation script, as well as extensive documentation, can be found at the project’s code repository and as a supplementary file accompanying this publication (Additional file 2).

We have taken various steps to improve the user experience. The application verifies every provided input information. Specific warnings or hints are returned to the user immediately during the parameter set-up or just before any further calculations are started. All the user input, warnings, and errors are logged throughout the whole session. After the session is finished, the log file is placed in the user-specified output folder. If an error occurs during the session, the user is notified of the location of the log file containing the additional details.

To ensure full reproducibility of the results, ENNGene logs all the parameters set by the user and exports them as a Yaml file. The file can be imported into the application at any time in the future, immediately setting all the parameters for the chosen task. This way, users can recreate all their results or quickly tune chosen hyperparameters. Above that, all the tracked parameters are exported into one .tsv file, shared across all the application sessions within the same output folder, with one line per session. This file can significantly streamline the comparison of multiple models, as the user has all the preprocessing and training parameters used to create each of the models in one place.

### Case Study: RBP24 classification - comparison to the state of the art

To demonstrate the functionality of ENNGene, we chose the problem of the RNA-binding proteins (RBPs) target site classification. CNNs were applied to the RBP target site classification problem for the first time in 2015 [3]. Since then, there have been many novel approaches, including several different network architectures. With the target RNA sequence as the primary feature, predicted secondary structure [7, 20, 31–34] and transcript region type [35, 36] are the other most commonly used features. Up to date, several different network types have been applied to the RBP binding site classification problem. Those include the CNN [7, 32], RNN [37], or often a combination of the two [20, 32, 33, 36, 38], deep belief network (DBN) [31, 35], deep residual network [39], and attention network [34]. ENNGene allows the users to use the two most widely applied architectures - CNN and hybrid CNN-RNN networks. In most cases, one model per RBP is trained. In a few exceptions, a multi-label classifier is built for all the proteins [36], for example, by applying classifier chains to learn the correlation between labels [40]. Using ENNGene, we have applied the two most commonly used network types (CNN and hybrid CNN-RNN) on the RBP24 dataset introduced in GraphProt [12], producing one model for each RBP in the dataset.

We selected this widely used benchmark dataset as it has already been used for benchmarking by several state-of-the-art methods [3, 19, 31, 33, 39]. The RBP24 dataset consists of 21 RNA-binding protein target sites from 24 CLIP-seq experiments. Since ENNGene works with input bed files, we extracted the interval coordinates from the sequence header from the original RBP24 dataset (see the extracted genomic coordinates in the Additional files 3 and 4; the original fasta files can be obtained from GraphProt repository at http://www.bioinf.uni-freiburg.de/Software/GraphProt/). Following the original dataset, intervals were mapped to the human (hg19) genome reference on a 150 nt long window centered at the initial coordinates. The length of 150 nucleotides has been previously used in several RBP target site predictors [31] and has been proven to be the best choice for secondary structure prediction [41]. The base-wise conservation scores by PhyloP [15] from the PHAST package [42], created by multiple alignments of 99 vertebrate genomes to the human genome, were obtained from the UCSC file storage. Only the PTBv1 dataset was mapped to the older (hg18) genome reference and the corresponding PhyloP files (with 43 vertebrate genomes aligned) according to the original dataset specification. ENNGene being a generic tool works well with any genomic assembly and depth of conservation file.

The number of samples in the RBP24 dataset widely differs between the experiments, with the ratio between positive and negative classes always close to 1:1. We used the original training files for model training and parameter search purposes, dividing them into training, validation, and testing datasets. The original testing samples were left entirely out as a ‘black-box’ dataset. We chose the best models based on the Area Under Receiver Operating Characteristic curve (AUROC) metric upon the testing dataset while prioritizing the simplest architecture available. We performed a human-guided search of selected hyperparameters since fully covering the data and model hyperparameter space by grid search would be too time-consuming for a simple demonstration. The tuned parameters were number and combination of branches, batch size, number of epochs, optimizer, learning rate, number of layers per section, as well as the type of the layer and specific parameters per layer (number of units/filters, bidirectionality, kernel size, dropout rate, batch normalization). The models were trained for different numbers of epochs, optionally using the early stopping with parameters set to patience = 10 and delta = 0.01 to avoid overfitting. A complete list of the hyperparameters used for each final model can be found in Additional file 5.

The left-out files were used only for the final evaluation of the already tuned models. We show that using ENNGene we were able to produce models that perform to the current state-of-the-art standard. For more than half of the experiments in the RBP24 dataset, ENNGene models outperform the state of the art, producing an improved average AUROC value of 0.947 on the whole RBP24 dataset (Table 1, Additional file 6). All the final trained models can be found at the project’s GitHub repository.

**Table 1.** AUROC values from the evaluation of the final models, together with the RBP24 dataset properties. Models created using ENNGene outperform the state-of-the-art tools on more than half of the experiments in the RBP24 dataset while also improving the average AUROC value on the whole dataset. The AUROC values of the other tools are taken from the original publications. The highest AUROC score reached for the experiment is highlighted in bold.

| Protein | RBP-24 |  |  |  |  |  |  |  |  |  |
| --- | --- | --- | --- | --- | --- | --- | --- | --- | --- | --- |
|  | Dataset info |  |  | Results (AUROC) |  |  |  |  |  |  |
|  | Clip method | # positives | # negatives | GraphProt | deepnet-rbp | DeepBind | iDeepV | iDeepE | DeepRKE | ENNGene |
| **Ago1-4** | PAR-CLIP | 36902 | 31310 | 0.895 | 0.881 | 0.919 | 0.925 | 0.915 | **0.932** | 0.927 |
| **Ago2** | HITS-CLIP | 48095 | 44251 | 0.765 | 0.809 | 0.879 | 0.886 | 0.884 | 0.9 | **0.94** |
| **ALKBH5** | PAR-CLIP | 1213 | 1197 | 0.68 | 0.714 | 0.668 | 0.643 | 0.758 | 0.74 | **0.877** |
| **C17ORF85** | PAR-CLIP | 1860 | 1849 | 0.8 | 0.82 | 0.755 | 0.74 | 0.83 | 0.824 | **0.916** |
| **C22ORF28** | PAR-CLIP | 9369 | 9136 | 0.751 | 0.792 | 0.809 | 0.823 | 0.837 | 0.832 | **0.908** |
| **CAPRIN1** | PAR-CLIP | 8140 | 7901 | 0.855 | 0.834 | 0.888 | 0.824 | 0.893 | 0.869 | **0.896** |
| **ELAVL1** | HITS-CLIP | 8595 | 8436 | 0.955 | 0.966 | 0.98 | 0.966 | 0.964 | 0.978 | **0.991** |
|  | PAR-CLIP (A) | 27275 | 23974 | 0.959 | 0.966 | 0.972 | 0.973 | **0.988** | 0.978 | 0.965 |
|  | PAR-CLIP (B) | 9464 | 9283 | 0.935 | 0.961 | 0.961 | 0.962 | 0.971 | 0.98 | **0.987** |
|  | PAR-CLIP (C) | 125202 | 113686 | 0.991 | 0.994 | 0.989 | 0.99 | 0.979 | **0.996** | 0.989 |
| **EWSR1** | PAR-CLIP | 16292 | 14720 | 0.935 | 0.966 | 0.969 | 0.962 | 0.969 | 0.971 | **0.972** |
| **FUS** | PAR-CLIP | 34581 | 31480 | 0.968 | 0.98 | 0.983 | 0.976 | 0.985 | **0.988** | 0.977 |
| **hnRNPC** | iCLIP | 21472 | 19794 | 0.952 | 0.962 | **0.979** | **0.979** | 0.976 | 0.978 | 0.977 |
| **IGF2BP1-3** | PAR-CLIP | 8539 | 6838 | 0.889 | 0.879 | 0.939 | 0.923 | **0.947** | 0.943 | 0.946 |
| **MOV10** | PAR-CLIP | 13793 | 12987 | 0.863 | 0.854 | 0.899 | 0.896 | 0.916 | 0.92 | **0.922** |
| **PUM2** | PAR-CLIP | 9116 | 8227 | 0.954 | 0.971 | 0.964 | 0.965 | 0.967 | 0.965 | **0.972** |
| **PTB** | HITS-CLIP | 44574 | 43700 | 0.937 | **0.983** | 0.944 | 0.936 | 0.944 | 0.953 | 0.954 |
| **QKI** | PAR-CLIP | 10276 | 9142 | 0.957 | 0.983 | 0.973 | 0.965 | 0.97 | 0.975 | **0.984** |
| **SFRS1** | HITS-CLIP | 19438 | 17195 | 0.898 | 0.931 | 0.929 | 0.905 | **0.946** | 0.945 | 0.939 |
| **TAF15** | PAR-CLIP | 7298 | 6606 | 0.97 | 0.983 | 0.978 | 0.978 | 0.976 | **0.985** | **0.985** |
| **TDP43** | iCLIP | 92031 | 75079 | 0.874 | 0.876 | 0.93 | 0.935 | 0.945 | **0.954** | 0.924 |
| **TIA1** | iCLIP | 18049 | 16135 | 0.861 | 0.891 | 0.929 | 0.941 | 0.937 | 0.942 | **0.959** |
| **TIAL1** | iCLIP | 42332 | 36652 | 0.833 | 0.87 | 0.922 | 0.929 | 0.934 | 0.946 | **0.95** |
| **ZC3H7B** | PAR-CLIP | 20962 | 20018 | 0.82 | 0.796 | 0.875 | 0.883 | 0.907 | **0.914** | 0.862 |
| **AVERAGE AUROC:** |  |  |  | 0.887 | 0.903 | 0.918 | 0.913 | 0.931 | 0.934 | **0.947** |

### Importance of network branches beyond sequence

Easily producing Deep Learning models at the state-of-the-art level is the primary objective of ENNGene. However, we are also interested in exploring additional features beyond sequence, namely evolutionary conservation and predicted RNA secondary structure. Previously [43], we have used a multi-branch approach with the same features to identify small RNA loci, outperforming the state of the art. Both the conservation and secondary structure branches were shown to be important for that task. In the RBP24 dataset presented here, we investigated networks of various configurations, starting from the simple sequence, adding either evolutionary conservation or secondary structure branches, and finally, all three branches together. We identified that adding an evolutionary conservation branch improved the performance of all but five protein datasets (FUS, HNRNPC, ELAVL1 - PAR-CLIP C, PTB, and TDP43) that had similar performance with or without the conservation feature. On the contrary, we had not registered improvement in any model when the secondary structure was included.

Evolutionary conservation features have been extensively used by machine learning classifiers predating the beginning of DL, along with many other features such as amino acid composition and hydrophobicity, predicted secondary structure, and other global protein features [44]. An essential concept in CNNs is that using convolutional layers can bypass the process of derivative feature generation, as the network itself learns how to derive complex features from raw data. However, unlike, for example, a predicted secondary structure, the conservation information can not be derived from the given sequence, and as such, it may provide the network with additional information. The group of authors behind the iDeepV and iDeepE methods [19, 39] showed that in both approaches, they were able to reach state-of-the-art performance on the RBP24 dataset without using secondary structure information. DeepCLIP [38] presents a hybrid CNN-RNN network that outperforms structure-based models, as well as the iDeepV and iDeepE models, using sequence data alone. The authors hypothesize that context-dependency can be better modeled implicitly by the recurrent layers rather than by predefined features, such as the predicted secondary structure. However, other models such as DeepRKE [33], Deepnet-rbp [31], and DLPRB [20] show a performance improvement when utilizing secondary structure features in a small fraction of proteins. That emphasizes the need for individually designed architectures for each protein separately. In the case of evolutionary conservation, which is not a derivative feature, it is clear that substantial additional information is indeed given to the network under training. Conservation features have not been used by other DL algorithms tackling this question, probably because of the difficulty of retrieving and preprocessing conservation data. ENNGene’s preprocessing module takes care of this process for the user and makes this important feature available.

It has been previously shown that the combination of derived/hand-crafted and learned features can lead to better performance [45]. We have avoided adding more hand-crafted features to retain the generality of ENNGene to deal with genomic questions. We decided to retain the secondary structure feature as it is a commonly used feature that we have found useful in other applications of CNNs [43].

### Interpretation of predictions

To gain an insight into what a DL model has learned, many studies utilizing CNNs use a heuristic based on convolution kernel analysis [3, 20, 32, 38]. This approach is characteristic for the Genomics field, as it allows one to visualize a learned pattern as a pseudo motif-representing position weight matrix [46]. However, for the full understanding of the relationship between an input sequence and an output prediction, simply analyzing filters of one convolutional layer is not sufficient.

To evaluate the importance of each nucleotide, [3] applied input modification, also known in the field as an in silico mutagenesis. In this approach, each nucleotide is substituted by every other possible nucleotide, and a prediction is re-calculated for every ‘mutated’ sequence to identify the essential parts. Such an exhaustive input perturbation involves very high computational costs. Increasing the number of nucleotides to be perturbed exponentially increases the computational time needed to calculate effects. Therefore, in most cases, only a part of the sequence is modified, based either on random selection or on a previously defined motif, and important parts of the sequence might be completely ignored [6].

Gradient-based approaches [47–53], to the contrary, are not only more computationally efficient but provide information about the whole sequence, allowing to find previously unexpected essential sections in the sequence. One issue of the earlier methods [47, 48] is neuron saturation. Given a sequence with two binding regions, such an algorithm would assign a low score to both regions, as each such region is not individually important enough for the prediction. To address the saturation problem, methods like DeepLift [52] and Integrated Gradients [11] use a background reference to properly distinguish the crucial elements from noise. Integrated Gradients, the method adopted by ENNGene, gained popularity due to its mathematical and theoretical justifications and computational efficiency compared to alternative approaches. IGs can be applied to any differentiable model, not strictly focusing on the CNNs, and are suitable for large networks and feature spaces without the undesirable computational overhead. Only one recent RBP target site classifier [36] utilizes the IGs to see what parts of the sequence and region, the two input types adopted in the study, were responsible for the predicted category. Similarly, for ENNGene, we use IGs to visualize important parts of the input within all branches of the trained model.

### Future Development

At this point, ENNGene allows the production of state-of-the-art multi-branch CNN and CNN-RNN models via a user-friendly interface while providing one level of interpretation of the produced results. Despite the ease of training new models with ENNGene, automatic identification of the optimal model architecture and hyperparameter search would be an important future step. Neural Architecture Search (NAS) is an automated search of an optimal architecture on given data, using predefined building blocks. Using NAS, the network building blocks are handled similarly to other hyperparameters. Since around 2015, there are already several different NAS techniques [54]. It has been shown that NAS can be on par or outperform hand-designed architectures [55, 56]. Recently, NAS was implemented on genomics data, improving processing time and complexity of use [57]. Even with the current computational power at hand, it is still a very resource-demanding process outside most research groups’ capabilities. We aim to incorporate some type of NAS in the future of ENNGene, as computational resources become more readily available and methods for NAS more efficient.

Currently, the process of manually tuning hyperparameters is fairly simple. Users can use the option to load the parameters from any previous session, change just the chosen parameters without setting up the rest, and start training again within moments. Nevertheless, we would like to add a separate module dedicated to automated hyperparameter tuning. The new module would provide options such as random or grid search of user-defined parameter space.

Finally, to support a broader user base in the future, we are considering setting up a dedicated server with the most common genome references available. That way, users would need to provide only the interval files, making the usage of ENNGene even easier. Depending on the current costs and funds available, the server might provide the GPU power, speeding up the NN training for the users with no other access to it.

## Conclusions

As genomic data generation accelerates, researchers in the field will increasingly require powerful but easy-to-use data science and machine learning methods. In this work, we have presented ENNGene, a tool that allows the development and interpretation of Deep Learning models without advanced programming knowledge. We believe that ENNGene will empower Genomics researchers to explore their data using the powerful models that ENNGene can produce. We have demonstrated these abilities by building state-of-the-art models for a well-known and competitive benchmark of RNA Binding Protein target sites. It is our intention to support ENNGene and its user base with continuous development and improvement over the following years, making it an important tool for the Genomics research community.

## Availability and requirements

**Project name**: ENNGene

**Project home page**: https://github.com/ML-Bioinfo-CEITEC/ENNGene

**Operating system(s)**: Linux

**Programming language**: Python 3

**Other requirements**: Anaconda, web browser

**License**: GNU Affero General Public License v3.0

**Any restrictions to use by non-academics**: None above the license

## List of abbreviations

AUROC: Area Under the Receiver Operating Characteristic curve
BED: Browser Extensible Data
BLSTM: Bidirectional Long Short-Term Memory
CNN: Convolutional Neural Network
CPU: Central Processing Unit
DBN: Deep Belief Network
DL: Deep Learning
GPU: Graphics Processing Unit
GRU: Gated Recurrent Unit
IGs: Integrated Gradients
LSTM: Long Short-Term Memory
ML: Machine Learning
NAS: Neural Architecture Search
RBP: RNA-Binding Protein
RNN: Recurrent Neural Network
ROC: Receiver Operating Characteristic
SGD: Stochastic Gradient Descent

## Declarations

### Ethics approval and consent to participate

Not applicable.

### Consent for publication

Not applicable.

### Availability of data and materials

All data generated or analysed during this study are included in the supplementary information files.

### Authors’ contributions

EC created the ENNGene application. FJ and JP implemented the Integrated Gradients for ENNGene. OV created the installation script. EC and OV conducted the RBP24 use case. EC, OV, and TM tested the application. EC and PA prepared the manuscript. All authors read and approved the final manuscript.

## Acknowledgments

Not applicable.

## Authors’ information (optional)

## Supplementary Information

**Additional file 1**

ENNGene v1.1.2

The latest version of the ENNGene application available on the day of the publication submission. The current version of the publication can always be found at the project’s GitHub repository at https://github.com/ML-Bioinfo-CEITEC/ENNGene.

**Additional file 2**

Documentation

Complete documentation of the current version of the ENNGene application (v1.2.2) in pdf format. Each module and its functions, together with all the parameters and available options, are described in the documentation.

**Additional file 3**

RBP24 input files

Bed files containing the genomic coordinates extracted from the RBP24 fasta files. Data from these files were split into training, validation, and testing datasets. The original fasta files are available at http://www.bioinf.uni-freiburg.de/Software/GraphProt/.

**Additional file 4**

RBP24 leave-out files

Bed files containing the genomic coordinates of the leave-out data, used only for the final evaluation of the best performing models. The resulting AUROC values are reported in table 1 and were used for comparison with the other publications and their reported AUROC values.

**Additional file 5**

RBP24 model parameters

Complete list of the ENNGene parameters used to obtain the final models. Parameters from Preprocessing, Training, and Evaluation modules are reported and ensure full reproducibility of the results.

**Additional file 6**

RBP24 ROC curves

ROC curves generated by ENNGene for each experiment in the RBP24 dataset. The ROC curves are based on the evaluation of the final models on the leave-out data. The AUROC values of the corresponding ROC curves were used for comparison with the other publications and their reported AUROC values.

