## Additional file 6 for "ENNGene: an Easy Neural Network model building tool for Genomics"

### ROC curves of final models for RBP24 dataset

A Receiver Operating Charateristic (ROC) metric is calculated as a ratio between the true positive rate and the false positive rate at various thresholds. The ROC curves generated by ENNGene are adjusted for multiclass classification problems, and thus can be applied to any number of classes.

For the RBP24 dataset, we chose the best models based on the Area Under Receiver Operating Characteristic curve (AUROC) metric upon the testing dataset (obtained from the data in the Supplementary file 3), while prioritizing the simplest architecture available. The ROC curves below are based on the evaluation of those models on the leave-out data (Supplementary file 4). The AUROC values of the corresponding ROC curves were used for the comparison with the other publications and their reported AUROC values.

Fig.1 AGO2 ROC curve

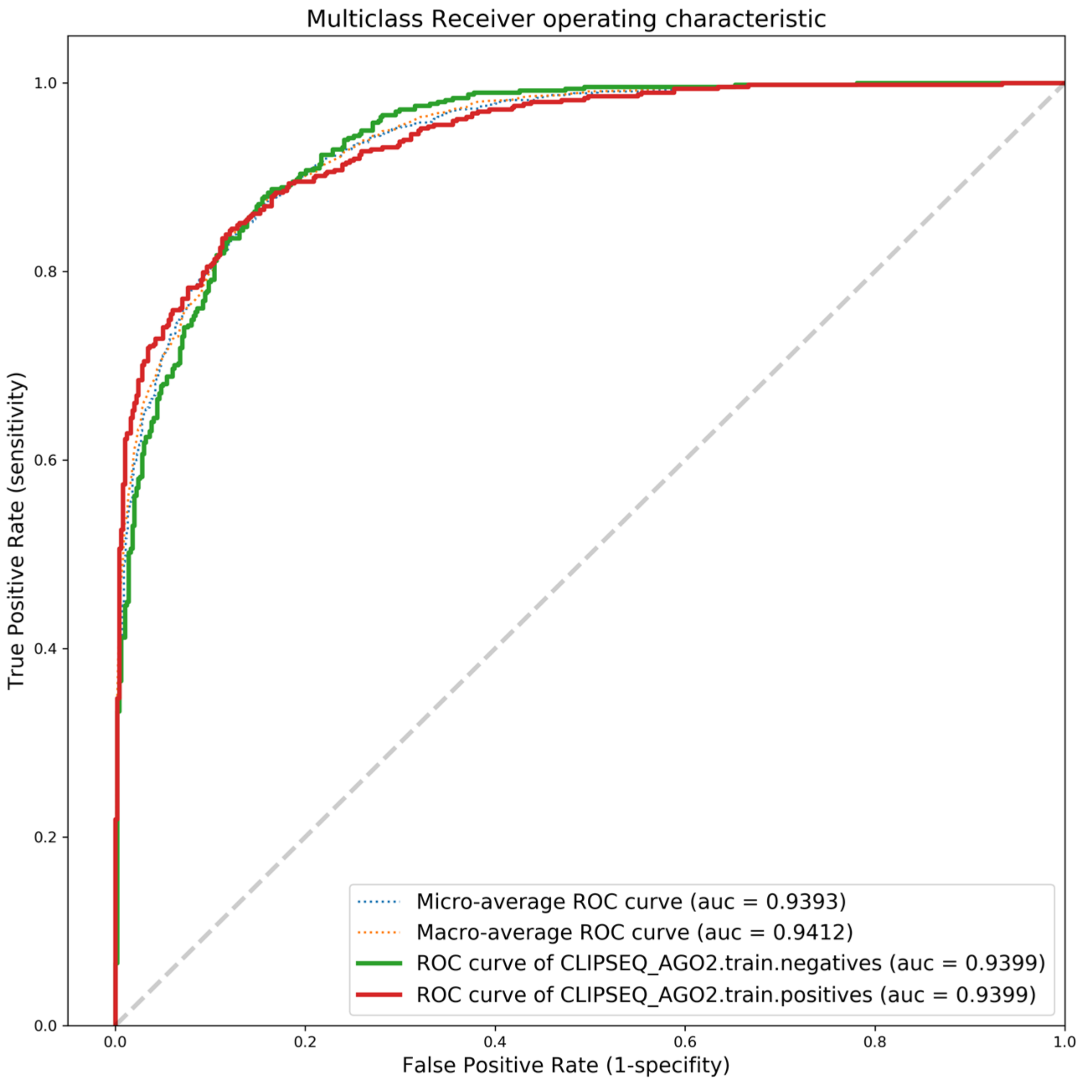

Fig.2 Ago1-4 ROC curve

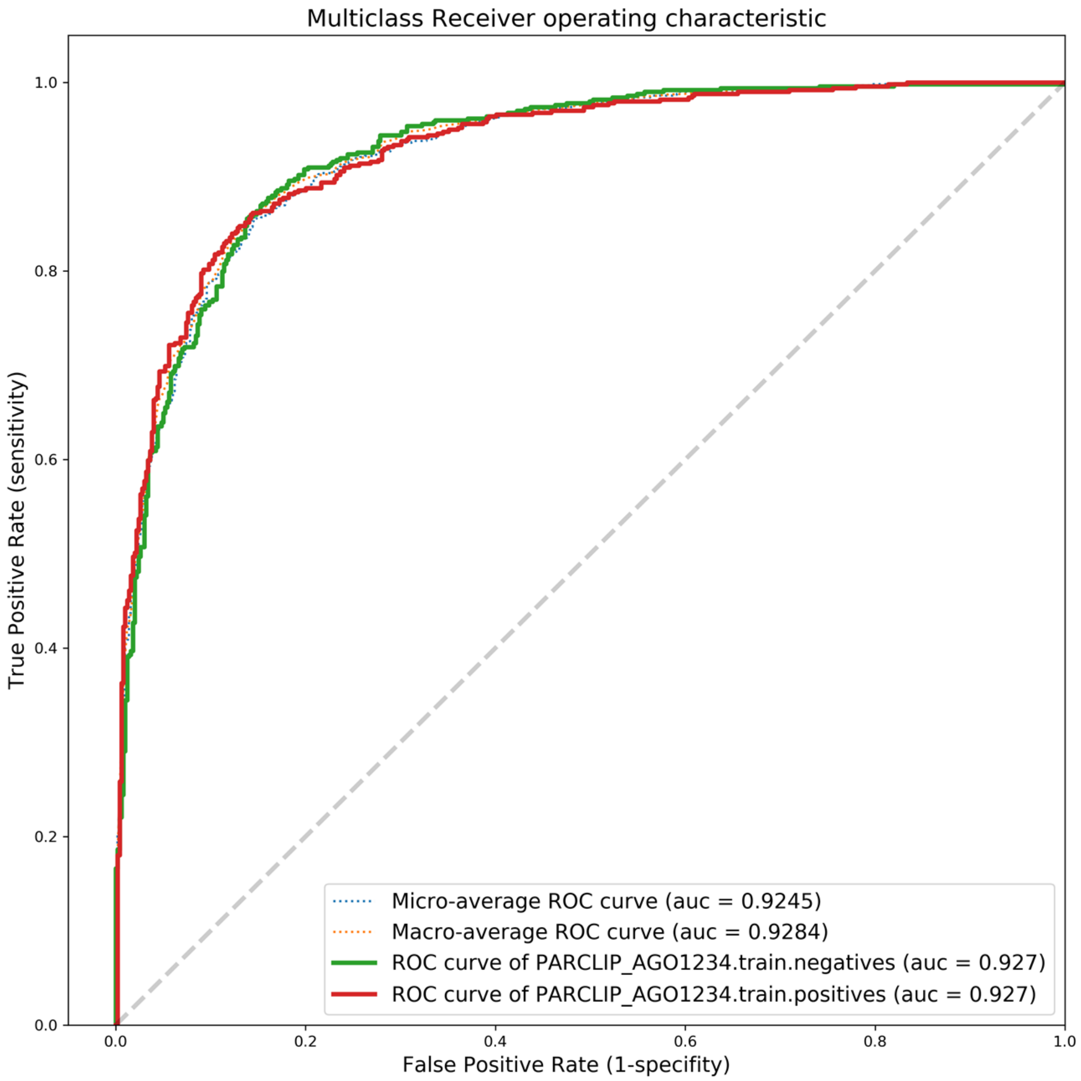

Fig.3 ALKBH5 ROC curve

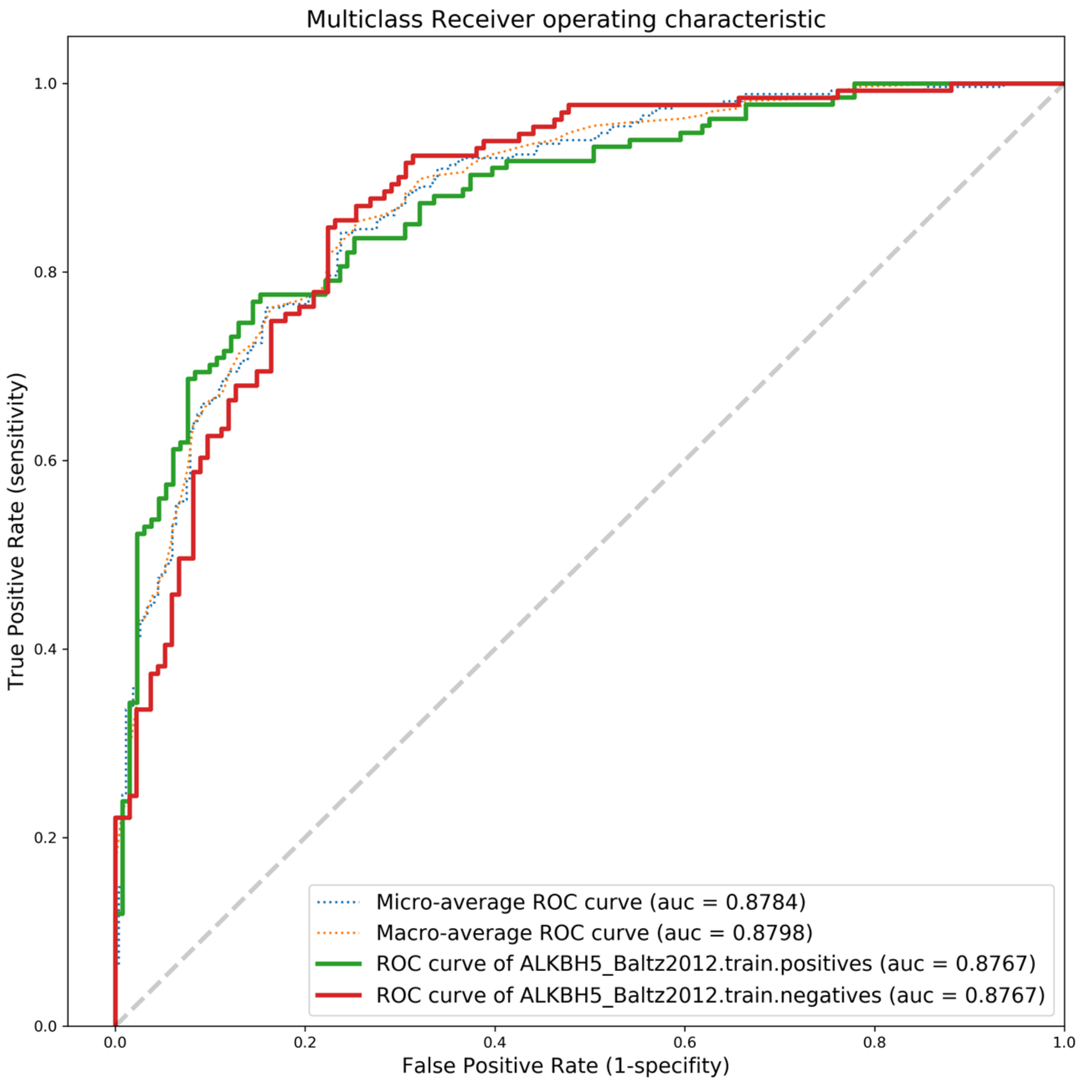

Fig.4 C17ORF85 ROC curve

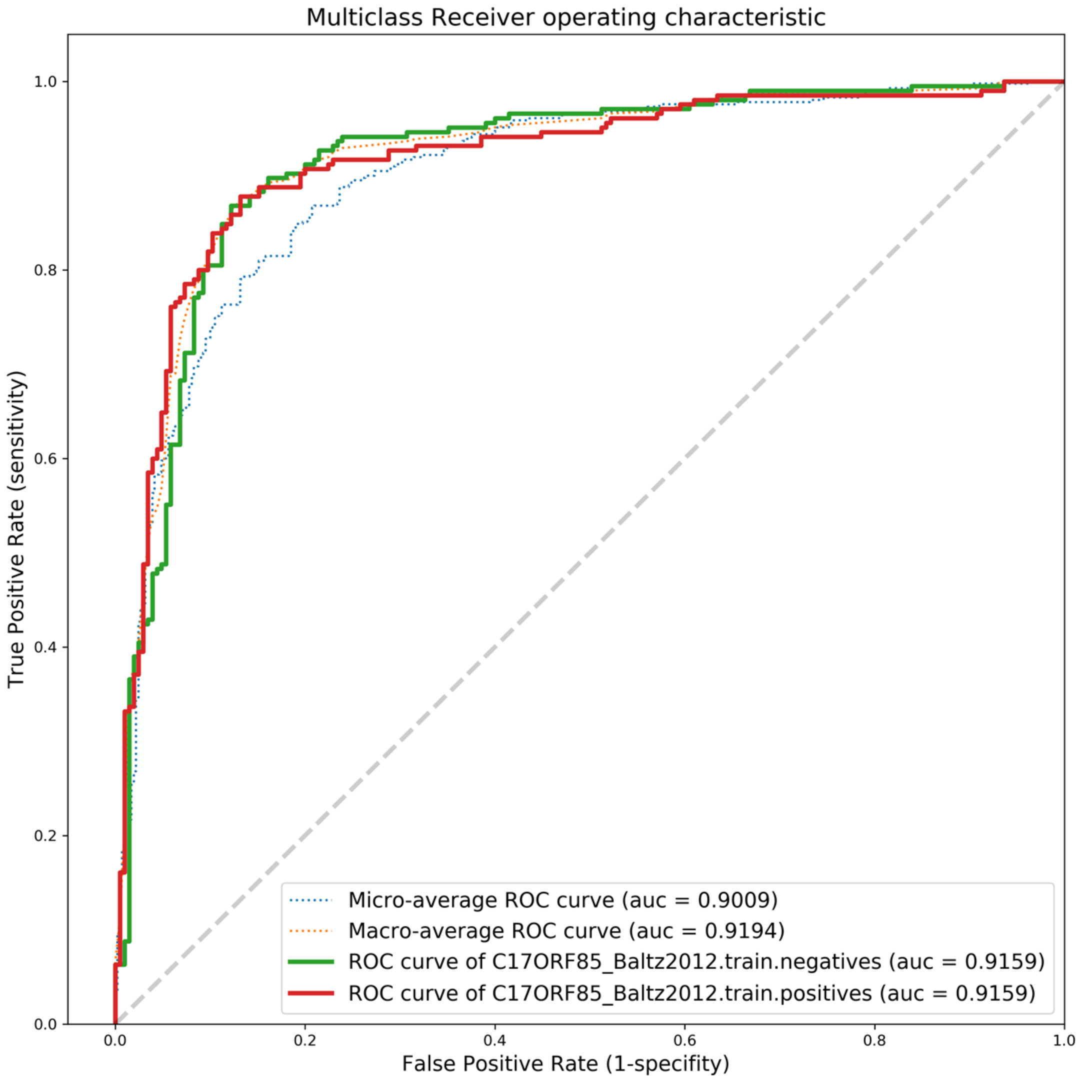

Fig.5 C22ORF28 ROC curve

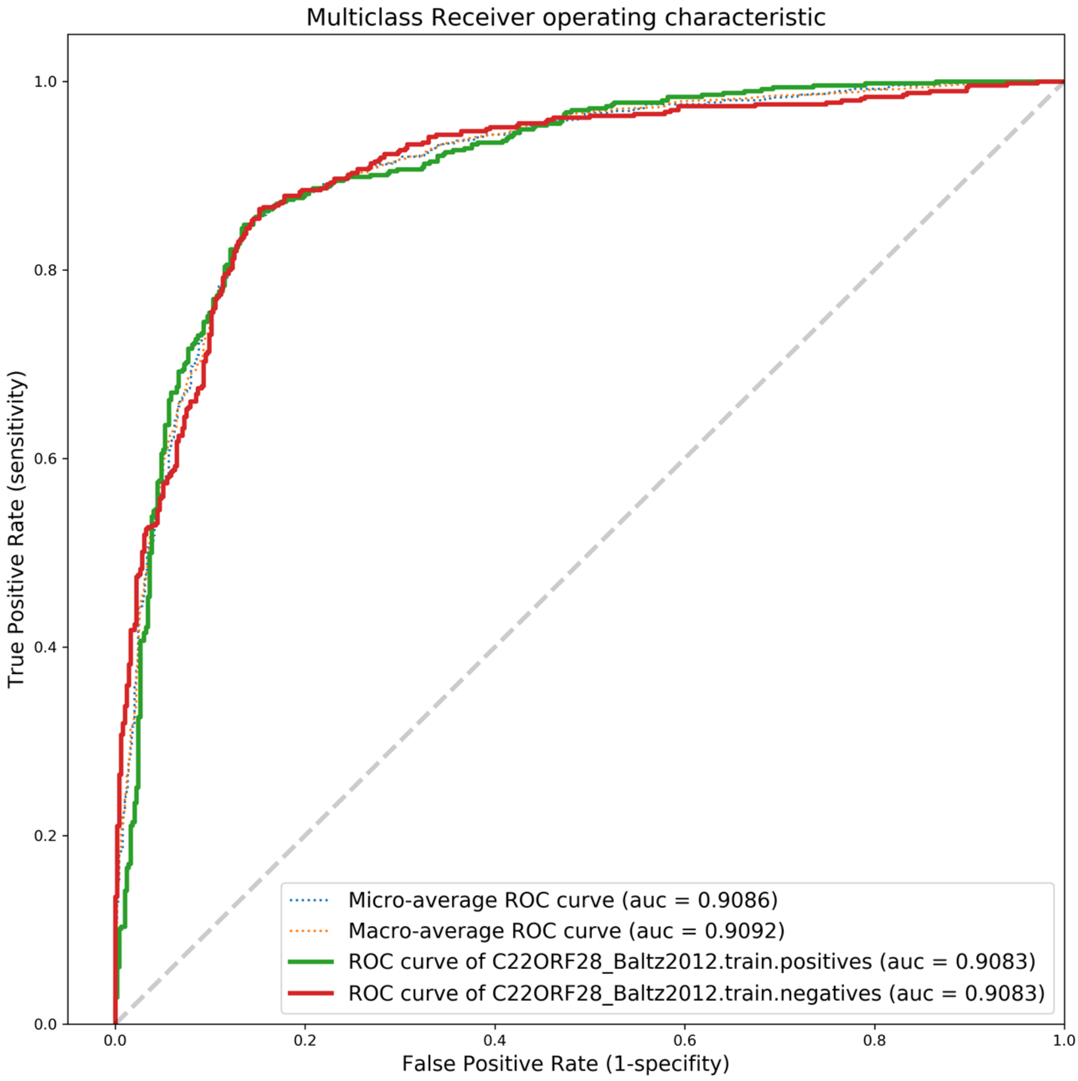

Fig.6 CAPRIN1 ROC curve

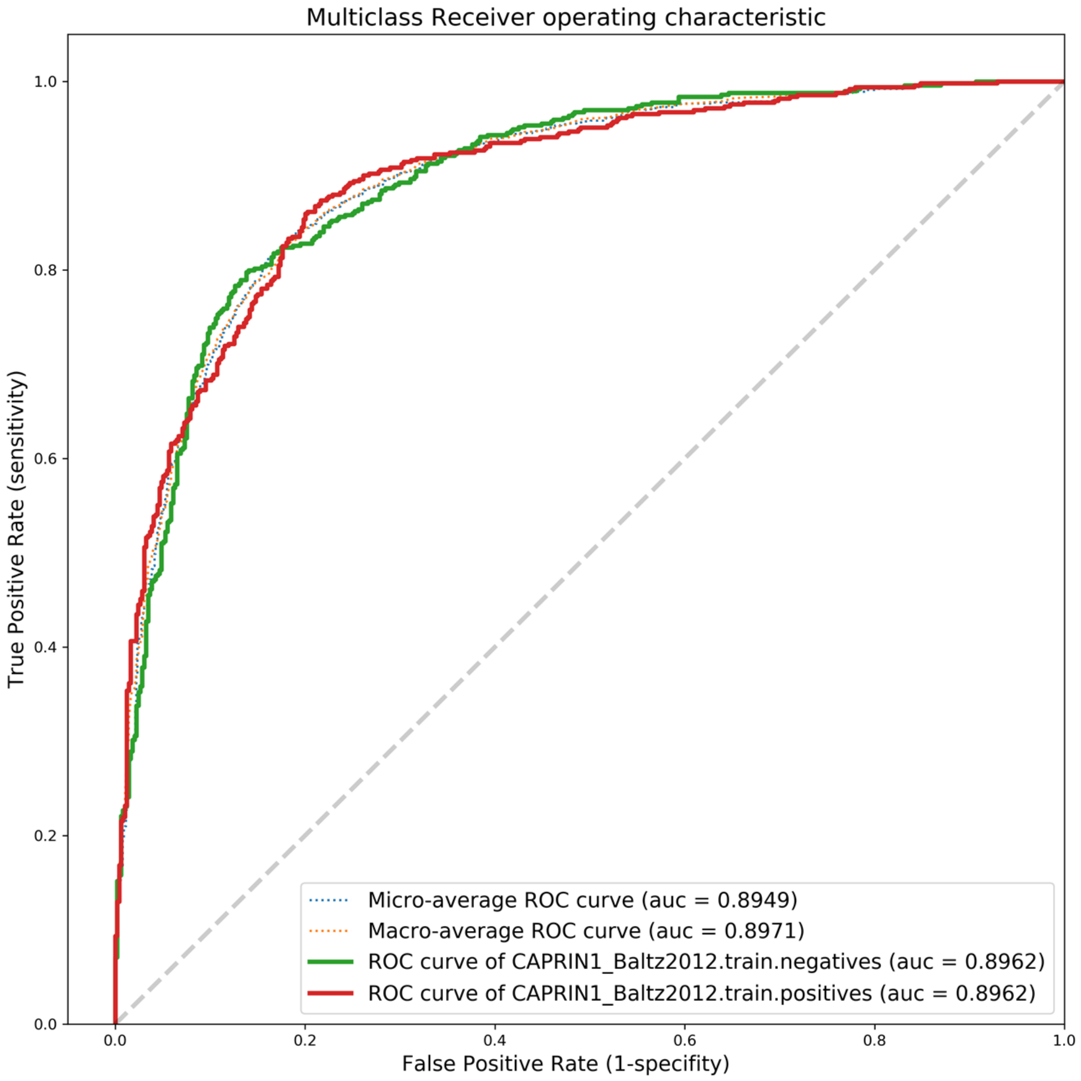

Fig.7 ELAVL1 (CLIP-Seq) ROC curve

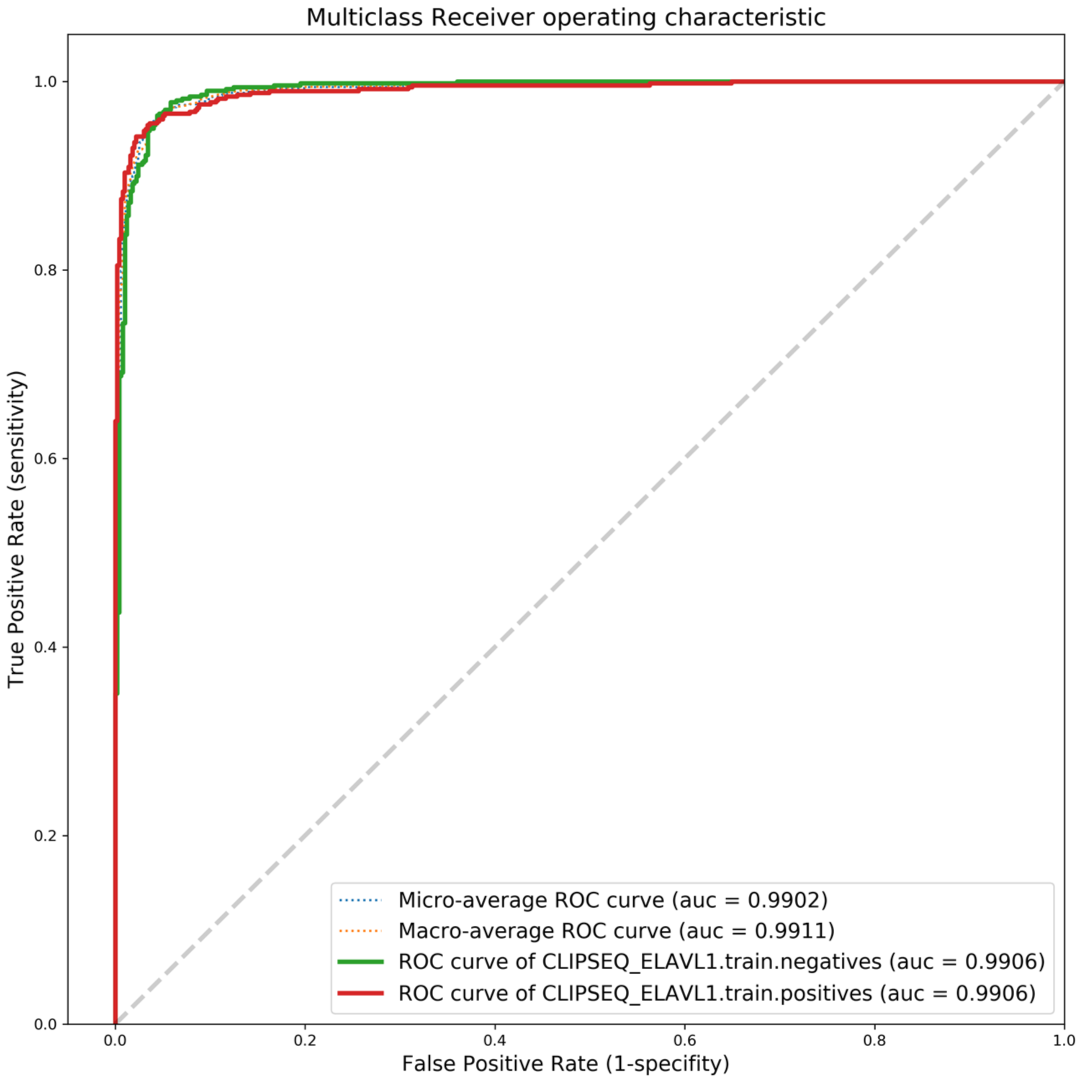

Fig.8 EWSR1 ROC curve

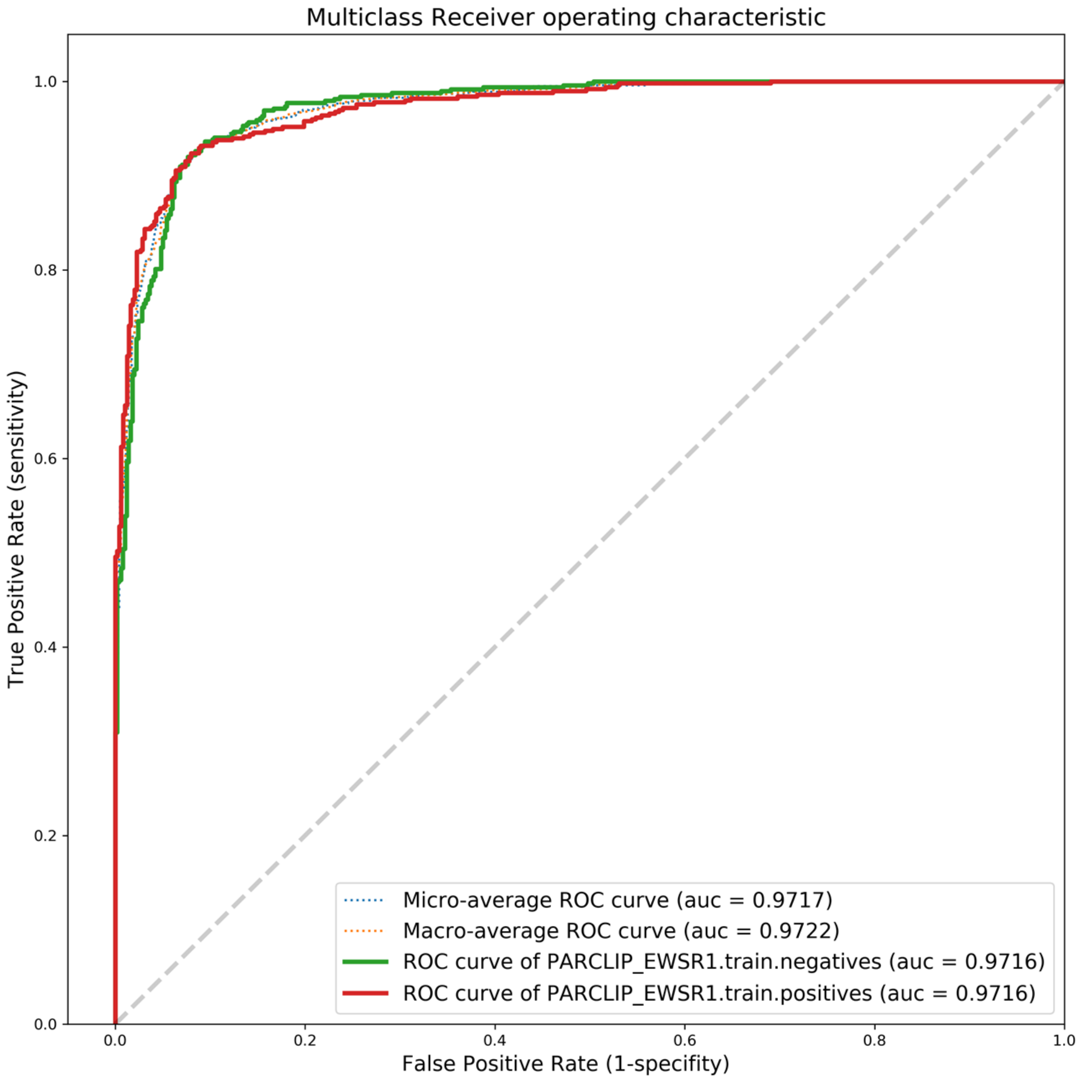

Fig.9 FUS ROC curve

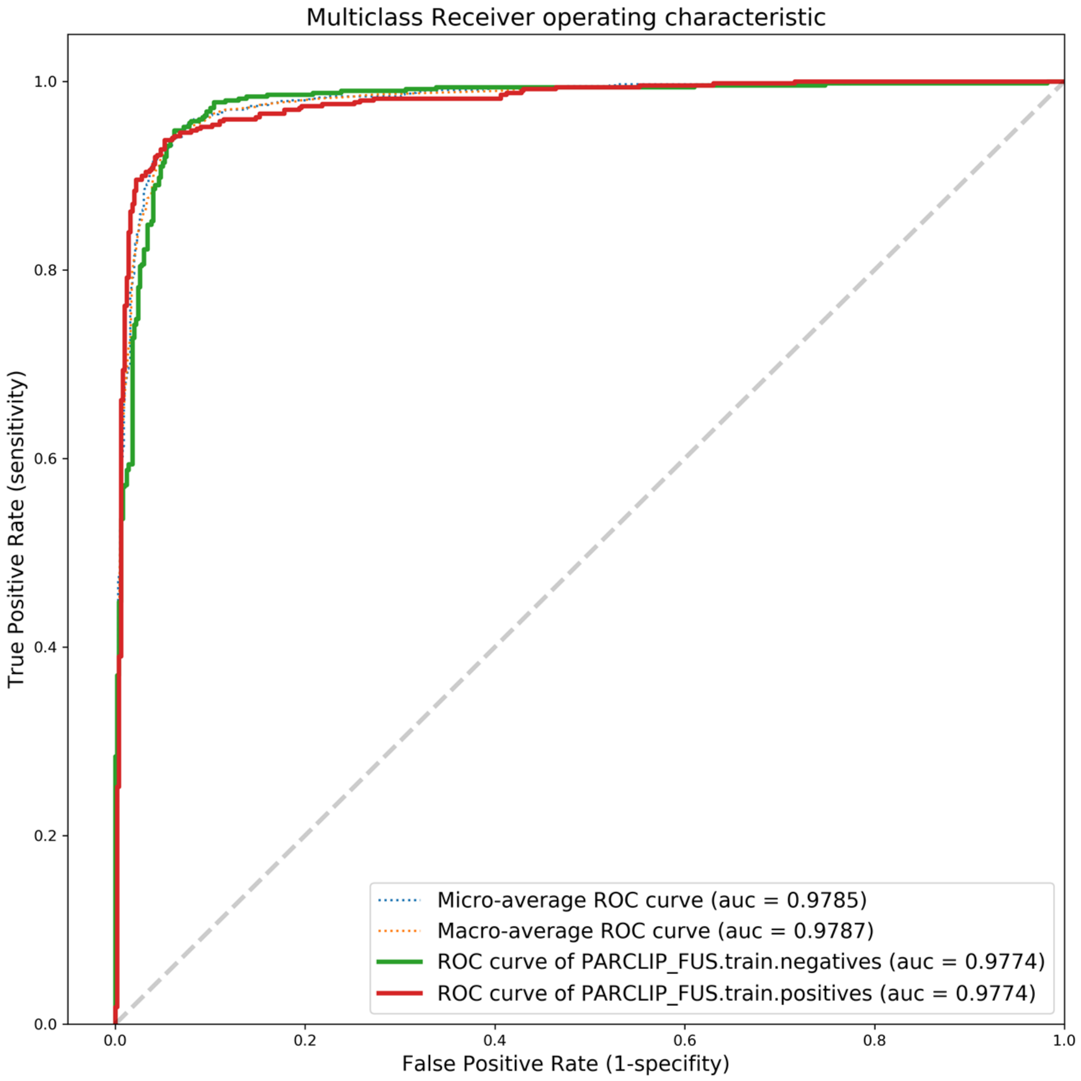

Fig.10 hnRNPC ROC curve

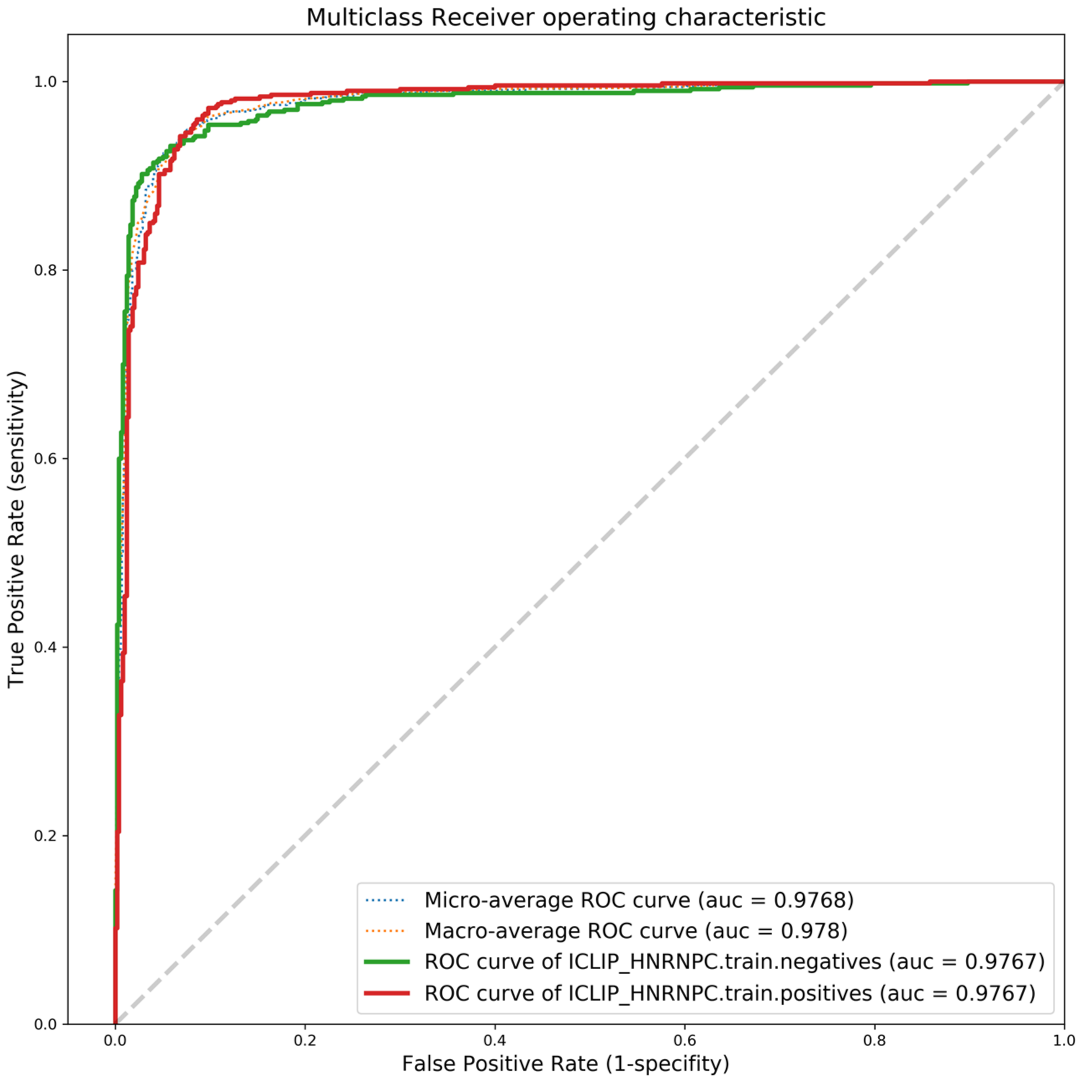

Fig.11 HUR ROC curve

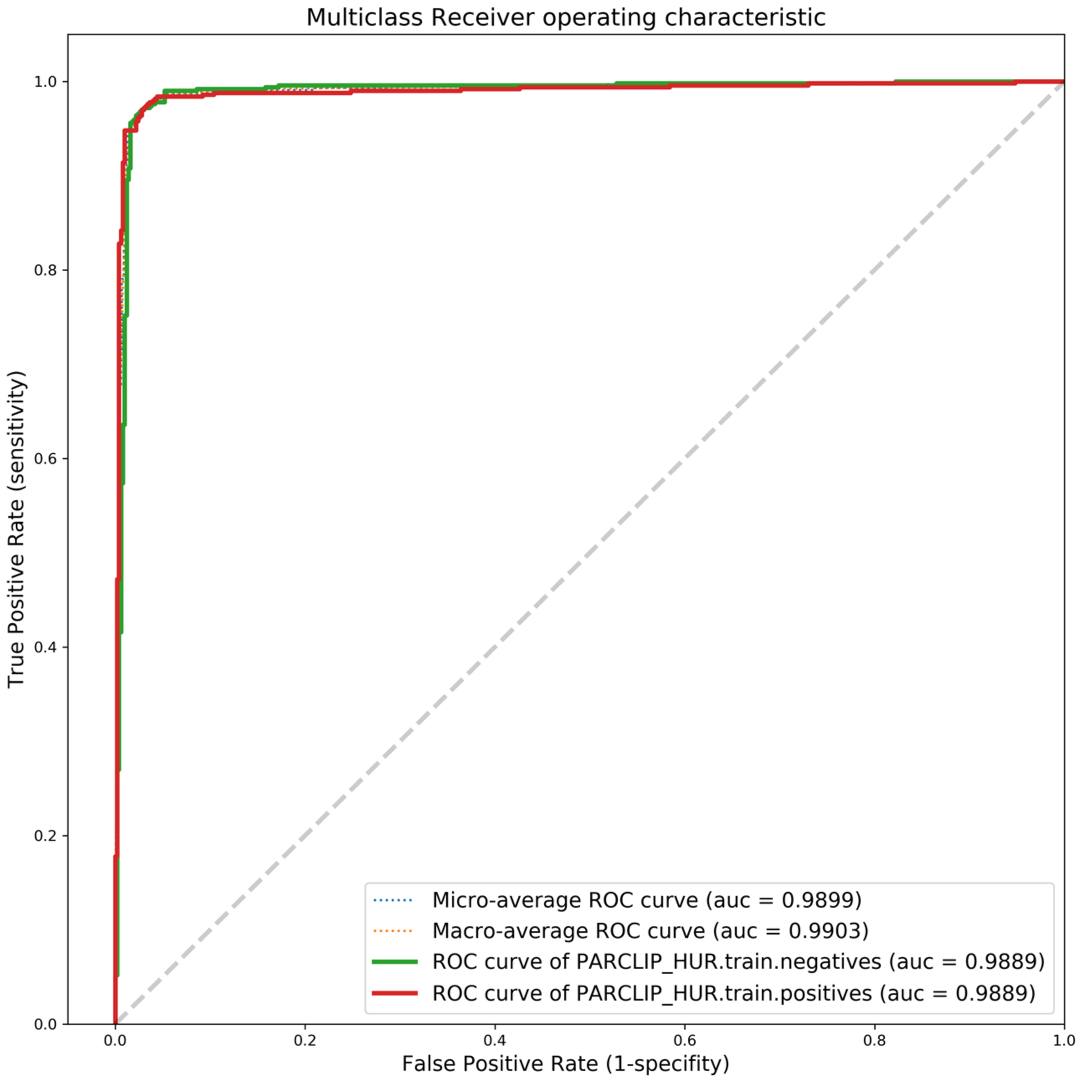

Fig.12 IGF2BP123 ROC curve

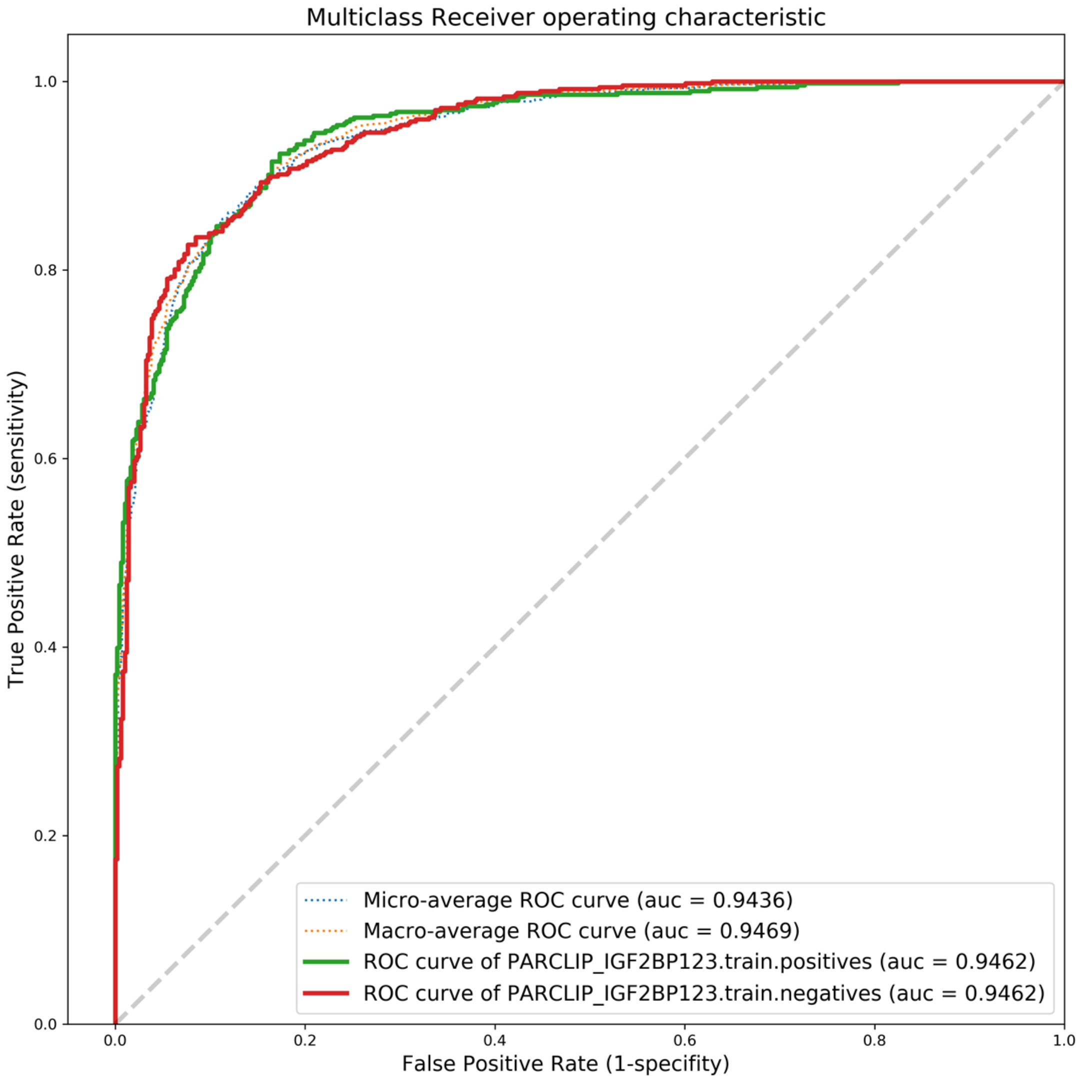

Fig.13 MOV10 ROC curve

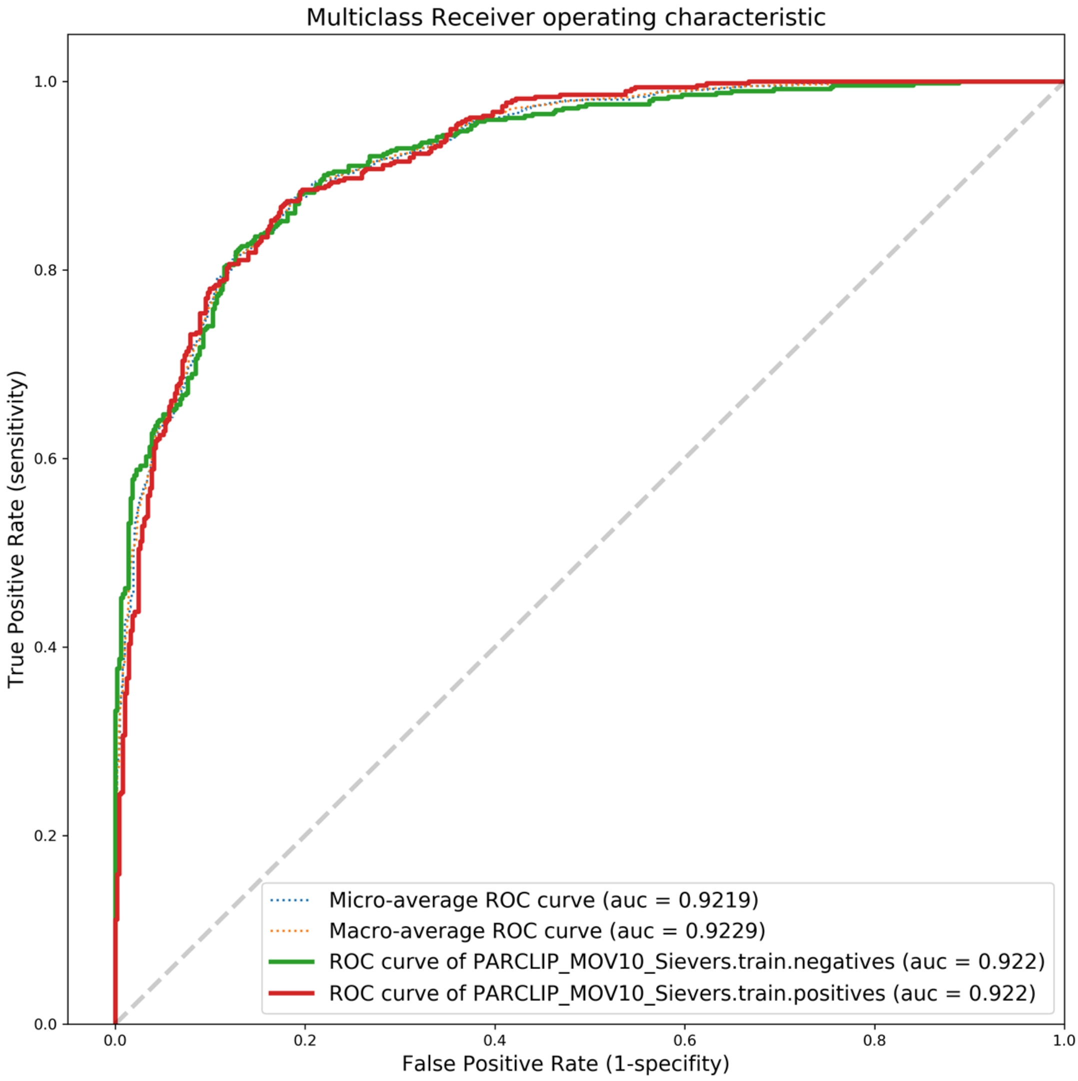

Fig.14 ELAVL1 (PARCLIP) ROC curve

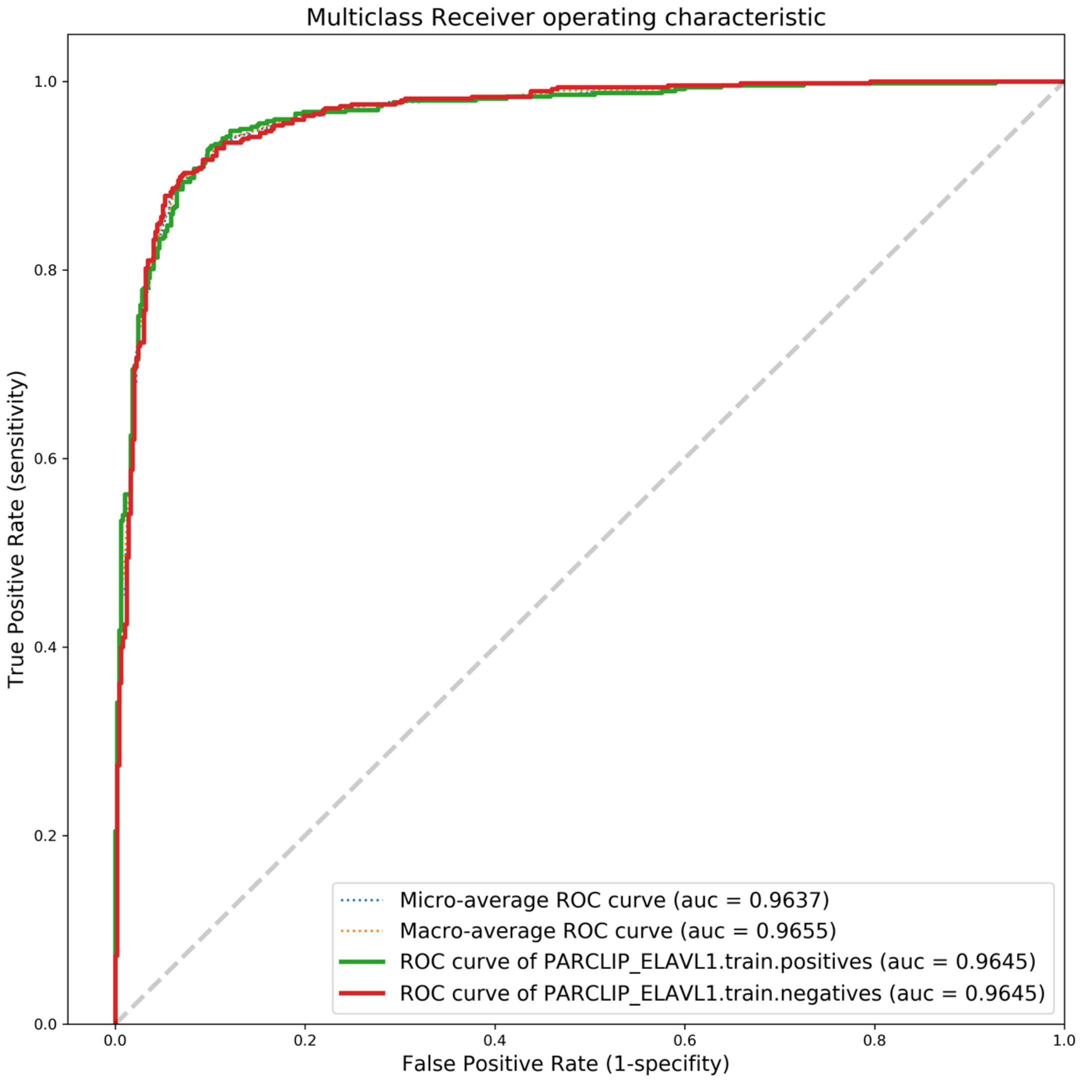

Fig.15 ELAVL1A (PARCLIP)

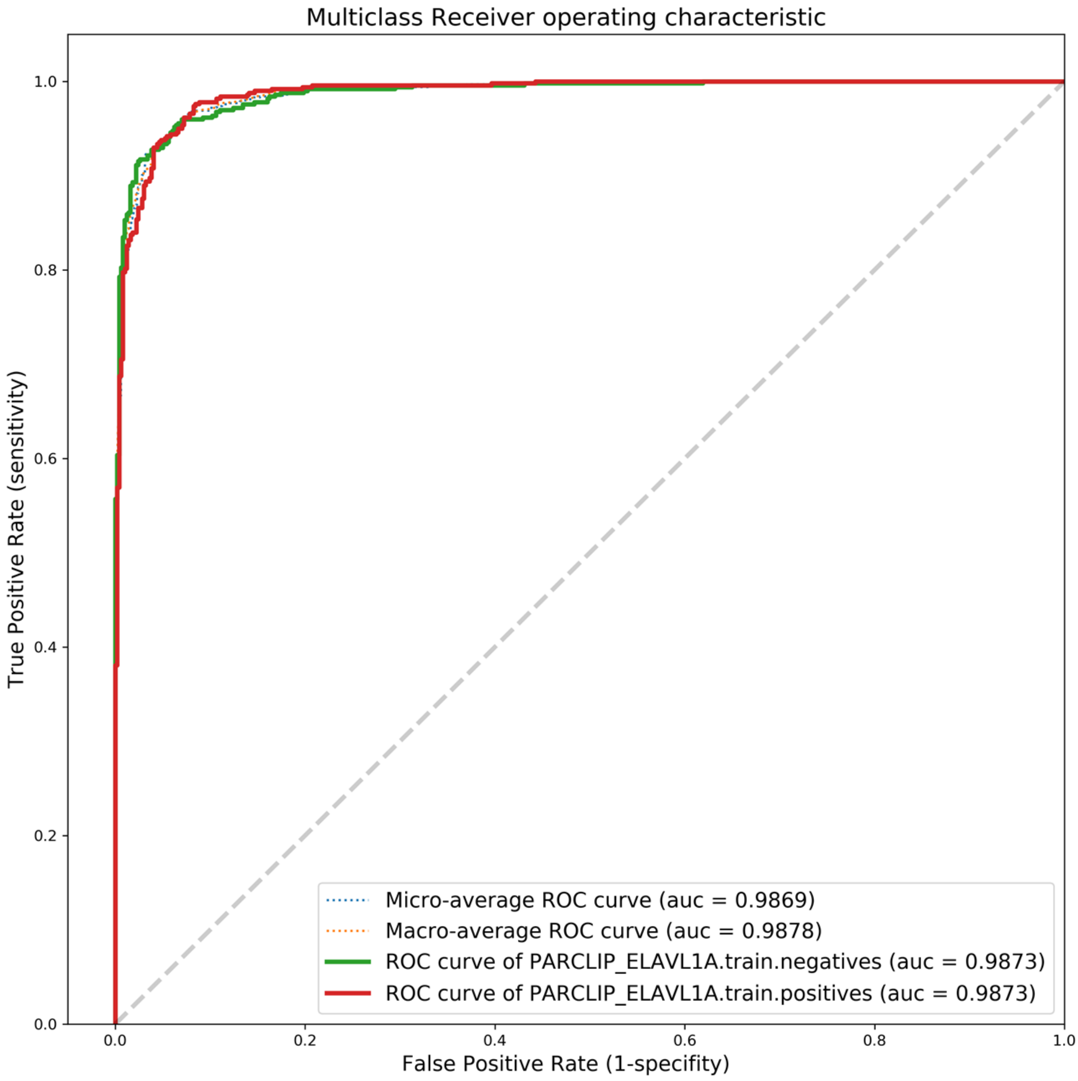

Fig.16 PTB ROC curve

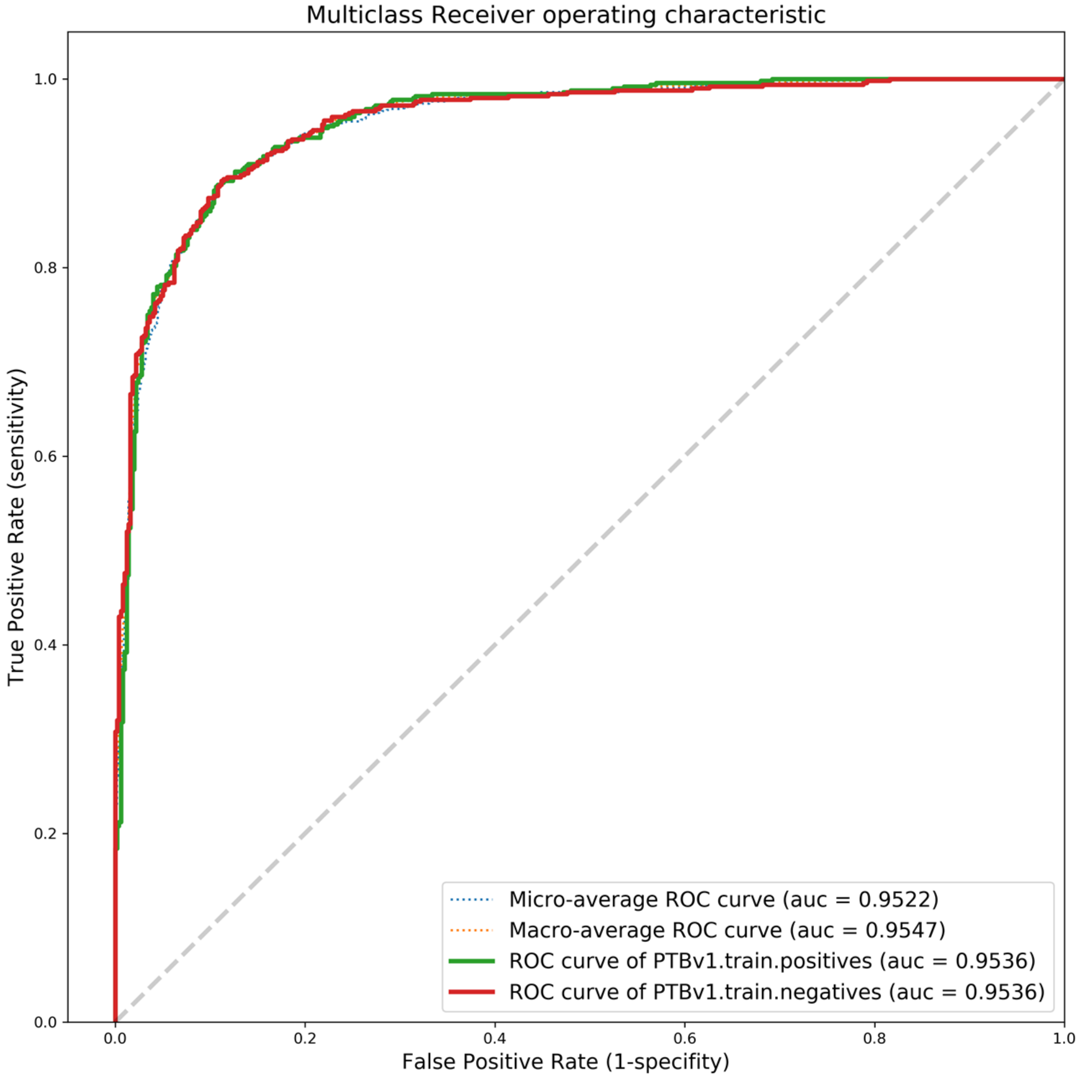

Fig.17 PUM2 ROC curve

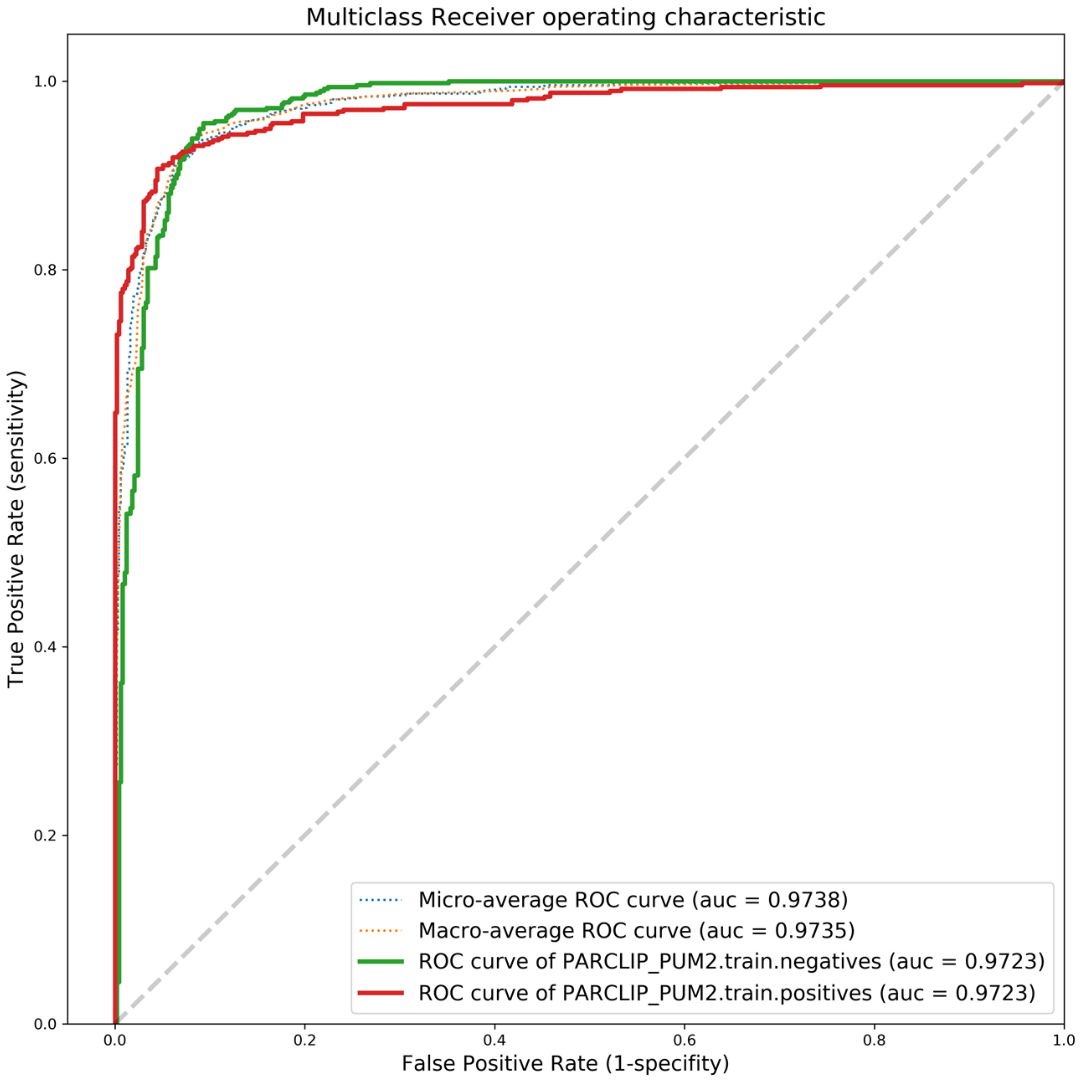

Fig.18 QKI ROC curve

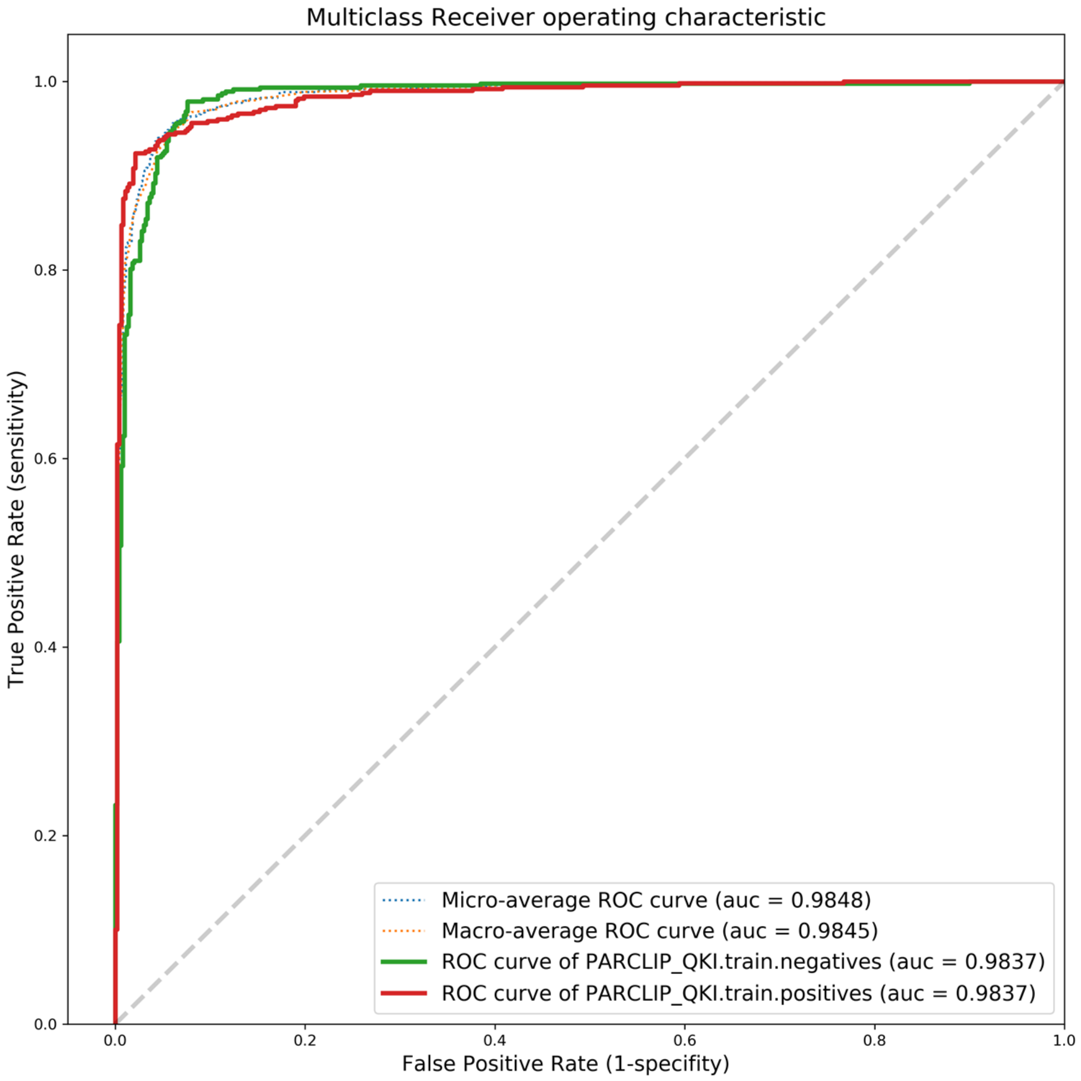

Fig.19 SFRS1 ROC curve

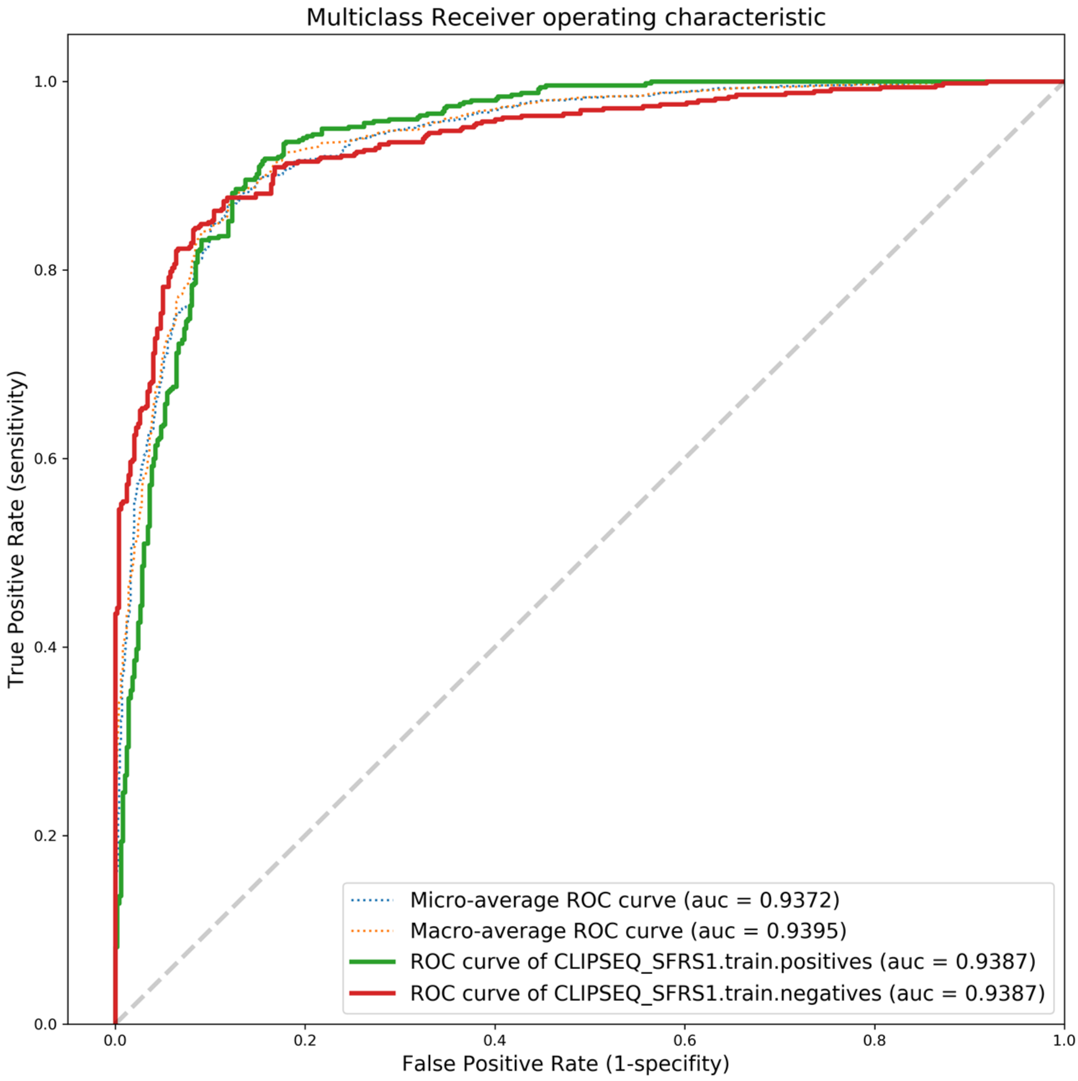

Fig.20 TAF15 ROC curve

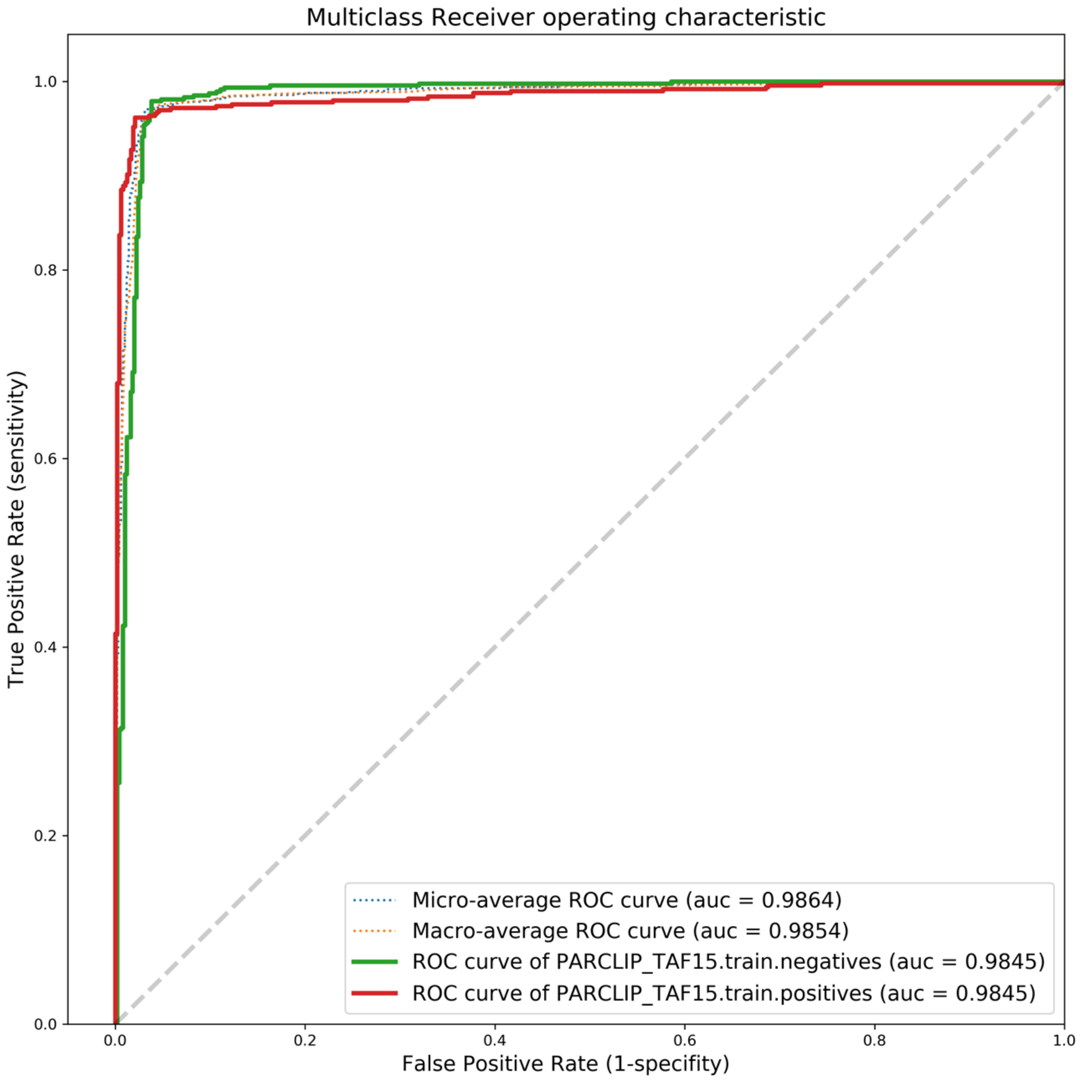

Fig.21 TDP43 ROC curve

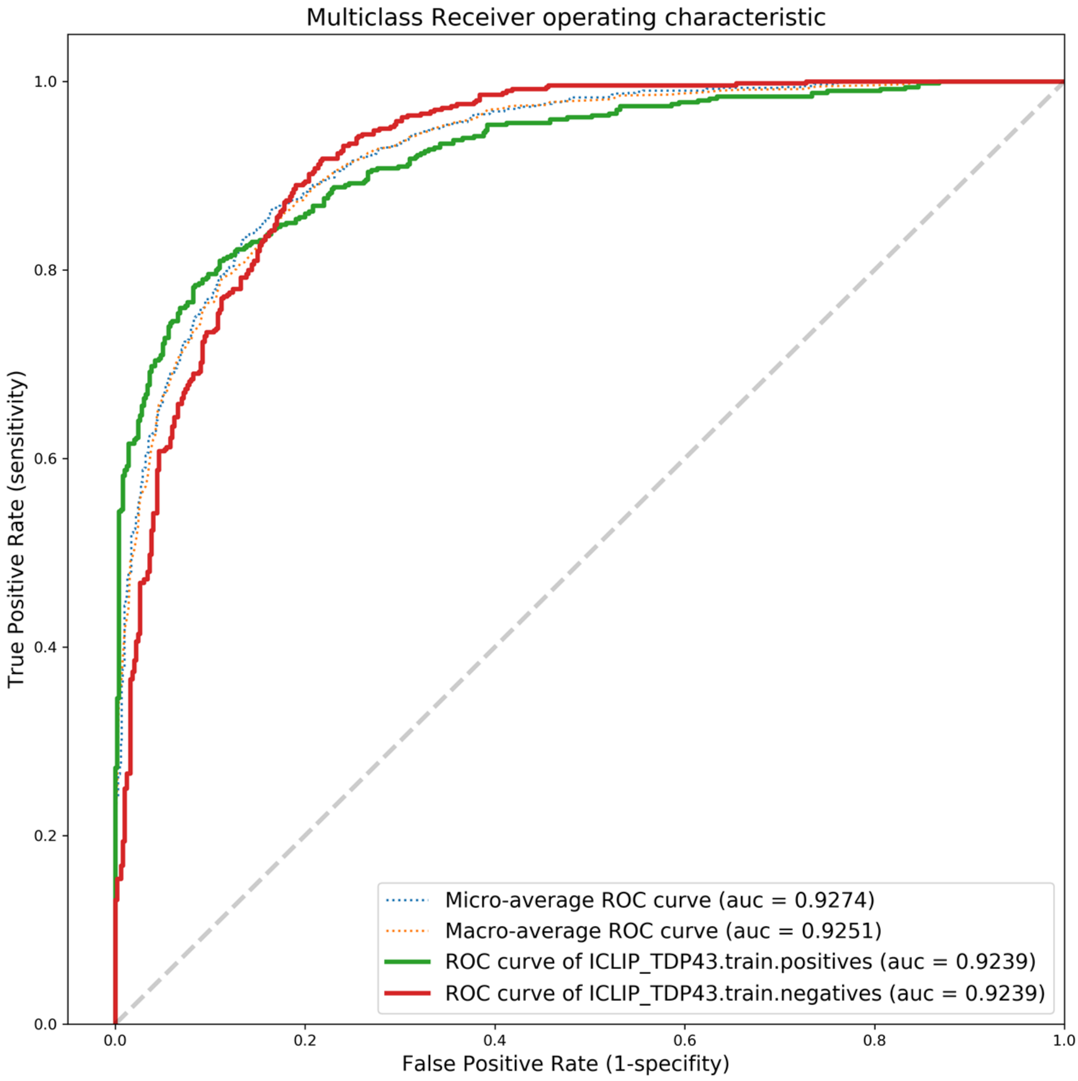

Fig.22 TIA1 ROC curve

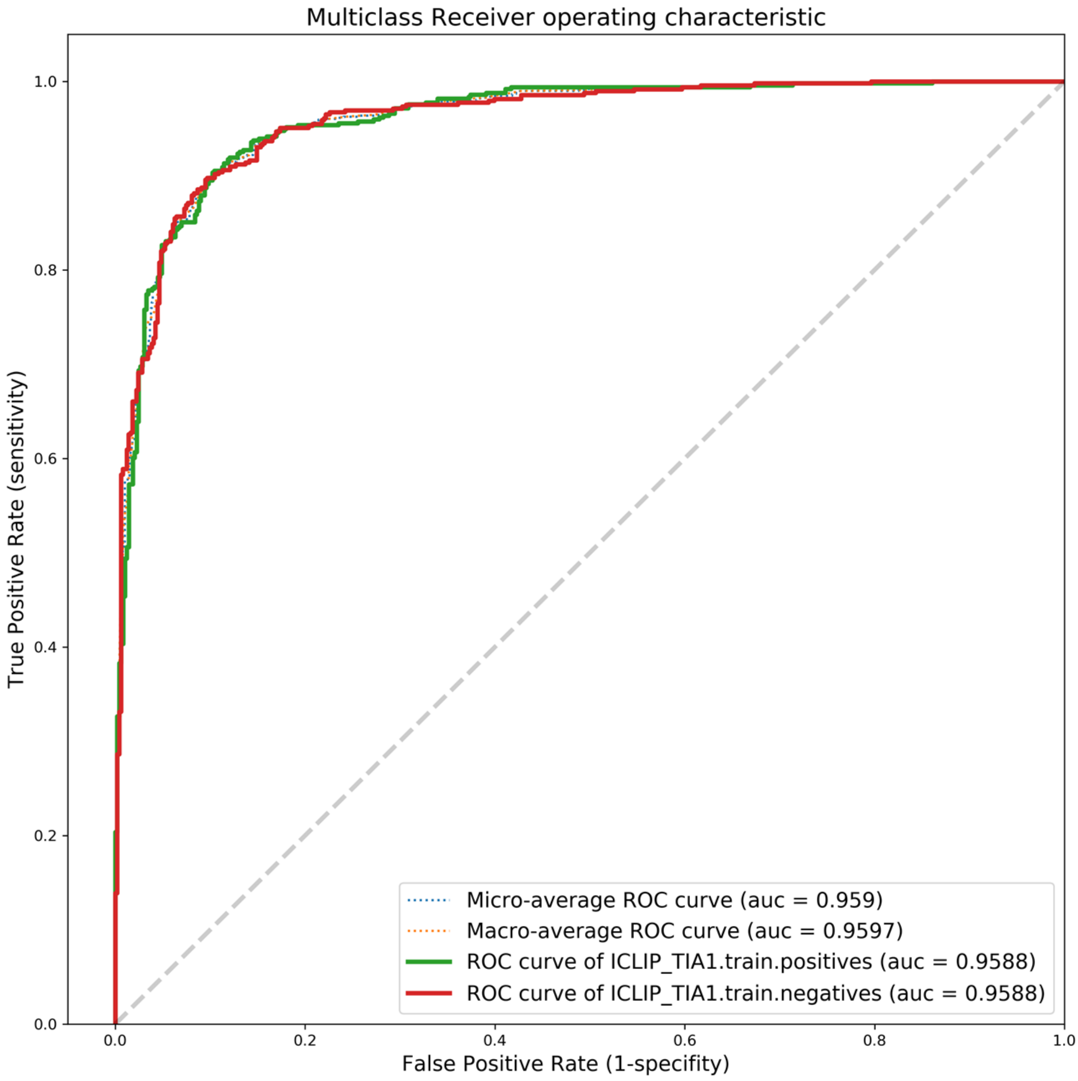

Fig.23 TIAL1 ROC curve

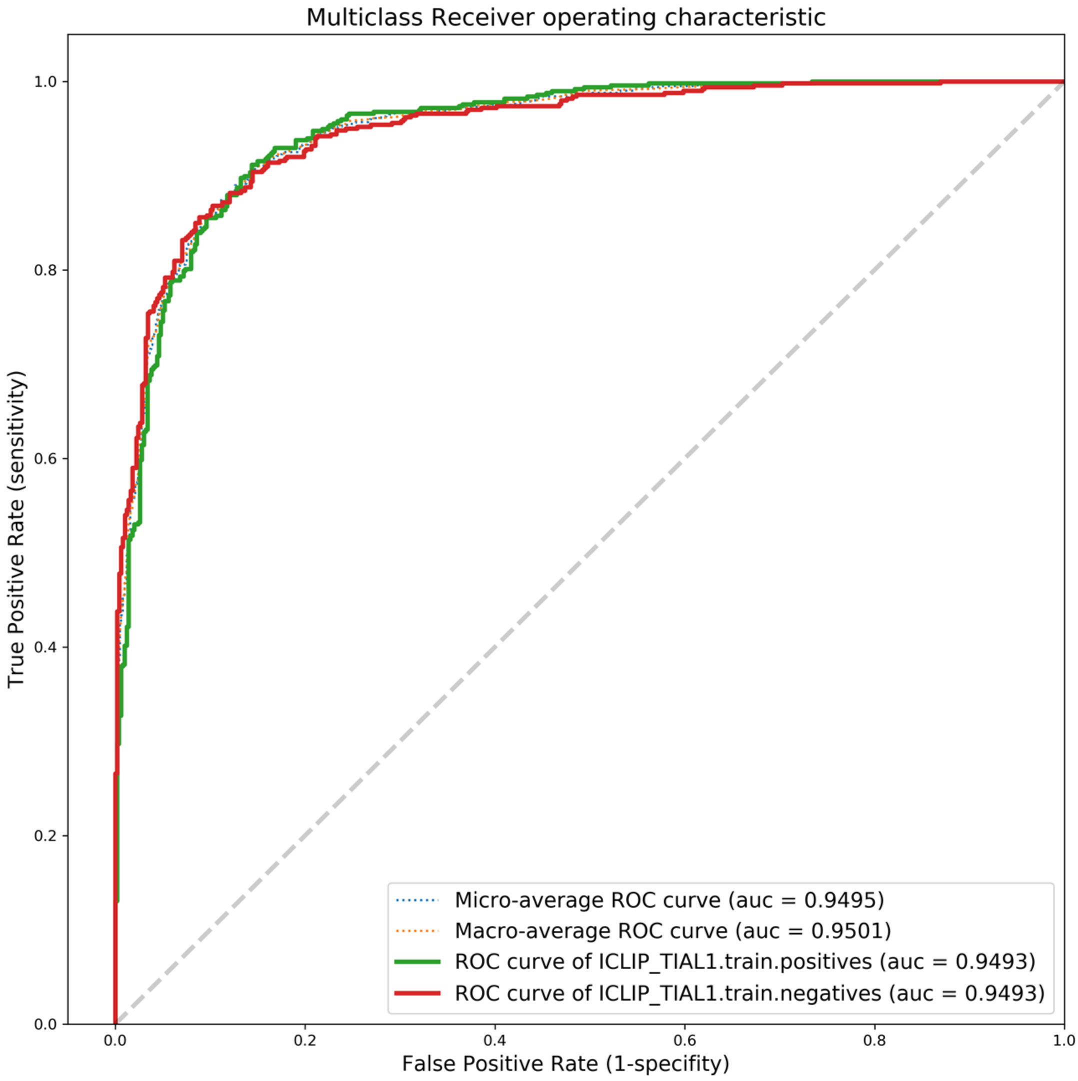

Fig.24 ZC3H7B ROC curve

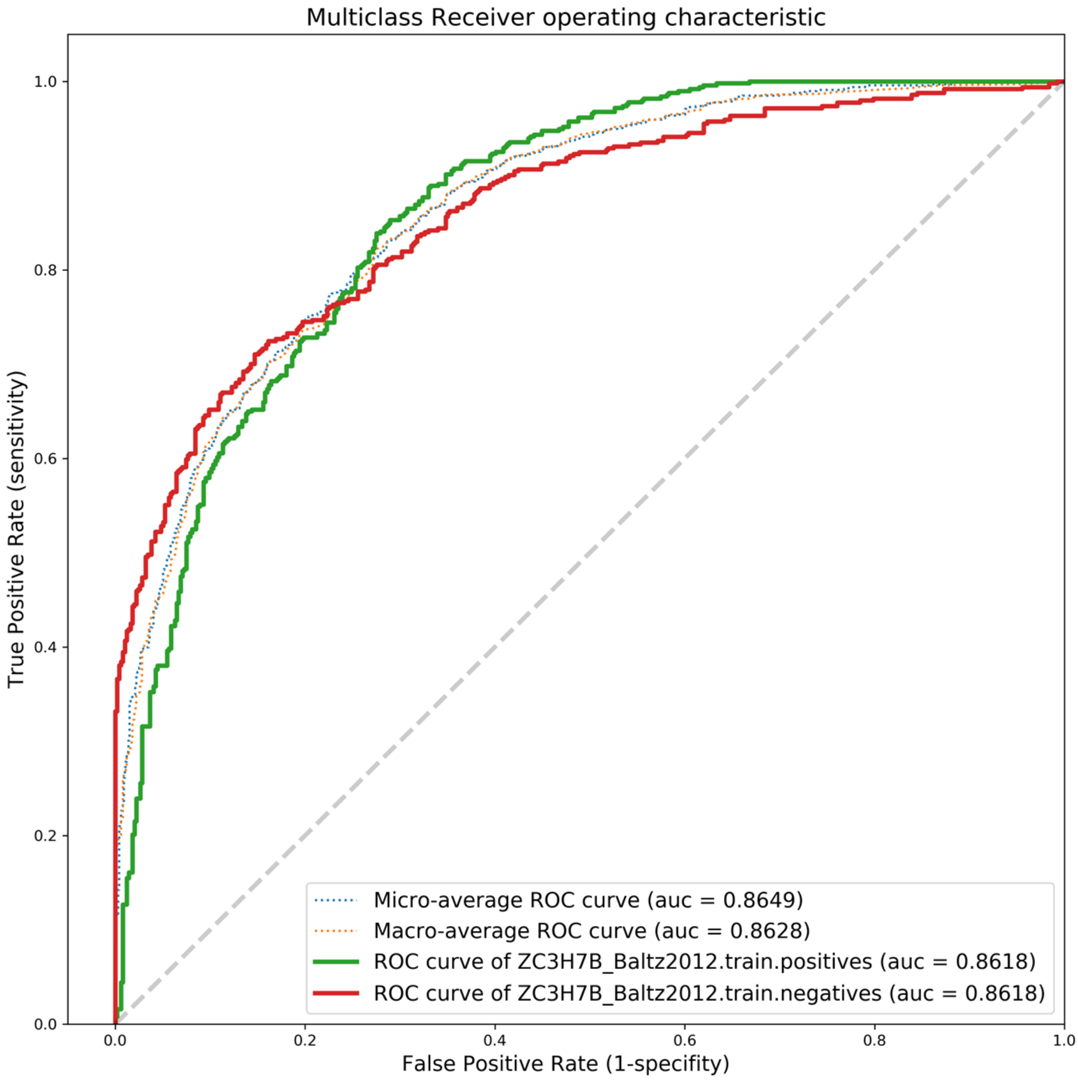
