## Additional file 1 for "ENNGene: an Easy Neural Network model building tool for Genomics": enngene.py

```
import base64
from datetime import datetime
import numpy as np
import streamlit as st  # TODO outside try-except, add check if installed
import logging
import os
import random as py_rand

import tempfile
import tensorflow as tf

from lib.utils.exceptions import MyException

try:
    logger = logging.getLogger('root')
    formatter = logging.Formatter('%(asctime)s - %(name)s - %(levelname)s - %(message)s',  datefmt='%m/%d/%Y %I:%M:%S %p')

    if any(type(handler) == logging.FileHandler for handler in logger.handlers):
        file_handler = [handler for handler in logger.handlers if type(handler) == logging.FileHandler][0]
        if not os.path.isfile(file_handler.baseFilename):
            logger.removeHandler(file_handler)

    if not any(type(handler) == logging.FileHandler for handler in logger.handlers):
        logfile_path = os.path.join(tempfile.gettempdir(), f'{datetime.now().strftime("%Y-%m-%d_%H:%M:%S")}_app.log')
        file_handler = logging.FileHandler(logfile_path, mode='a')
        file_handler.setLevel(logging.DEBUG)
        file_handler.setFormatter(formatter)
        logger.addHandler(file_handler)

    if not any(type(handler) == logging.StreamHandler for handler in logger.handlers):
        console_handler = logging.StreamHandler()
        console_handler.setLevel(logging.DEBUG)
        console_handler.setFormatter(formatter)
        logger.addHandler(console_handler)

except Exception as err:
    st.warning(f'Failed to load logger, continuing without logging...')


def enngene():
    st.sidebar.title('ENNGene')
    st.sidebar.markdown('')

    available_subcommands = {'Preprocessing': 'preprocess',
                             'Training': 'train',
                             'Evaluation': 'evaluate',
                             'Prediction': 'predict'}

    subcommand = available_subcommands[st.sidebar.selectbox(
        'Select a task to be run:',
        list(available_subcommands.keys())
    )]

    st.sidebar.markdown('')
    st.sidebar.markdown('[Documentation](https://github.com/ML-Bioinfo-CEITEC/ENNGene/blob/master/README.md)')
    st.sidebar.markdown('[FAQ](https://github.com/ML-Bioinfo-CEITEC/ENNGene/blob/master/FAQ.md)')
    st.sidebar.markdown('[GitHub](https://github.com/ML-Bioinfo-CEITEC/ENNGene)')

    st.sidebar.markdown('---')
    encoded_ceitec_logo = base64.b64encode(open(os.path.join(os.path.dirname(os.path.abspath(__file__)), 'imgs/CEITEC_logo_K-0.png'), "rb").read()).decode()
    encoded_muni_logo = base64.b64encode(open(os.path.join(os.path.dirname(os.path.abspath(__file__)), 'imgs/muni-lg-eng-rgb.png'), "rb").read()).decode()
    html_cl = f'<a href="https://www.ceitec.eu/" target="_blank"><img src="data:image/png;base64, {encoded_ceitec_logo}" width="100%" /></a>'
    html_ml = f'<a href="https://www.muni.cz/en" target="_blank"><img src="data:image/png;base64, {encoded_muni_logo}" width="100%" /></a>'
    st.sidebar.markdown(html_cl, unsafe_allow_html=True)
    st.sidebar.markdown(html_ml, unsafe_allow_html=True)
    st.sidebar.markdown('---')

    logger.debug(f'ENNGene started with the following subcommand: {subcommand}')

    np.random.seed(89)
    py_rand.seed(123)
    tf.random.set_seed(456)

    module_path = f'lib.{subcommand}.{subcommand}'
    subcommand_class = ''.join(x.title() for x in subcommand.split('_'))
    module = __import__(module_path, fromlist=[subcommand_class])
    # use dispatch pattern to invoke object of class with same name as the subcommand
    subcommand = getattr(module, subcommand_class)
    subcommand()


if __name__ == '__main__':
    try:
        enngene()
    except MyException as err:
        logger.exception(f'{err.__class__.__name__}: {err}')
        st.warning(f'{err}\n Exiting...')
    except Exception as err:
        logger.exception(f'{err.__class__.__name__}: {err}')
        file_handler = [handler for handler in logger.handlers if type(handler) == logging.FileHandler]
        if file_handler:
            logfile_path = file_handler[0].baseFilename
        st.warning(
            f'Unexpected error occurred in the application. Exiting... \n For more details check the log file at {logfile_path}.')
```
