## Additional file 1 for "ENNGene: an Easy Neural Network model building tool for Genomics": evaluate.py

```
import datetime
import logging
import numpy as np
import os
import streamlit as st
import tensorflow as tf
import yaml

from ..utils.dataset import Dataset
from ..utils import sequence as seq
from ..utils.subcommand import Subcommand
from ..utils.exceptions import UserInputError

logger = logging.getLogger('root')


class Evaluate(Subcommand):
    SEQ_TYPES = {'BED file': 'bed',
                 'FASTA file': 'fasta',
                 'Blackbox dataset': 'blackbox'}
    
    def __init__(self):
        # test module - either on already mapped blackbox dataset, or not encoded dataset, eg from different experiment to check transferbality of the results

        self.params = {'task': 'Evaluate'}
        self.validation_hash = {'is_model_file': [],
                                'is_bed': [],
                                'is_blackbox': [],
                                'is_fasta': [],
                                'is_wig_dir': []}
        self.params['model_folder'] = None

        st.markdown('# Evaluation')
        st.markdown('')
        self.general_options()

        st.markdown('## Model')
        self.model_options()

        # TODO add option to use already prepared file ? (sequences.tsv)
        st.markdown('## Sequences')
        self.sequence_options(self.SEQ_TYPES, evaluation=True)

        st.markdown('')
        self.params['ig'] = st.checkbox('Calculate Integrated Gradients', self.defaults['ig'])
        if self.params['ig']:
            self.params['smoothgrad'] = st.checkbox('Apply smoothgrad method', self.defaults['smoothgrad'])
            st.markdown('###### **WARNING**: Calculating the integrated gradients is a time-consuming process, '
                        'it may take several minutes up to few hours (depending on the number of sequences). ' +
                        'Smoothgrad increases time complexity of integrated gradients by a factor of 20.')


        self.validate_and_run(self.validation_hash)

    def run(self):
        status = st.empty()
        status.text('Preparing sequences...')

        if self.previous_param_file:
            with open(self.previous_param_file, 'r') as file:
                previous_params = yaml.safe_load(file)
                klasses = previous_params['Preprocess']['klasses']
                klass_alphabet = {klass: i for i, klass in enumerate(klasses)}
                encoded_labels = seq.onehot_encode_alphabet(klass_alphabet)
        else:
            raise UserInputError('Could not read class labels from parameters.yaml file).')

        self.params['eval_dir'] = os.path.join(self.params['output_folder'], 'evaluation',
                                               f'{str(datetime.datetime.now().strftime("%Y%m%d-%H%M"))}')
        self.ensure_dir(self.params['eval_dir'])

        prepared_file_path = os.path.join(self.params['eval_dir'], 'sequences.tsv')

        if self.params['seq_type'] == 'bed' or self.params['seq_type'] == 'fasta':
            if self.params['seq_type'] == 'bed':
                dataset = Dataset(bed_file=self.params['seq_source'], branches=self.params['branches'], category='eval',
                                  win=self.params['win'], win_place=self.params['win_place'])
                status.text(f"Mapping intervals to {len(self.params['branches'])} branch(es) and exporting...")
                dataset.sort_datapoints()
            elif self.params['seq_type'] == 'fasta':
                dataset = Dataset(fasta_file=self.params['seq_source'], branches=self.params['branches'], category='eval',
                                      win=self.params['win'], win_place=self.params['win_place'])
            dataset.map_to_branches(
                self.references, self.params['strand'], prepared_file_path, status, predict=True, ncpu=self.ncpu)
        elif self.params['seq_type'] == 'blackbox':
            dataset = Dataset.load_from_file(self.params['seq_source'])
            pass

        eval_x = dataset.encode_branches(dataset, self.params['branches'])
        eval_y = dataset.labels(encoding=encoded_labels)

        status.text('Evaluating model...')
        model = tf.keras.models.load_model(self.params['model_file'])
        predicted = self.evaluate_model(encoded_labels, model, eval_x, eval_y, self.params, self.params['eval_dir'])

        for i, klass in enumerate(self.params['klasses']):
            dataset.df[klass] = [y[i] for y in predicted]
        dataset.df['highest scoring class'] = self.get_klass(predicted, self.params['klasses'])

        placeholder = st.empty()
        if self.params['ig']:
            status.text('Calculating Integrated Gradients...')
            self.calculate_ig(dataset, model, eval_x, self.params['klasses'], self.params['branches'], self.params['smoothgrad'])

        placeholder.text('Exporting results...')
        result_file = os.path.join(self.params['eval_dir'], 'results.tsv')
        ignore = ['seq_encoded', 'fold_encoded', 'seq', 'fold', 'cons']
        dataset.save_to_file(ignore_cols=ignore, outfile_path=result_file)

        header = self.eval_header()
        row = self.eval_row(self.params)

        if self.previous_param_file:
            with open(self.previous_param_file, 'r') as file:
                previous_params = yaml.safe_load(file)
            if 'Train' in previous_params.keys():
                # Parameters missing in older versions of the code
                novel_params = {'auc': None, 'avg_precision': None}
                parameters = novel_params
                parameters.update(previous_params['Train'])
                header += f"{self.train_header()}"
                row += f"{self.train_row(parameters)}"
                if 'Preprocess' in previous_params.keys():
                    novel_params = {'win_place': 'rand'}  # It's always been 'random' for the previous versions
                    parameters = novel_params
                    parameters.update(previous_params['Preprocess'])
                    header += f'{self.preprocess_header()}\n'
                    row += f"{self.preprocess_row(parameters)}\n"
                else:
                    header += '\n'
                    row += '\n'
            else:
                header += '\n'
                row += '\n'

        self.finalize_run(logger, self.params['eval_dir'], self.params, header, row, placeholder, self.previous_param_file)
        status.text('Finished!')
        logger.info('Finished!')

    @staticmethod
    def default_params():
        return {
            'model_source': 'from_app',
            'model_folder': '',
            'model_file': '',
            'branches': [],
            'win': 100,
            'no_klasses': 2,
            'klasses': [],
            'seq_type': 'bed',
            'seq_source': '',
            'strand': True,
            'fasta_ref': '',
            'cons_dir': '',
            'win_place': 'center',
            'ig': True,
            'smoothgrad': False,
            'output_folder': os.path.join(os.path.expanduser('~'), 'enngene_output')
        }
```
