## Additional file 1 for "ENNGene: an Easy Neural Network model building tool for Genomics": predict.py

```
import datetime
import logging
import numpy as np
import os
import streamlit as st
import yaml
import tensorflow as tf


# TODO export the env when releasing, check pandas == 1.1.1
from ..utils.dataset import Dataset
from ..utils.subcommand import Subcommand

logger = logging.getLogger('root')


class Predict(Subcommand):
    SEQ_TYPES = {'BED file': 'bed',
                 'FASTA file': 'fasta',
                 'Text input': 'text'}

    def __init__(self):
        self.params = {'task': 'Predict'}
        self.validation_hash = {'is_model_file': [],
                                'is_bed': [],
                                'is_fasta': [],
                                'is_multiline_text': [],
                                'is_wig_dir': []}
        self.params['model_folder'] = None

        st.markdown('# Prediction')
        st.markdown('')
        self.general_options()

        st.markdown('## Model')
        self.model_options()

        st.markdown('## Sequences')
        self.sequence_options(self.SEQ_TYPES, evaluation=False)

        st.markdown('')
        self.params['ig'] = st.checkbox('Calculate Integrated Gradients', self.defaults['ig'])
        if self.params['ig']:
            self.params['smoothgrad'] = st.checkbox('Apply smoothgrad method', self.defaults['smoothgrad'])
            st.markdown('###### **WARNING**: Calculating the integrated gradients is a time-consuming process, '
                        'it may take several minutes up to few hours (depending on the number of sequences). ' +
                        'Smoothgrad increases time consumption of integrated gradients by a factor of 20.')

        self.validate_and_run(self.validation_hash)

    def run(self):
        status = st.empty()
        status.text('Preparing sequences...')

        self.params['predict_dir'] = os.path.join(self.params['output_folder'], 'prediction',
                                 f'{str(datetime.datetime.now().strftime("%Y%m%d-%H%M"))}')
        self.ensure_dir(self.params['predict_dir'])

        prepared_file_path = os.path.join(self.params['predict_dir'], 'sequences.tsv')

        if self.params['seq_type'] == 'bed':
            dataset = Dataset(bed_file=self.params['seq_source'], branches=self.params['branches'], category='predict',
                              win=self.params['win'], win_place=self.params['win_place'])
            status.text(f"Mapping intervals to {len(self.params['branches'])} branch(es) and exporting...")
            dataset.sort_datapoints()
        elif self.params['seq_type'] == 'fasta' or self.params['seq_type'] == 'text':
            if self.params['seq_type'] == 'fasta':
                dataset = Dataset(fasta_file=self.params['seq_source'], branches=self.params['branches'], category='predict',
                                  win=self.params['win'], win_place=self.params['win_place'])
            elif self.params['seq_type'] == 'text':
                dataset = Dataset(text_input=self.params['seq_source'], branches=self.params['branches'], category='predict',
                                  win=self.params['win'], win_place=self.params['win_place'])

        dataset.map_to_branches(
            self.references, self.params['strand'], prepared_file_path, status, predict=True, ncpu=self.ncpu)

        predict_x = dataset.encode_branches(dataset, self.params['branches'])

        status.text('Calculating predictions...')

        model = tf.keras.models.load_model(self.params['model_file'])
        predict_y = model.predict(
            predict_x,
            verbose=1)

        for i, klass in enumerate(self.params['klasses']):
            dataset.df[klass] = [y[i] for y in predict_y]
        dataset.df['highest scoring class'] = self.get_klass(predict_y, self.params['klasses'])

        placeholder = st.empty()

        if self.params['ig']:
            status.text('Calculating Integrated Gradients...')
            self.calculate_ig(dataset, model, eval_x, self.params['klasses'], self.params['branches'], self.params['smoothgrad'])

        placeholder.text('Exporting results...')
        result_file = os.path.join(self.params['predict_dir'], 'results.tsv')
        ignore = ['name', 'score', 'klass', 'seq_encoded', 'fold_encoded', 'seq', 'fold', 'cons']
        dataset.save_to_file(ignore_cols=ignore, outfile_path=result_file)

        header = self.predict_header()
        row = self.predict_row(self.params)

        if self.previous_param_file:
            with open(self.previous_param_file, 'r') as file:
                previous_params = yaml.safe_load(file)
            if 'Train' in previous_params.keys():
                # Parameters missing in older versions of the code
                novel_params = {'auc': None, 'avg_precision': None}
                parameters = novel_params
                parameters.update(previous_params['Train'])
                header += f"{self.train_header()}"
                row += f"{self.train_row(parameters)}"
                if 'Preprocess' in previous_params.keys():
                    novel_params = {'win_place': 'rand'}  # It's always been 'random' for the previous versions
                    parameters = novel_params
                    parameters.update(previous_params['Preprocess'])
                    header += f'{self.preprocess_header()}\n'
                    row += f"{self.preprocess_row(parameters)}\n"
                else:
                    header += '\n'
                    row += '\n'
            else:
                header += '\n'
                row += '\n'

        self.finalize_run(logger, self.params['predict_dir'], self.params, header, row, placeholder, self.previous_param_file)
        status.text('Finished!')
        logger.info('Finished!')

    @staticmethod
    def default_params():
        return {
            'model_source': 'from_app',
            'model_folder': '',
            'model_file': '',
            'branches': [],
            'win': 100,
            'no_klasses': 2,
            'klasses': [],
            'seq_type': 'bed',
            'seq_source': '',
            'strand': True,
            'fasta_ref': '',
            'cons_dir': '',
            'win_place': 'center',
            'ig': True,
            'smoothgrad': False,
            'output_folder': os.path.join(os.path.expanduser('~'), 'enngene_output')
        }
```
