## Additional file 1 for "ENNGene: an Easy Neural Network model building tool for Genomics": preprocess.py

```
import datetime
import logging
import os
import re
import shutil
import streamlit as st
import subprocess

from ..utils.dataset import Dataset
from ..utils import file_utils as f
from ..utils.exceptions import UserInputError
from ..utils import sequence as seq
from ..utils.subcommand import Subcommand


logger = logging.getLogger('root')


# noinspection DuplicatedCode
class Preprocess(Subcommand):

    def __init__(self):
        self.params = {'task': 'Preprocess'}
        self.validation_hash = {'is_bed': [],
                                'is_fasta': [],
                                'is_wig_dir': [],
                                'not_empty_branches': [],
                                'is_full_dataset': [],
                                'is_ratio': [],
                                'not_empty_chromosomes': [],
                                'min_two_files': [],
                                'uniq_files': [],
                                'uniq_klasses': []}
        self.params['klasses'] = []
        self.params['full_dataset_dir'] = ''
        self.klass_sizes = {}

        st.markdown('# Preprocessing')
        st.markdown('')

        # TODO add show/hide separate section after stateful operations are allowed
        self.general_options()

        self.params['use_mapped'] = st.checkbox('Use already preprocessed file from a previous run', self.defaults['use_mapped'])

        if not self.params['use_mapped']:
            default_branches = [self.get_dict_key(b, self.BRANCHES) for b in self.defaults['branches']]
            self.params['branches'] = list(map(lambda name: self.BRANCHES[name],
                                               st.multiselect('Branches',
                                                              list(self.BRANCHES.keys()),
                                                              default=default_branches)))
            self.validation_hash['not_empty_branches'].append(self.params['branches'])
            cons_warning = st.empty()

            if 'fold' in self.params['branches']:
                # currently used only as an option for RNAfold
                max_cpu = os.cpu_count() or 1
                self.ncpu = st.slider('Number of CPUs to be used for folding (max = all available CPUs on the machine).',
                                      min_value=1, max_value=max_cpu, value=max_cpu)
            else:
                self.ncpu = 1

            self.references = {}
            if 'seq' in self.params['branches']:
                self.params['strand'] = st.checkbox('Apply strand', self.defaults['strand'])
            if 'seq' in self.params['branches'] or 'fold' in self.params['branches']:
                self.params['fasta'] = st.text_input('Path to the reference fasta file', value=self.defaults['fasta'])
                self.references.update({'seq': self.params['fasta'], 'fold': self.params['fasta']})
                self.validation_hash['is_fasta'].append(self.params['fasta'])
            if 'cons' in self.params['branches']:
                cons_warning.markdown('**WARNING**: Calculating the conservation score is a time-consuming process, '
                                      'it may take up to few hours (based on the size of the wig files).')
                self.params['cons_dir'] = st.text_input('Path to folder containing reference conservation files',
                                                        value=self.defaults['cons_dir'])
                self.references.update({'cons': self.params['cons_dir']})
                self.validation_hash['is_wig_dir'].append(self.params['cons_dir'])

            self.params['win'] = int(st.number_input('Window size', min_value=3, value=self.defaults['win']))
            self.params['win_place'] = self.WIN_PLACEMENT[st.radio(
                'Choose a way to place the window upon the sequence:',
                list(self.WIN_PLACEMENT.keys()), index=self.get_dict_index(self.defaults['win_place'], self.WIN_PLACEMENT))]
            st.markdown('## Input Coordinate Files')

            warning = st.empty()
            no_files = st.number_input('Number of input files (= no. of classes):', min_value=2,
                                       value=max(2, len(self.defaults['input_files'])))
            self.params['input_files'] = [None] * no_files
            for i, file in enumerate(self.defaults['input_files']):
                if len(self.params['input_files']) >= i+1:
                    self.params['input_files'][i] = self.defaults['input_files'][i]
            self.params['klasses'] = [None] * no_files

            self.allowed_extensions = ['.bed', '.narrowPeak']

            for i in range(no_files):
                file = st.text_input(f'File no. {i+1} (.bed)',
                    value=(self.defaults['input_files'][i] if len(self.defaults['input_files']) > i else ''))
                self.params['input_files'][i] = file

                if not file: continue
                self.validation_hash['is_bed'].append({'file': file, 'evaluation': False})
                if os.path.isfile(file):
                    file_name = os.path.basename(file)
                    if any(ext in file_name for ext in self.allowed_extensions):
                        for ext in self.allowed_extensions:
                            if ext in file_name:
                                klass = file_name.replace(ext, '')
                                self.params['klasses'][i] = klass
                                self.klass_sizes.update({klass: (int(subprocess.check_output(['wc', '-l', file]).split()[0]))})
                    else:
                        warning.markdown(
                            '**WARNING**: Only files of following format are allowed: {}.'.format(', '.join(self.allowed_extensions)))
                else:
                    st.markdown(f'##### **WARNING**: Input file no. {i+1} does not exist.')

            self.validation_hash['min_two_files'].append(self.params['input_files'])
            self.validation_hash['uniq_files'].append(self.params['input_files'])
            self.validation_hash['uniq_klasses'].append(self.params['klasses'])
        else:
            # When using already mapped file
            self.params['full_dataset_dir'] = st.text_input(f"Folder from the previous run of the task (must contain 'full_datasets' subfolder)", value=self.defaults['full_dataset_dir'])
            if self.params['full_dataset_dir']:
                self.params['full_dataset_file'] = os.path.join(self.params['full_dataset_dir'], 'full_datasets', 'merged_all.tsv.zip')
                self.validation_hash['is_full_dataset'].append({'file_path': self.params['full_dataset_file'], 'branches': self.params['branches']})

                if self.params['full_dataset_file']:
                    try:
                        self.params['klasses'], self.params['valid_chromosomes'], self.params['branches'], self.klass_sizes = \
                            Dataset.load_and_cache(self.params['full_dataset_file'])
                    except Exception:
                        raise UserInputError('The file with mapped dataset does not exist or is not valid, sorry.')

        st.markdown('## Dataset Size Reduction')
        st.markdown('###### Input a decimal number if you want to reduce the sample size by a ratio (e.g. 0.1 to get 10%), '
                    'or an integer if you wish to select final dataset size (e.g. 5000 if you want exactly 5000 samples).')
        default_reduce = [klass for klass in self.defaults['reducelist'] if klass in self.params['klasses']]
        self.params['reducelist'] = st.multiselect('Classes to be reduced (first specify input files)',
                                                   self.params['klasses'], default_reduce)
        if self.params['reducelist']:
            default_ratio = {k: v for k, v in self.defaults['reduceratio'].items() if k in self.params['klasses']}
            self.params['reduceratio'] = default_ratio
            for klass in self.params['reducelist']:
                if klass not in self.params['reduceratio'].keys():
                    self.params['reduceratio'][klass] = 0.5
                self.params['reduceratio'].update({klass: float(st.number_input(
                    f'Target {klass} dataset size (original size: {self.klass_sizes[klass]} rows)',
                    min_value=0.00001, value=default_ratio[klass], format='%.2f'))})
            st.markdown('###### WARNING: The data are reduced randomly across the dataset. Thus in a rare occasion, when later '
                    'splitting the dataset by chromosomes, some categories may end up empty. Thus it\'s recommended '
                    'to be used in combination with random split.')

        st.markdown('## Data Split')
        split_options = {'Random': 'rand',
                         'By chromosomes': 'by_chr'}
        self.params['split'] = split_options[st.radio(
            'Choose a way to split datasets into train, test, validation and blackbox categories:',
            list(split_options.keys()), index=self.get_dict_index(self.defaults['split'], split_options))]
        if self.params['split'] == 'by_chr':
            chr_ready = False
            if self.params['use_mapped']:
                if self.params['full_dataset_file']:
                    chr_ready = True
                else:
                    st.markdown('**The mapped file must be provided first to infer available chromosomes.**')
            else:
                if 'seq' in self.params['branches'] or 'fold' in self.params['branches']:
                    if self.params['fasta']:
                        try:
                            self.params['valid_chromosomes'] = seq.read_and_cache(self.params['fasta'])
                            chr_ready = True
                        except Exception:
                            raise UserInputError('Sorry, could not parse given fasta file. Please check the path.')
                    else:
                        st.markdown('**Fasta file with reference genome must be provided to infer available chromosomes.**')
                elif 'cons' in self.params['branches']:
                    st.markdown('###### WARNING: When conservation score branch selected only, the split is done based on separate wig files provided. '
                                'Note that to be able to do that, the wig files must contain the chromosome name in the exact same form as your bed files and must not contain dots within the chromosome name.')
                    if self.params['cons_dir']:
                        chrom_files = f.list_files_in_dir(self.params['cons_dir'], 'wig')
                        chromosomes = []
                        for file in chrom_files:
                            match = re.search(r'.*\.*.*(chr[^.]*)\..*', os.path.basename(file))
                            if match and match.group(1):
                                chromosomes.append(match.group(1))
                        self.params['valid_chromosomes'] = list(set(chromosomes))
                        chr_ready = True
                    else:
                        st.markdown('**Folder with conservation score (wig) files must be provided to infer available chromosomes.**')
                else:
                    st.markdown('**Please choose at least one branch, and provide necessary reference files to infer available chromosomes.**')

            if chr_ready:
                if self.params['valid_chromosomes']:
                    if self.params['fasta'] and not self.params['use_mapped']:
                        st.markdown("##### WARNING: While selecting the chromosomes, you may ignore the yellow warning box, \
                                    and continue selecting even while it's present, as long as you work within one selectbox "
                                    "(e.g. you can select multiple chromosomes within training dataset, but than "
                                    "you have to wait until the warning disappears before you start working with the validation set).")
                    self.params['chromosomes'] = self.defaults['chromosomes']
                    self.params['chromosomes'].update({'train': set(st.multiselect(
                        'Training Dataset', self.params['valid_chromosomes'], list(self.defaults['chromosomes']['train'])))})
                    self.params['chromosomes'].update({'validation': set(
                       st.multiselect('Validation Dataset', self.params['valid_chromosomes'], list(self.defaults['chromosomes']['validation'])))})
                    self.params['chromosomes'].update({'test': set(
                        st.multiselect('Test Dataset', self.params['valid_chromosomes'], list(self.defaults['chromosomes']['test'])))})
                    self.params['chromosomes'].update({'blackbox': set(
                      st.multiselect('BlackBox Dataset (optional)', self.params['valid_chromosomes'], list(self.defaults['chromosomes']['blackbox'])))})
                    self.validation_hash['not_empty_chromosomes'].append(list(self.params['chromosomes'].items()))
                else:
                    raise UserInputError('Sorry, did not find any valid chromosomes in given fasta file.')

        elif self.params['split'] == 'rand':
            self.params['split_ratio'] = st.text_input(
                'List a target ratio between the categories (required format: train:validation:test:blackbox)',
                value=self.defaults['split_ratio'])
            self.validation_hash['is_ratio'].append(self.params['split_ratio'])
            st.markdown('###### Note: If you do not want to use the blackbox dataset (for later evaluation), you can just set it\'s size to 0.')

        self.validate_and_run(self.validation_hash)

    def run(self):
        status = st.empty()

        self.params['datasets_dir'] = os.path.join(self.params['output_folder'], 'datasets', f'{str(datetime.datetime.now().strftime("%Y-%m-%d_%H:%M"))}')
        self.ensure_dir(self.params['datasets_dir'])
        full_data_dir_path = os.path.join(self.params['datasets_dir'], 'full_datasets')
        self.ensure_dir(full_data_dir_path)
        full_data_file_path = os.path.join(full_data_dir_path, 'merged_all.tsv')

        if self.params['use_mapped']:
            status.text('Reading in already mapped file with all the samples...')
            merged_dataset = Dataset.load_from_file(self.params['full_dataset_file'])
            # FIXME copied file is broken
            shutil.copyfile(self.params['full_dataset_file'], full_data_file_path)
            # Keep only selected branches
            cols = ['chrom_name', 'seq_start', 'seq_end', 'strand_sign', 'klass'] + self.params['branches']
            merged_dataset.df = merged_dataset.df[cols]
        else:
            # Accept one file per class and create one Dataset per each
            initial_datasets = set()
            status.text('Reading in given interval files and applying window...')
            for file in self.params['input_files']:
                klass = os.path.basename(file)
                for ext in self.allowed_extensions:
                    if ext in klass:
                        klass = klass.replace(ext, '')

                initial_datasets.add(
                    Dataset(klass=klass, branches=self.params['branches'], bed_file=file, win=self.params['win'],
                            win_place=self.params['win_place']))

            # Merging data from all klasses to map them more efficiently all together at once
            merged_dataset = Dataset(branches=self.params['branches'], df=Dataset.merge_dataframes(initial_datasets))

            # First ensure order of the data by chr_name and seq_start within, mainly for conservation
            status.text(
                f"Mapping all intervals from to {len(self.params['branches'])} branch(es) and exporting...")
            merged_dataset.sort_datapoints().map_to_branches(
                self.references, self.params['strand'], full_data_file_path, status, self.ncpu)

        status.text('Processing mapped samples...')
        mapped_datasets = set()
        for klass in self.params['klasses']:
            df = merged_dataset.df[merged_dataset.df['klass'] == klass]
            mapped_datasets.add(Dataset(klass=klass, branches=self.params['branches'], df=df))

        split_datasets = set()
        for dataset in mapped_datasets:
            # Reduce size of selected klasses
            if self.params['reducelist'] and (dataset.klass in self.params['reducelist']):
                status.text(f'Reducing number of samples in klass {format(dataset.klass)}...')
                ratio = self.params['reduceratio'][dataset.klass]
                dataset.reduce(ratio)

            # Split datasets into train, validation, test and blackbox datasets
            if self.params['split'] == 'by_chr':
                split_subdatasets = Dataset.split_by_chr(dataset, self.params['chromosomes'])
            elif self.params['split'] == 'rand':
                split_subdatasets = Dataset.split_random(dataset, self.params['split_ratio'])
            split_datasets = split_datasets.union(split_subdatasets)

        # Merge datasets of the same category across all the branches (e.g. train = pos + neg)
        status.text('Redistributing samples to categories and exporting into final files...')
        final_datasets = Dataset.merge_by_category(split_datasets)

        for dataset in final_datasets:
            dir_path = os.path.join(self.params['datasets_dir'], 'final_datasets')
            self.ensure_dir(dir_path)
            file_path = os.path.join(dir_path, f'{dataset.category}.tsv')
            dataset.save_to_file(file_path, ignore_cols=['name', 'score'], do_zip=True)

        self.finalize_run(logger, self.params['datasets_dir'], self.params,
                          f'{self.preprocess_header()} \n',
                          f'{self.preprocess_row(self.params)} \n')
        status.text('Finished!')
        logger.info('Finished!')

    @staticmethod
    def default_params():
        return {'branches': [],
                'chromosomes': {'train': [], 'validation': [], 'test': [], 'blackbox': []},
                'cons_dir': '',
                'fasta': '',
                'full_dataset_dir': '',
                'full_dataset_file': '',
                'input_files': [],
                'output_folder': os.path.join(os.path.expanduser('~'), 'enngene_output'),
                'reducelist': [],
                'reduceratio': {},
                'split': 'rand',
                'split_ratio': '7:1:1:1',
                'strand': True,
                'use_mapped': False,
                'valid_chromosomes': [],
                'win': 100,
                'win_place': 'center',
                }
```
