## Additional file 1 for "ENNGene: an Easy Neural Network model building tool for Genomics": callbacks.py

```
import logging
import numpy as np
import os
import pandas as pd
import tensorflow as tf

logger = logging.getLogger('root')


class ProgressMonitor(tf.keras.callbacks.Callback):

    def __init__(self, epochs, progress_bar=None, progress_status=None, chart=None, metrics=None):
        self.epochs = epochs
        self.progress_bar = progress_bar
        self.progress_status = progress_status
        self.chart = chart
        self.metrics = metrics

    def on_epoch_begin(self, epoch, logs=None):
        self.progress_bar.progress((epoch+1)/self.epochs)
        self.progress_status.text(f'Epoch {epoch+1}/{self.epochs}')

    def on_epoch_end(self, epoch, logs=None):
        epoch_data = pd.DataFrame([['Training loss', round(logs['loss'], 2), epoch+1],
                                   ['Validation loss', round(logs['val_loss'], 2), epoch+1]],
                                  columns=['Metric', 'Metric value', 'Epoch'])
        for metric in self.metrics:
            # For now plot only acc and auc during the training
            metric_name = metric if metric == 'accuracy' else metric.name
            if metric_name == 'auc' or metric_name == 'accuracy':
                metric_df = pd.DataFrame([[f'Training {metric_name}', round(logs[metric_name], 2), epoch+1],
                                          [f'Validation {metric_name}', round(logs[f'val_{metric_name}'], 2), epoch+1]],
                                         columns=['Metric', 'Metric value', 'Epoch'])
                epoch_data = epoch_data.append(metric_df, ignore_index=True)

        self.chart.add_rows(epoch_data)


# Code is authored by https://github.com/titu1994/keras-one-cycle, originally ported from https://github.com/fastai/fastai
# See the LICENSE at the onecycle_LICENSE file.
class OneCycleLR(tf.keras.callbacks.Callback):

    def __init__(self,
                 max_lr,
                 end_percentage=0.1,
                 scale_percentage=None,
                 maximum_momentum=0.95,
                 minimum_momentum=0.85,
                 verbose=True):
        """ This callback implements a cyclical learning rate policy (CLR).
        This is a special case of Cyclic Learning Rates, where we have only 1 cycle.
        After the completion of 1 cycle, the learning rate will decrease rapidly to
        100th its initial lowest value.

        # Arguments:
            max_lr: Float. Initial learning rate. This also sets the
                starting learning rate (which will be 10x smaller than
                this), and will increase to this value during the first cycle.
            end_percentage: Float. The percentage of all the epochs of training
                that will be dedicated to sharply decreasing the learning
                rate after the completion of 1 cycle. Must be between 0 and 1.
            scale_percentage: Float or None. If float, must be between 0 and 1.
                If None, it will compute the scale_percentage automatically
                based on the `end_percentage`.
            maximum_momentum: Optional. Sets the maximum momentum (initial)
                value, which gradually drops to its lowest value in half-cycle,
                then gradually increases again to stay constant at this max value.
                Can only be used with SGD Optimizer.
            minimum_momentum: Optional. Sets the minimum momentum at the end of
                the half-cycle. Can only be used with SGD Optimizer.
            verbose: Bool. Whether to print the current learning rate after every
                epoch.

        # Reference
            - [A disciplined approach to neural network hyper-parameters: Part 1 -- learning rate, batch size, weight_decay, and weight decay](https://arxiv.org/abs/1803.09820)
            - [Super-Convergence: Very Fast Training of Residual Networks Using Large Learning Rates](https://arxiv.org/abs/1708.07120)
        """
        super(OneCycleLR, self).__init__()

        if end_percentage < 0. or end_percentage > 1.:
            raise ValueError("`end_percentage` must be between 0 and 1")

        if scale_percentage is not None and (scale_percentage < 0. or scale_percentage > 1.):
            raise ValueError("`scale_percentage` must be between 0 and 1")

        self.initial_lr = max_lr
        self.end_percentage = end_percentage
        self.scale = float(scale_percentage) if scale_percentage is not None else float(end_percentage)
        self.max_momentum = maximum_momentum
        self.min_momentum = minimum_momentum
        self.verbose = verbose

        if self.max_momentum is not None and self.min_momentum is not None:
            self._update_momentum = True
        else:
            self._update_momentum = False

        self.clr_iterations = 0.
        self.history = {}

        self.epochs = None
        self.batch_size = None
        self.samples = None
        self.steps = None
        self.num_iterations = None
        self.mid_cycle_id = None

    def _reset(self):
        """
        Reset the callback.
        """
        self.clr_iterations = 0.
        self.history = {}

    def compute_lr(self):
        """
        Compute the learning rate based on which phase of the cycle it is in.

        - If in the first half of training, the learning rate gradually increases.
        - If in the second half of training, the learning rate gradually decreases.
        - If in the final `end_percentage` portion of training, the learning rate
            is quickly reduced to near 100th of the original min learning rate.

        # Returns:
            the new learning rate
        """
        if self.clr_iterations > 2 * self.mid_cycle_id:
            current_percentage = (self.clr_iterations - 2 * self.mid_cycle_id)
            current_percentage /= float((self.num_iterations - 2 * self.mid_cycle_id))
            new_lr = self.initial_lr * (1. + (current_percentage *
                                              (1. - 100.) / 100.)) * self.scale

        elif self.clr_iterations > self.mid_cycle_id:
            current_percentage = 1. - (
                    self.clr_iterations - self.mid_cycle_id) / self.mid_cycle_id
            new_lr = self.initial_lr * (1. + current_percentage *
                                        (self.scale * 100 - 1.)) * self.scale

        else:
            current_percentage = self.clr_iterations / self.mid_cycle_id
            new_lr = self.initial_lr * (1. + current_percentage *
                                        (self.scale * 100 - 1.)) * self.scale

        if self.clr_iterations == self.num_iterations:
            self.clr_iterations = 0

        return new_lr

    def compute_momentum(self):
        """
         Compute the momentum based on which phase of the cycle it is in.

        - If in the first half of training, the momentum gradually decreases.
        - If in the second half of training, the momentum gradually increases.
        - If in the final `end_percentage` portion of training, the momentum value
            is kept constant at the maximum initial value.

        # Returns:
            the new momentum value
        """
        if self.clr_iterations > 2 * self.mid_cycle_id:
            new_momentum = self.max_momentum

        elif self.clr_iterations > self.mid_cycle_id:
            current_percentage = 1. - ((self.clr_iterations - self.mid_cycle_id) / float(
                self.mid_cycle_id))
            new_momentum = self.max_momentum - current_percentage * (
                    self.max_momentum - self.min_momentum)

        else:
            current_percentage = self.clr_iterations / float(self.mid_cycle_id)
            new_momentum = self.max_momentum - current_percentage * (
                    self.max_momentum - self.min_momentum)

        return new_momentum

    def on_train_begin(self, logs={}):
        logs = logs or {}

        self.epochs = self.params['epochs']
        self.batch_size = self.params['batch_size']
        self.samples = self.params['samples']
        self.steps = self.params['steps']

        if self.steps is not None:
            self.num_iterations = self.epochs * self.steps
        else:
            if (self.samples % self.batch_size) == 0:
                remainder = 0
            else:
                remainder = 1
            self.num_iterations = (self.epochs + remainder) * self.samples // self.batch_size

        self.mid_cycle_id = int(self.num_iterations * ((1. - self.end_percentage)) / float(2))

        self._reset()
        tf.keras.backend.set_value(self.model.optimizer.lr, self.compute_lr())

        if self._update_momentum:
            if not hasattr(self.model.optimizer, 'momentum'):
                raise ValueError("Momentum can be updated only on SGD optimizer !")

            new_momentum = self.compute_momentum()
            tf.keras.backend.set_value(self.model.optimizer.momentum, new_momentum)

    def on_batch_end(self, epoch, logs=None):
        logs = logs or {}

        self.clr_iterations += 1
        new_lr = self.compute_lr()

        self.history.setdefault('lr', []).append(
            tf.keras.backend.get_value(self.model.optimizer.lr))
        tf.keras.backend.set_value(self.model.optimizer.lr, new_lr)

        if self._update_momentum:
            if not hasattr(self.model.optimizer, 'momentum'):
                raise ValueError("Momentum can be updated only on SGD optimizer !")

            new_momentum = self.compute_momentum()

            self.history.setdefault('momentum', []).append(
                tf.keras.backend.get_value(self.model.optimizer.momentum))
            tf.keras.backend.set_value(self.model.optimizer.momentum, new_momentum)

        for k, v in logs.items():
            self.history.setdefault(k, []).append(v)

    def on_epoch_end(self, epoch, logs=None):
        if self.verbose:
            if self._update_momentum:
                logger.info(" - lr: %0.5f - momentum: %0.2f " %
                      (self.history['lr'][-1], self.history['momentum'][-1]))

            else:
                logger.info(" - lr: %0.5f " % (self.history['lr'][-1]))


# Code is authored by https://github.com/titu1994/keras-one-cycle, originally ported from https://github.com/fastai/fastai
class LRFinder(tf.keras.callbacks.Callback):

    def __init__(self,
                 num_samples,
                 batch_size,
                 minimum_lr=1e-5,
                 maximum_lr=10.,
                 lr_scale='exp',
                 validation_data=None,
                 validation_sample_rate=5,
                 stopping_criterion_factor=4.,
                 loss_smoothing_beta=0.98,
                 save_dir=None,
                 verbose=True):
        """
        This class uses the Cyclic Learning Rate history to find a
        set of learning rates that can be good initializations for the
        One-Cycle training proposed by Leslie Smith in the paper referenced
        below.

        A port of the Fast.ai implementation for Keras.

        # Note
        This requires that the model be trained for exactly 1 epoch. If the model
        is trained for more epochs, then the metric calculations are only done for
        the first epoch.

        # Interpretation
        Upon visualizing the loss plot, check where the loss starts to increase
        rapidly. Choose a learning rate at somewhat prior to the corresponding
        position in the plot for faster convergence. This will be the maximum_lr lr.
        Choose the max value as this value when passing the `max_val` argument
        to OneCycleLR callback.

        Since the plot is in log-scale, you need to compute 10 ^ (-k) of the x-axis

        # Arguments:
            num_samples: Integer. Number of samples in the dataset.
            batch_size: Integer. Batch size during training.
            minimum_lr: Float. Initial learning rate (and the minimum).
            maximum_lr: Float. Final learning rate (and the maximum).
            lr_scale: Can be one of ['exp', 'linear']. Chooses the type of
                scaling for each update to the learning rate during subsequent
                batches. Choose 'exp' for large range and 'linear' for small range.
            validation_data: Requires the validation dataset as a tuple of
                (X, y) belonging to the validation set. If provided, will use the
                validation set to compute the loss metrics. Else uses the training
                batch loss. Will warn if not provided to alert the user.
            validation_sample_rate: Positive or Negative Integer. Number of batches to sample from the
                validation set per iteration of the LRFinder. Larger number of
                samples will reduce the variance but will take longer time to execute
                per batch.

                If Positive > 0, will sample from the validation dataset
                If Megative, will use the entire dataset
            stopping_criterion_factor: Integer or None. A factor which is used
                to measure large increase in the loss value during training.
                Since callbacks cannot stop training of a model, it will simply
                stop logging the additional values from the epochs after this
                stopping criterion has been met.
                If None, this check will not be performed.
            loss_smoothing_beta: Float. The smoothing factor for the moving
                average of the loss function.
            save_dir: Optional, String. If passed a directory path, the callback
                will save the running loss and learning rates to two separate numpy
                arrays inside this directory. If the directory in this path does not
                exist, they will be created.
            verbose: Whether to print the learning rate after every batch of training.

        # References:
            - [A disciplined approach to neural network hyper-parameters: Part 1 -- learning rate, batch size, weight_decay, and weight decay](https://arxiv.org/abs/1803.09820)
        """
        super(LRFinder, self).__init__()

        if lr_scale not in ['exp', 'linear']:
            raise ValueError("`lr_scale` must be one of ['exp', 'linear']")

        if validation_data is not None:
            self.validation_data = validation_data
            self.use_validation_set = True

            if validation_sample_rate > 0 or validation_sample_rate < 0:
                self.validation_sample_rate = validation_sample_rate
            else:
                raise ValueError("`validation_sample_rate` must be a positive or negative integer other than 0")
        else:
            self.use_validation_set = False
            self.validation_sample_rate = 0

        self.num_samples = num_samples
        self.batch_size = batch_size
        self.initial_lr = minimum_lr
        self.final_lr = maximum_lr
        self.lr_scale = lr_scale
        self.stopping_criterion_factor = stopping_criterion_factor
        self.loss_smoothing_beta = loss_smoothing_beta
        self.save_dir = save_dir
        self.verbose = verbose

        self.num_batches_ = num_samples // batch_size
        self.current_lr_ = minimum_lr

        if lr_scale == 'exp':
            self.lr_multiplier_ = (maximum_lr / float(minimum_lr)) ** (
                    1. / float(self.num_batches_))
        else:
            extra_batch = int((num_samples % batch_size) != 0)
            self.lr_multiplier_ = np.linspace(
                minimum_lr, maximum_lr, num=self.num_batches_ + extra_batch)

        # If negative, use entire validation set
        if self.validation_sample_rate < 0:
            self.validation_sample_rate = self.validation_data[0].shape[0] // batch_size

        self.current_batch_ = 0
        self.current_epoch_ = 0
        self.best_loss_ = 1e6
        self.running_loss_ = 0.

        self.history = {}

    def on_train_begin(self, logs=None):

        self.current_epoch_ = 1
        tf.keras.backend.set_value(self.model.optimizer.lr, self.initial_lr)

    def on_epoch_begin(self, epoch, logs=None):
        self.current_batch_ = 0

        if self.current_epoch_ > 1:
            msg = "\n\nLearning rate finder should be used only with a single epoch. " \
                       "Hereafter, the callback will not measure the losses.\n\n"
            logger.warning(msg)

    def on_batch_begin(self, batch, logs=None):
        self.current_batch_ += 1

    def on_batch_end(self, batch, logs=None):
        if self.current_epoch_ > 1:
            return

        if self.use_validation_set:
            X, Y = self.validation_data[0], self.validation_data[1]

            # use 5 random batches from test set for fast approximate of loss
            num_samples = self.batch_size * self.validation_sample_rate

            if num_samples > X.shape[0]:
                num_samples = X.shape[0]

            idx = np.random.choice(X.shape[0], num_samples, replace=False)
            x = X[idx]
            y = Y[idx]

            values = self.model.evaluate(x, y, batch_size=self.batch_size, verbose=False)
            loss = values[0]
        else:
            loss = logs['loss']

        # smooth the loss value and bias correct
        running_loss = self.loss_smoothing_beta * loss + (
                1. - self.loss_smoothing_beta) * loss
        running_loss = running_loss / (
                1. - self.loss_smoothing_beta**self.current_batch_)

        # stop logging if loss is too large
        if self.current_batch_ > 1 and self.stopping_criterion_factor is not None and (
                running_loss >
                self.stopping_criterion_factor * self.best_loss_):

            if self.verbose:
                logger.info(" - LRFinder: Skipping iteration since loss is %d times as large as best loss (%0.4f)"
                      % (self.stopping_criterion_factor, self.best_loss_))
            return

        if running_loss < self.best_loss_ or self.current_batch_ == 1:
            self.best_loss_ = running_loss

        current_lr = tf.keras.backend.get_value(self.model.optimizer.lr)

        self.history.setdefault('running_loss_', []).append(running_loss)
        if self.lr_scale == 'exp':
            self.history.setdefault('log_lrs', []).append(np.log10(current_lr))
        else:
            self.history.setdefault('log_lrs', []).append(current_lr)

        # compute the lr for the next batch and update the optimizer lr
        if self.lr_scale == 'exp':
            current_lr *= self.lr_multiplier_
        else:
            current_lr = self.lr_multiplier_[self.current_batch_ - 1]

        tf.keras.backend.set_value(self.model.optimizer.lr, current_lr)

        # save the other metrics as well
        for k, v in logs.items():
            self.history.setdefault(k, []).append(v)

        if self.verbose:
            if self.use_validation_set:
                logger.info(" - LRFinder: val_loss: %1.4f - lr = %1.8f " %
                      (values[0], current_lr))
            else:
                logger.info(" - LRFinder: lr = %1.8f " % current_lr)

    def on_epoch_end(self, epoch, logs=None):
        if self.save_dir is not None and self.current_epoch_ <= 1:
            if not os.path.exists(self.save_dir):
                os.makedirs(self.save_dir)

            losses_path = os.path.join(self.save_dir, 'losses.npy')
            lrs_path = os.path.join(self.save_dir, 'lrs.npy')

            np.save(losses_path, self.losses)
            np.save(lrs_path, self.lrs)

            if self.verbose:
                logger.info("\tLR Finder : Saved the losses and learning rate values in path : {%s}" % (self.save_dir))

        self.current_epoch_ += 1

    def plot_schedule(self, clip_beginning=None, clip_endding=None):
        """
        Plots the schedule from the callback itself.

        # Arguments:
            clip_beginning: Integer or None. If positive integer, it will
                remove the specified portion of the loss graph to remove the large
                loss values in the beginning of the graph.
            clip_endding: Integer or None. If negative integer, it will
                remove the specified portion of the ending of the loss graph to
                remove the sharp increase in the loss values at high learning rates.
        """
        try:
            import matplotlib.pyplot as plt
            plt.style.use('seaborn-white')
        except ImportError:
            logger.warning("Matplotlib not found. Please use `pip install matplotlib` first.")
            return

        if clip_beginning is not None and clip_beginning < 0:
            clip_beginning = -clip_beginning

        if clip_endding is not None and clip_endding > 0:
            clip_endding = -clip_endding

        losses = self.losses
        lrs = self.lrs

        if clip_beginning:
            losses = losses[clip_beginning:]
            lrs = lrs[clip_beginning:]

        if clip_endding:
            losses = losses[:clip_endding]
            lrs = lrs[:clip_endding]

        plt.plot(lrs, losses)
        plt.title('Learning rate vs Loss')
        plt.xlabel('learning rate')
        plt.ylabel('loss')
        plt.show()

    @classmethod
    def restore_schedule_from_dir(cls,
                                  directory,
                                  clip_beginning=None,
                                  clip_endding=None):
        """
        Loads the training history from the saved numpy files in the given directory.

        # Arguments:
            directory: String. Path to the directory where the serialized numpy
                arrays of the loss and learning rates are saved.
            clip_beginning: Integer or None. If positive integer, it will
                remove the specified portion of the loss graph to remove the large
                loss values in the beginning of the graph.
            clip_endding: Integer or None. If negative integer, it will
                remove the specified portion of the ending of the loss graph to
                remove the sharp increase in the loss values at high learning rates.

        Returns:
            tuple of (losses, learning rates)
        """
        if clip_beginning is not None and clip_beginning < 0:
            clip_beginning = -clip_beginning

        if clip_endding is not None and clip_endding > 0:
            clip_endding = -clip_endding

        losses_path = os.path.join(directory, 'losses.npy')
        lrs_path = os.path.join(directory, 'lrs.npy')

        if not os.path.exists(losses_path) or not os.path.exists(lrs_path):
            logger.info("%s and %s could not be found at directory : {%s}" %
                  (losses_path, lrs_path, directory))

            losses = None
            lrs = None

        else:
            losses = np.load(losses_path)
            lrs = np.load(lrs_path)

            if clip_beginning:
                losses = losses[clip_beginning:]
                lrs = lrs[clip_beginning:]

            if clip_endding:
                losses = losses[:clip_endding]
                lrs = lrs[:clip_endding]

        return losses, lrs

    @classmethod
    def plot_schedule_from_file(cls,
                                directory,
                                clip_beginning=None,
                                clip_endding=None):
        """
        Plots the schedule from the saved numpy arrays of the loss and learning
        rate values in the specified directory.

        # Arguments:
            directory: String. Path to the directory where the serialized numpy
                arrays of the loss and learning rates are saved.
            clip_beginning: Integer or None. If positive integer, it will
                remove the specified portion of the loss graph to remove the large
                loss values in the beginning of the graph.
            clip_endding: Integer or None. If negative integer, it will
                remove the specified portion of the ending of the loss graph to
                remove the sharp increase in the loss values at high learning rates.
        """
        try:
            import matplotlib.pyplot as plt
            plt.style.use('seaborn-white')
        except ImportError:
            logger.warning("Matplotlib not found. Please use `pip install matplotlib` first.")
            return

        losses, lrs = cls.restore_schedule_from_dir(
            directory,
            clip_beginning=clip_beginning,
            clip_endding=clip_endding)

        if losses is None or lrs is None:
            return
        else:
            plt.plot(lrs, losses)
            plt.title('Learning rate vs Loss')
            plt.xlabel('learning rate')
            plt.ylabel('loss')
            plt.savefig(directory + '/lr_finder', dpi=300)
            plt.clf()

    @property
    def lrs(self):
        return np.array(self.history['log_lrs'])

    @property
    def losses(self):
        return np.array(self.history['running_loss_'])
```
