## Additional file 1 for "ENNGene: an Easy Neural Network model building tool for Genomics": layers.py

```
from tensorflow.keras.layers import Dense, Dropout, Conv1D, GRU, MaxPooling1D, BatchNormalization, \
    LeakyReLU, LSTM, Bidirectional


class MyConv1D:

    @staticmethod
    def build(x, filters=40, kernel=4, batchnorm=False, dropout=None, **kwargs):
        x = Conv1D(filters=filters, kernel_size=kernel, padding='same')(x)
        x = LeakyReLU()(x)
        if batchnorm: x = BatchNormalization()(x)
        x = MaxPooling1D(pool_size=2, padding='same')(x)
        if dropout: x = Dropout(rate=dropout, noise_shape=None, seed=None)(x)
        return x

#
# class MyLocallyConnected1D:
#
#     @staticmethod
#     def build(x, filters=40, kernel=4, batchnorm=False, dropout=None, **kwargs):
#         x = LocallyConnected1D(filters=filters, kernel_size=kernel, padding='valid')(x)
#         x = LeakyReLU()(x)
#         if batchnorm: x = BatchNormalization()(x)
#         x = MaxPooling1D(pool_size=2, padding='same')(x)
#         if dropout: x = Dropout(rate=dropout, noise_shape=None, seed=None)(x)
#         return x
#

class MyDense:

    @staticmethod
    def build(x, units=32, batchnorm=False, dropout=None, **kwargs):
        x = Dense(units)(x)
        x = LeakyReLU()(x)
        if batchnorm: x = BatchNormalization()(x)
        if dropout: x = Dropout(rate=dropout, noise_shape=None, seed=None)(x)
        return x


class MyLSTM:

    @staticmethod
    def build(x, units=32, bidirect=True, return_seq=False, batchnorm=False, dropout=None, **kwargs):
        if bidirect:
            x = Bidirectional(LSTM(units, return_sequences=return_seq))(x)
        else:
            x = LSTM(units, return_sequences=return_seq)(x)
        if batchnorm: x = BatchNormalization()(x)
        if dropout: x = Dropout(rate=dropout, noise_shape=None, seed=None)(x)
        return x


class MyGRU:

    @staticmethod
    def build(x, units=32, bidirect=True, return_seq=False, batchnorm=False, dropout=None, **kwargs):
        if bidirect:
            x = Bidirectional(GRU(units, return_sequences=return_seq))(x)
        else:
            x = GRU(units, return_sequences=return_seq)(x)
        if batchnorm: x = BatchNormalization()(x)
        if dropout: x = Dropout(rate=dropout, noise_shape=None, seed=None)(x)
        return x


LAYERS = {'Convolution layer': MyConv1D,
          # 'Locally Connected 1D layer': MyLocallyConnected1D,
          'Dense layer': MyDense,
          'GRU': MyGRU,
          'LSTM': MyLSTM}
BRANCH_LAYERS = {'Convolution layer': MyConv1D}
                 # 'Locally Connected 1D layer': MyLocallyConnected1D}
COMMON_LAYERS = {'Dense layer': MyDense,
                 'GRU': MyGRU,
                 'LSTM': MyLSTM}
```
