## Additional file 1 for "ENNGene: an Easy Neural Network model building tool for Genomics": model_builder.py

```
import tensorflow as tf
from tensorflow.keras.layers import Dense, Flatten, Input
from tensorflow.keras.models import Model

from .layers import LAYERS


class ModelBuilder:

    def __init__(self, branches=None, labels=None, branch_shapes=None, branches_layers=None, common_layers=None):
        if not branches and labels and branch_shapes and branches_layers and common_layers:
            raise AttributeError('Missing parameters to build a model.')
        self.branches = branches
        self.labels = labels
        self.branch_shapes = branch_shapes
        self.branches_layers = branches_layers

        if len(common_layers) == 1:
            self.common_layers = common_layers
        else:  # when stacking multiple RNNs the first layers must return sequences
            updated_common = []
            for i, layer in enumerate(common_layers):
                if layer['name'] in ('GRU', 'LSTM') and i+1 != len(common_layers) and\
                        (common_layers[i+1]['name'] in ('GRU', 'LSTM')):
                    layer['args'].update({'return_seq': True})
                updated_common.append(layer)
            self.common_layers = updated_common

    def build_model(self):
        inputs = []
        branches_models = []
        for branch in self.branches:
            # batch size left undefined, thus variable
            x = Input(shape=(self.branch_shapes[branch][1:]))
            inputs.append(x)

            for layer in self.branches_layers[branch]:
                x = LAYERS[layer['name']]().build(x, **layer['args'])
            branches_models.append(x)

        if len(self.branches) == 1:
            x = branches_models[0]
        else:
            x = tf.keras.layers.concatenate(branches_models)

        for i, layer in enumerate(self.common_layers):
            # we must flatten the output right before the first Dense layer
            if layer['name'] == 'Dense layer' and ((i == 0) or self.common_layers[i-1]['name'] != 'Dense layer'):
                x = Flatten()(x)
            x = LAYERS[layer['name']]().build(x, **layer['args'])

        output = Dense(units=len(self.labels), activation='softmax')(x)

        if len(self.branches) == 1:
            model = Model(inputs[0], output)
        else:
            model = Model(inputs, output)

        return model
```
