## Additional file 1 for "ENNGene: an Easy Neural Network model building tool for Genomics": train.py

```
import altair as alt
import datetime
import copy
import os
import logging
import math
import matplotlib.pyplot as plt
import numpy as np
import pandas as pd
import streamlit as st
import tensorflow as tf
import yaml

from tensorflow.keras.callbacks import ModelCheckpoint, EarlyStopping, CSVLogger, LearningRateScheduler, TensorBoard
from tensorflow.keras.optimizers import SGD, RMSprop, Adam

from .callbacks import ProgressMonitor, LRFinder, OneCycleLR
from .layers import BRANCH_LAYERS, COMMON_LAYERS
from .model_builder import ModelBuilder
from ..utils.dataset import Dataset
from ..utils.exceptions import UserInputError
from ..utils import file_utils as f
from ..utils import sequence as seq
from ..utils.subcommand import Subcommand

logger = logging.getLogger('root')


class Train(Subcommand):
    TRAIN_METRICS = {'accuracy': 'Accuracy (threshold = 0.5)'}
        # tf.keras.metrics.AUC(name='auc'): 'AUC',  # TODO get implementation for softmax fc
        # tf.keras.metrics.Precision(name='precision'): 'Precision (threshold = 0.5)',
        # tf.keras.metrics.Recall(name='recall'): 'Recall (threshold = 0.5)'

    def __init__(self):
        self.params = {'task': 'Train'}
        self.validation_hash = {'not_empty_branches': [],
                                'is_dataset_dir': []}

        st.markdown('# Training')
        st.markdown('')

        self.general_options()

        self.params['input_folder'] = st.text_input(
            'Datasets folder', value=self.defaults['input_folder'])
        self.validation_hash['is_dataset_dir'].append(self.params['input_folder'])

        if self.params['input_folder']:
            if not os.path.isdir(self.params['input_folder']):
                raise UserInputError('Given folder does not seem to exist.')
            note = ''
            previous_param_files = [f for f in os.listdir(self.params['input_folder']) if (f == 'parameters.yaml') and
                                    (os.path.isfile(os.path.join(self.params['input_folder'], f)))]
            if len(previous_param_files) == 1:
                self.previous_param_file = os.path.join(self.params['input_folder'], previous_param_files[0])
                with open(self.previous_param_file, 'r') as file:
                    user_params = yaml.safe_load(file)
                    preprocess_branches = user_params['Preprocess']['branches']
            else:
                # We could allow to continue without it, but the user would have to select correct branches
                raise UserInputError('Did not find parameters.yaml file in the given dataset folder.')
        else:
            preprocess_branches = []
            note = ' (You must first provide the dataset folder (containing also the parameters.yaml file))'

        available_branches = [self.get_dict_key(b, self.BRANCHES) for b in preprocess_branches]
        default_branches = [self.get_dict_key(b, self.BRANCHES) for b in self.defaults['branches']]
        self.params['branches'] = list(map(lambda name: self.BRANCHES[name],
                                           st.multiselect('Branches'+note,
                                                          available_branches,
                                                          default=default_branches)))
        self.validation_hash['not_empty_branches'].append(self.params['branches'])

        self.params['tb'] = st.checkbox('Output TensorBoard log files', value=self.defaults['tb'])

        st.markdown('## Training Options')
        # TODO make sure batch size is smaller than dataset size
        self.params['batch_size'] = st.number_input('Batch size', min_value=1, value=self.defaults['batch_size'])
        self.params['epochs'] = st.number_input('No. of training epochs', min_value=1, value=self.defaults['epochs'])
        self.params['early_stop'] = st.checkbox('Apply early stopping (patience 10, delta 0.01)', value=self.defaults['early_stop'])
        self.params['optimizer'] = self.OPTIMIZERS[st.selectbox(
            'Optimizer', list(self.OPTIMIZERS.keys()), index=self.get_dict_index(self.defaults['optimizer'], self.OPTIMIZERS))]
        if self.params['optimizer'] == 'sgd':
            # TODO move lr finder to hyperparam tuning, runs one epoch on small sample, does not need valid and test data
            lr_options = {'Use fixed learning rate (applies learning rate value throughout whole training)': 'fixed',
                          'Use learning rate scheduler (gradually decreasing from the learning rate value)': 'lr_scheduler',
                          # 'Use learning rate finder (beta)': 'lr_finder',
                          'Apply one cycle policy (uses the learning rate value as max)': 'one_cycle'}
            self.params['lr_optim'] = lr_options[st.radio('Learning rate options',
                                                          list(lr_options.keys()),
                                                          index=self.get_dict_index(self.defaults['lr_optim'], lr_options))]
        else:
            self.params['lr_optim'] = 'fixed'
        self.params['lr'] = st.number_input(
            'Learning rate', min_value=0.0001, max_value=0.1, value=self.defaults['lr'], step=0.0001, format='%.4f')

        # TODO change the logic when stateful design is enabled
        # self.branch_layers = {}
        # for branch in self.branches:
        #     self.branch_layers.update({branch: [list(LAYERS.keys())[0]]})
        #
        # for branch in self.branch_layers.keys():
        #     st.markdown(f'**{branch} branch**')
        #     for i, layer in enumerate(self.branch_layers[branch]):
        #         new_layer = st.selectbox('Layer', list(LAYERS.keys()), key=f'layer{branch}{i}')
        #         self.branch_layers[branch][i] = new_layer
        #     if st.button('Add layer', key=f'{branch}addlayer'):
        #         self.branch_layers[branch].append(
        #             st.selectbox('Layer', list(LAYERS.keys()), key=f'layer{branch}{len(self.branch_layers[branch])}'))
        #         print(self.branch_layers[branch])

        default_layer_args = {'batchnorm': False, 'dropout': 0.0, 'filters': 40, 'kernel': 4, 'units': 32, 'bidirect': True}

        st.markdown('## Network Architecture')

        self.params['no_branches_layers'] = st.number_input(
            'Number of layers in the branches:', min_value=0, value=self.defaults['no_branches_layers'], key=f'branches_no')
        self.params['branches_layers'] = self.defaults['branches_layers']

        for i, branch in enumerate(self.params['branches']):
            st.markdown(f'### {i+1}. {self.get_dict_key(branch, self.BRANCHES)} branch')
            self.params['branches_layers'][branch] = self.defaults['branches_layers'][branch][0:self.params['no_branches_layers']]
            for i in range(self.params['no_branches_layers']):
                if self.params_loaded and (i < len(self.defaults['branches_layers'][branch])):
                    default_args = default_layer_args
                    default_args.update(self.defaults['branches_layers'][branch][i]['args'])
                    layer = copy.deepcopy(self.defaults['branches_layers'][branch][i])
                    checkbox = True
                else:
                    default_args = default_layer_args
                    layer = {'name': 'Convolution layer', 'args': {'batchnorm': False, 'dropout': 0.0, 'filters': 40, 'kernel': 4}}
                    checkbox = False
                st.markdown(f'#### Layer {i + 1}')
                default_i = list(BRANCH_LAYERS.keys()).index(layer['name'])
                layer.update(dict(name=st.selectbox('Layer type', list(BRANCH_LAYERS.keys()), index=default_i, key=f'layer{branch}{i}')))
                layer = self.layer_options(layer, i, checkbox, default_args, branch)
                st.markdown('---')
                if len(self.params['branches_layers'][branch]) > i:
                    self.params['branches_layers'][branch][i] = layer
                else:
                    self.params['branches_layers'][branch].append(layer)

        st.markdown(f"### {len(self.params['branches'])+1}. Connected (after branches' concatenation)")
        self.params['common_layers'] = self.defaults['common_layers']
        self.params['no_common_layers'] = st.number_input(
            'Number of layers after concatenation of branches:', min_value=0, value=self.defaults['no_common_layers'], key=f'common_no')
        for i in range(self.params['no_common_layers']):
            if i == 0:
                allowed_layers = ['Dense layer', 'Convolution layer', 'GRU', 'LSTM']  # 'Locally Connected 1D layer',
            else:
                previous_layer = self.params['common_layers'][i-1]['name']
                if previous_layer in ('Convolution layer', 'GRU', 'LSTM'):  # 'Locally Connected 1D layer',
                    allowed_layers = ['Dense layer', previous_layer]
                elif previous_layer == 'Dense layer':
                    allowed_layers = ['Dense layer']

            if self.params_loaded and (i < len(self.defaults['common_layers'])) \
                    and self.defaults['common_layers'][i]['name'] in allowed_layers:
                default_args = default_layer_args
                default_args.update(self.defaults['common_layers'][i]['args'])
                layer = copy.deepcopy(self.defaults['common_layers'][i])
                checkbox = True
            else:
                default_args = default_layer_args
                layer = {'name': 'Dense layer', 'args': {'batchnorm': False, 'dropout': 0.0, 'units': 32}}
                checkbox = False
            default_i = list(COMMON_LAYERS.keys()).index(layer['name'])

            st.markdown(f'#### Layer {i + 1}')
            layer.update(dict(name=st.selectbox('Layer type', allowed_layers, index=default_i, key=f'common_layer{i}')))
            layer = self.layer_options(layer, i, checkbox, default_args)
            st.markdown('---')
            if len(self.params['common_layers']) > i:
                self.params['common_layers'][i] = layer
            else:
                self.params['common_layers'].append(layer)

        self.validate_and_run(self.validation_hash)

    # TODO adjust when stateful ops enabled
    def layer_options(self, layer, i, checkbox, defaults=None, branch=None):
        if st.checkbox('Show advanced options', value=checkbox, key=f'show{branch}{i}'):
            layer['args'].update({'batchnorm': st.checkbox(
                'Batch normalization', value=defaults['batchnorm'], key=f'batch{branch}{i}')})
            layer['args'].update({'dropout': st.slider(
                'Dropout rate', min_value=0.0, max_value=1.0, value=defaults['dropout'], key=f'do{branch}{i}', format='%.2f')})
            if layer['name'] in ['Convolution layer', 'Locally Connected 1D layer']:
                layer['args'].update({'filters': st.number_input('Number of filters:', min_value=1, value=
                defaults['filters'], key=f'filters{branch}{i}')})
                layer['args'].update({'kernel': st.number_input('Kernel size:', min_value=1, value=
                defaults['kernel'], key=f'kernel{branch}{i}')})
            elif layer['name'] in ['Dense layer', 'GRU', 'LSTM']:
                layer['args'].update({'units': st.number_input('Number of units:', min_value=1, value=
                defaults['units'], key=f'units{branch}{i}')})
                if layer['name'] in ['GRU', 'LSTM']:
                    layer['args'].update({'bidirect': st.checkbox(
                        'Bidirectional', value=defaults['bidirect'], key=f'bidirect{branch}{i}')})

        return layer

    @staticmethod
    def parse_data(dataset_files, branches, label_encoding):
        dictionary = {}
        for file in dataset_files:
            dataset = Dataset.load_from_file(file)
            if dataset.category == 'blackbox': continue
            dictionary.update({dataset.category: {}})

            values = dataset.encode_branches(dataset, branches)
            dictionary[dataset.category].update({'values': values})
            dictionary[dataset.category].update({'labels': dataset.labels(encoding=label_encoding)})

        return [dictionary['train']['values'], dictionary['validation']['values'], dictionary['test']['values'],
                dictionary['train']['labels'], dictionary['validation']['labels'], dictionary['test']['labels']]

    @staticmethod
    def get_shapes(data, branches):
        shapes = {}

        if len(branches) == 1:
            shapes.update({branches[0]: data.shape})
        else:
            # Assuming fixed order of the branches
            for i, branch in enumerate(branches):
                shapes.update({branch: data[i].shape})

        return shapes

    def run(self):
        status = st.empty()
        status.text('Initializing network...')

        candidate_files = f.list_files_in_dir(self.params['input_folder'], 'zip')
        categories = ['train', 'validation', 'test', 'blackbox']
        dataset_files = [file for file in candidate_files if any(category in os.path.basename(file) for category in categories)]

        if self.previous_param_file:
            with open(self.previous_param_file, 'r') as file:
                previous_params = yaml.safe_load(file)
                klasses = previous_params['Preprocess']['klasses']
                klass_alphabet = {klass: i for i, klass in enumerate(klasses)}
                encoded_labels = seq.onehot_encode_alphabet(klass_alphabet)
        else:
            raise UserInputError('Could not read class labels from parameters.yaml file).')
        train_x, valid_x, test_x, train_y, valid_y, test_y = self.parse_data(dataset_files, self.params['branches'], encoded_labels)
        branch_shapes = self.get_shapes(train_x, self.params['branches'])

        self.params['train_dir'] = os.path.join(self.params['output_folder'], 'training',
                                 f'{str(datetime.datetime.now().strftime("%Y%m%d-%H%M"))}')
        self.ensure_dir(self.params['train_dir'])

        model = ModelBuilder(self.params['branches'], encoded_labels, branch_shapes, self.params['branches_layers'], self.params['common_layers']).build_model()
        optimizer = self.create_optimizer(self.params['optimizer'], self.params['lr'])
        model.compile(
            optimizer=optimizer,
            loss=['categorical_crossentropy'],
            metrics=[list(self.TRAIN_METRICS.keys())])

        # Train the model
        status.text('Training the network...')

        progress_bar = st.progress(0)
        progress_status = st.empty()
        chart = st.altair_chart(self.initialize_altair_chart(), use_container_width=True)

        callbacks = self.create_callbacks(
            self.params['train_dir'], self.params['lr_optim'], self.params['tb'], self.params['epochs'], progress_bar, progress_status, chart, self.params['early_stop'],
            self.params['lr'], branch_shapes[self.params['branches'][0]][0])

        history = self.train(model, self.params['epochs'], self.params['batch_size'], callbacks, train_x, valid_x, train_y, valid_y).history
        # if self.params['lr_optim'] == 'lr_finder': self.params['epochs'] = 1
        # if self.params['lr_optim'] == 'lr_finder': LRFinder.plot_schedule_from_file(self.params['train_dir'])

        if self.params['early_stop']:
            early_epochs = [callback for callback in callbacks if type(callback) == EarlyStopping][0]
            if early_epochs and early_epochs.stopped_epoch != 0:
                self.params['epochs'] = early_epochs.stopped_epoch

        self.log_train_val_metrics(history, self.params)

        # Plot training metrics
        # if self.params['lr_optim'] != 'lr_finder':
        train_plot_dir = os.path.join(self.params['train_dir'], 'plots', 'training_metrics')
        self.ensure_dir(train_plot_dir)
        for metric, title in self.TRAIN_METRICS.items():
            metric_name = metric if metric == 'accuracy' else metric.name
            self.plot_training_metric(history, metric_name, title, train_plot_dir)
        self.plot_training_metric(history, 'loss', 'Loss', train_plot_dir)

        try:
            tf.keras.utils.plot_model(model, to_file=f"{self.params['train_dir']}/model.png", show_shapes=True, dpi=300)
        except:
            logger.warning('Did not exported model plot due to an error.')

        model_json = model.to_json()
        with open(f"{self.params['train_dir']}/model.json", 'w') as json_file:
            json_file.write(model_json)

        status.text('Evaluating model...')
        eval_plot_dir = os.path.join(self.params['train_dir'], 'plots', 'evaluation_metrics')
        self.ensure_dir(eval_plot_dir)
        self.evaluate_model(encoded_labels, model, test_x, test_y, self.params, eval_plot_dir)

        # Prepare tsv row content
        header = self.train_header()
        row = self.train_row(self.params)
        if 'Preprocess' in previous_params.keys():
            # Parameters missing in older versions of the code
            novel_params = {'win_place': 'rand'}  # It's always been 'random' for the previous versions
            parameters = novel_params
            parameters.update(previous_params['Preprocess'])
            header += f'{self.preprocess_header()}\n'
            row += f"{self.preprocess_row(parameters)}\n"
        else:
            header += '\n'
            row += '\n'

        self.finalize_run(logger, self.params['train_dir'], self.params, header, row, previous_param_file=self.previous_param_file)
        status.text('Finished!')
        logger.info('Finished!')

    @staticmethod
    def step_decay_schedule(initial_lr, drop=0.5, epochs_drop=10.0):
        def schedule(epoch):
            return initial_lr * math.pow(drop, math.floor(epoch + 1) / epochs_drop)
        return LearningRateScheduler(schedule)

    @staticmethod
    def create_callbacks(out_dir, lr_optim, tb, epochs, progress_bar, progress_status, chart, early_stop, lr, sample):
        mcp = ModelCheckpoint(filepath=out_dir + '/model.hdf5',
                              verbose=0,
                              save_best_only=True)

        csv_logger = CSVLogger(out_dir + '/log.csv',
                               append=True,
                               separator='\t')

        progress = ProgressMonitor(epochs, progress_bar, progress_status, chart, Train.TRAIN_METRICS.keys())

        callbacks = [mcp, csv_logger, progress]

        if early_stop:
            earlystopper = EarlyStopping(monitor='val_loss',
                                         patience=10,
                                         min_delta=0.01,
                                         verbose=1,
                                         mode='auto')
            callbacks.append(earlystopper)

        if lr_optim != 'fixed':
            # if lr_optim == 'lr_finder':
            #     callbacks.append(LRFinder(num_samples=sample,
            #                               batch_size=sample//30,
            #                               minimum_lr=1e-5,
            #                               maximum_lr=1e0,
            #                               lr_scale='exp',
            #                               save_dir=out_dir))
            if lr_optim == 'one_cycle':
                callbacks.append(OneCycleLR(max_lr=lr,
                                            end_percentage=0.1,
                                            scale_percentage=None,
                                            maximum_momentum=0.95,
                                            minimum_momentum=0.85,
                                            verbose=True))
            elif lr_optim == 'lr_scheduler':
                callbacks.append(Train.step_decay_schedule(initial_lr=lr))

        if tb:
            tb_dir = os.path.join(out_dir, 'tb')
            callbacks.append(TensorBoard(log_dir=tb_dir, histogram_freq=1, profile_batch=3))

        return callbacks

    @staticmethod
    def create_optimizer(chosen, learning_rate):
        if chosen == 'sgd':
            optimizer = SGD(
                lr=learning_rate,
                momentum=0.9,
                nesterov=True)
        elif chosen == 'rmsprop':
            optimizer = RMSprop(
                lr=learning_rate
            )
        elif chosen == 'adam':
            optimizer = Adam(
                lr=learning_rate,
            )

        return optimizer

    @staticmethod
    def train(model, epochs, batch_size, callbacks, train_x, valid_x, train_y, valid_y):
        history = model.fit(
            train_x,
            train_y,
            batch_size=batch_size,
            epochs=epochs,
            verbose=1,
            validation_data=(valid_x, valid_y),
            callbacks=callbacks)

        return history

    @staticmethod
    def log_train_val_metrics(history, params):
        st.text('Final metric values:')

        params['train_loss'] = str(round(history['loss'][-1], 4))
        params['train_acc'] = str(round(history['accuracy'][-1], 4))
        # params['train_auc'] = str(round(history['auc'][-1], 4))

        logger.info('Training loss: ' + params['train_loss'])
        logger.info('Training accuracy: ' + params['train_acc'])
        # logger.info('Training AUC: ' + params['train_auc'])

        params['val_loss'] = str(round(history['val_loss'][-1], 4))
        params['val_acc'] = str(round(history['val_accuracy'][-1], 4))
        # params['val_auc'] = str(round(history['val_auc'][-1], 4))

        logger.info('Validation loss: ' + params['val_loss'])
        logger.info('Validation accuracy: ' + params['val_acc'])
        # logger.info('Validation AUC: ' + params['val_auc'])

        st.text(f"Final achieved training loss: {params['train_loss']}\n"
                f"Final achieved training accuracy: {params['train_acc']}\n"
                # f"Final achieved training AUC: {params['train_auc']}\n"
                "\n"
                f"Final achieved validation loss: {params['val_loss']} \n"
                f"Final achieved validation accuracy: {params['val_acc']} \n")
                # f"Final achieved validation AUC: {params['val_auc']} \n\n")

    @staticmethod
    def plot_training_metric(history, metric, title, out_dir):
        val_metric = f'val_{metric}'
        file_path = os.path.join(out_dir, f'{metric}')

        plt.plot(history[metric], label='Training')
        plt.plot(history[val_metric], label='Validation')

        if metric == 'accuracy' or metric == 'auc' or metric == 'precision' or metric == 'recall':
            plt.ylim(0.0, 1.05)
        elif metric == 'loss':
            plt.ylim(0.0, max(max(history[metric]), max(history[val_metric])) + 0.05)

        plt.title(title)
        plt.ylabel(title)
        plt.xlabel('Epoch')
        plt.legend(loc='lower right')
        plt.savefig(file_path, format='png', dpi=300)
        plt.clf()

    @staticmethod
    def initialize_altair_chart():
        source = pd.DataFrame([], columns=['Metric', 'Metric value', 'Epoch'])
        nearest = alt.selection(type='single', nearest=True, on='mouseover', fields=['Epoch'], empty='none')
        line = alt.Chart(source).mark_line().encode(x='Epoch:Q', y='Metric value:Q', color='Metric:N')
        selectors = alt.Chart(source).mark_point().encode(x='Epoch:Q', opacity=alt.value(0)).add_selection(nearest)
        points = line.mark_point().encode(opacity=alt.condition(nearest, alt.value(1), alt.value(0)))
        text = line.mark_text(align='left', dx=5, dy=-5).encode(text=alt.condition(nearest, 'Metric value:Q', alt.value(' ')))
        rules = alt.Chart(source).mark_rule(color='gray').encode(x='Epoch:Q').transform_filter(nearest)
        alt_chart = alt.layer(line, selectors, points, rules, text)
        return alt_chart

    @staticmethod
    def default_params():
        return {'batch_size': 256,
                'branches': [],
                'branches_layers': {'seq': [], 'fold': [], 'cons': []},
                'common_layers': [],
                'early_stop': True,
                'epochs': 100,
                'input_folder': '',
                'lr': 0.005,
                'lr_optim': 'fixed',
                'no_branches_layers': 1,
                'no_common_layers': 1,
                'optimizer': 'sgd',
                'output_folder': os.path.join(os.path.expanduser('~'), 'enngene_output'),
                'tb': True}
```
