## Additional file 1 for "ENNGene: an Easy Neural Network model building tool for Genomics": dataset.py

```
import _io
import datetime
import logging
import numpy as np
import os
import pandas as pd
import re
import streamlit as st
import subprocess
import tempfile

from functools import reduce
from zipfile import ZipFile, ZIP_DEFLATED

from .exceptions import UserInputError, ProcessError
from . import file_utils as f
from . import sequence as seq

logger = logging.getLogger('root')


class Dataset:

    @classmethod
    @st.cache(hash_funcs={_io.TextIOWrapper: lambda _: None}, suppress_st_warning=True)
    def load_and_cache(cls, file_path):
        with st.spinner('Reading in the mapped file. May take up to few minutes...'):
            full_dataset = cls.load_from_file(file_path)
            klasses = list(full_dataset.df['klass'].unique())
            valid_chromosomes = list(full_dataset.df['chrom_name'].unique())
            klass_sizes = {}
            for klass in klasses:
                count = len(full_dataset.df[(full_dataset.df['klass'] == klass)])
                klass_sizes.update({klass: count})
        return klasses, valid_chromosomes, full_dataset.branches, klass_sizes

    @classmethod
    def load_from_file(cls, file_path):
        name = os.path.basename(file_path).replace('.tsv.zip', '')
        df = pd.read_csv(file_path, sep='\t', header=0)
        branches = [col for col in df.columns if col not in ['chrom_name', 'seq_start', 'seq_end', 'strand_sign', 'klass']]
        category = name if (name in ['train', 'test', 'validation', 'blackbox']) else None

        return cls(branches=branches, category=category, df=df)

    @classmethod
    def split_by_chr(cls, dataset, chrs_by_category):
        split_datasets = set()
        for category, chr_list in chrs_by_category.items():
            df = dataset.df[dataset.df['chrom_name'].isin(chr_list)]
            split_datasets.add(
                Dataset(klass=dataset.klass, branches=dataset.branches, category=category, df=df))

        return split_datasets

    @classmethod
    def split_random(cls, dataset, ratio):
        # so far the categories are fixed, not sure if there would be need for custom categories
        ratio_list = ratio.split(':')
        ratio_list = [float(x) for x in ratio_list]
        dataset_size = dataset.df.shape[0]
        total = sum(ratio_list)
        np.random.shuffle(dataset.df.values)
        split_datasets = set()

        validation_size = int(dataset_size * ratio_list[1] / total)
        test_size = (int(dataset_size * ratio_list[2] / total))
        blackbox_size = (int(dataset_size * ratio_list[3] / total))
        dfs = {'validation': dataset.df[:validation_size]}
        dfs.update({'test': dataset.df[validation_size:(validation_size + test_size)]})
        dfs.update({'blackbox': dataset.df[(validation_size + test_size):(validation_size + test_size + blackbox_size)]})
        dfs.update({'train': dataset.df[(validation_size + test_size + blackbox_size):]})

        for category, df in dfs.items():
            split_datasets.add(
                Dataset(klass=dataset.klass, branches=dataset.branches, category=category, df=df))

        return split_datasets

    @classmethod
    def merge_dataframes(cls, dataset_list):
        dataframes = [dataset.df for dataset in dataset_list]
        merged_df = reduce(lambda left, right: pd.merge(left, right, how='outer'), dataframes)
        return merged_df

    @classmethod
    def merge_by_category(cls, set_of_datasets):
        datasets_by_category = {}
        for dataset in set_of_datasets:
            if dataset.category not in datasets_by_category.keys():
                datasets_by_category.update({dataset.category: []})
            datasets_by_category[dataset.category].append(dataset)

        merged_datasets = set()
        for category, datasets in datasets_by_category.items():
            branches = datasets[0].branches
            merged_df = cls.merge_dataframes(datasets)
            merged_datasets.add(cls(branches=branches, category=category, df=merged_df))

        return merged_datasets

    def __init__(self, klass=None, branches=None, category=None, win=None, win_place=None,
                 bed_file=None, fasta_file=None, text_input=None, df=None):
        self.branches = branches  # list of seq, cons or fold branches
        self.klass = klass  # e.g. positive or negative
        self.category = category  # predict, or train, validation, test or blackbox for separated datasets
        self.df = df if df is not None else pd.DataFrame()

        evaluation = category == 'eval'
        if bed_file:
            input_type = 'bed'
            self.df = self.read_in_bed(bed_file, evaluation)
        elif fasta_file:
            input_type = 'fasta'
            self.df = self.read_in_fasta(fasta_file, evaluation)
        elif text_input:
            input_type = 'fasta'
            self.df = self.read_in_text(text_input)

        if win:
            self.df = Dataset.apply_window(self.df, win, win_place, input_type)

    def read_in_bed(self, bed_file, evaluation=False):
        df = pd.read_csv(bed_file, sep='\t', header=None)

        if len(df.columns) == 3 or (evaluation and len(df.columns) == 4):
            if evaluation:
                df.columns = ['klass', 'chrom_name', 'seq_start', 'seq_end']
            else:
                df.columns = ['chrom_name', 'seq_start', 'seq_end']
            df['strand_sign'] = '.'; df['name'] = ''; df['score'] = np.nan
            df[['seq_start', 'seq_end']] = df[['seq_start', 'seq_end']].astype(int)
            df['chrom_name'] = df['chrom_name'].astype(str)
        elif len(df.columns) >= 6:
            if evaluation:
                df = df[[0, 1, 2, 3, 4, 5, 6]]
                df.columns = ['klass', 'chrom_name', 'seq_start', 'seq_end', 'name', 'score', 'strand_sign']
            else:
                df = df[[0, 1, 2, 3, 4, 5]]
                df.columns = ['chrom_name', 'seq_start', 'seq_end', 'name', 'score', 'strand_sign']
            df[['seq_start', 'seq_end', 'score']] = df[['seq_start', 'seq_end', 'score']].astype(int, errors='ignore')
            df[['chrom_name', 'name', 'strand_sign']] = df[['chrom_name', 'name', 'strand_sign']].astype(str, errors='ignore')
        else:
            raise UserInputError('Invalid format of a .bed file.')
        if self.klass and not evaluation:
            df['klass'] = self.klass

        if evaluation:
            df['klass'] = df['klass'].astype(str)

        # def check_valid(row):
        #     return row if seq.is_valid_chr(row['chrom_name']) else None
        # df = df.apply(check_valid, axis=1).dropna()

        return df

    def reduce(self, ratio):
        np.random.shuffle(self.df.values)
        if ratio <= 1:
            # handle as a ratio
            last = int(self.df.shape[0] * ratio)
        elif ratio < len(self.df.values):
            # handle as a final size
            last = int(ratio)
        elif ratio >= len(self.df.values):
            # keep the full dataset
            last = len(self.df.values)-1
        self.df = self.df[:last]
        return self

    def labels(self, encoding=None):
        labels = self.df['klass']
        if encoding:
            encoded_labels = [seq.translate(item, encoding) for item in labels]
            return np.array(encoded_labels)
        else:
            return np.array(labels)

    def map_to_branches(self, references, strand, outfile_path, status, predict=False, ncpu=1):
        mapped = False
        # map seq branch first so that we can replace the df without loosing anny information
        branches = sorted(self.branches, key=lambda x: (x != 'seq', x != 'fold'))

        for branch in branches:
            if branch == 'seq':
                if 'seq' not in self.df.columns:
                    status.text(f'Mapping intervals to the fasta reference...')
                    self.df = self.map_to_fasta(self.df, branch, strand, references[branch], predict)
                mapped = True
            elif branch == 'cons':
                status.text(f'Mapping intervals to the wig reference... \n'
                            f'Note: This is rather slow process, it may take a while.')
                self.df = Dataset.map_to_wig(branch, self.df, references[branch])
            elif branch == 'fold':
                status.text(f'Folding the sequences...')
                if mapped:
                    self.df['fold'] = self.df['seq']  # already finished above
                else:
                    self.df = self.map_to_fasta(self.df, branch, strand, references[branch], predict)
                # TODO for prediction/evaluation key_cols = seq pravdepodobne
                key_cols = ['chrom_name', 'seq_start', 'seq_end', 'strand_sign']

                seq_branch = 'seq' in branches
                self.df = self.fold_branch(self.df, key_cols, seq_branch, ncpu)

        self.df.dropna(subset=branches, inplace=True)
        self.save_to_file(outfile_path, ignore_cols=['name', 'score'], do_zip=True)
        return self

    def sort_datapoints(self):
        self.df = self.df.sort_values(by=['chrom_name', 'seq_start'])
        self.df.reset_index(drop=True, inplace=True)
        return self

    def save_to_file(self, outfile_path, do_zip=False, ignore_cols=None):
        to_export = [col for col in self.df.columns if col not in ignore_cols] if ignore_cols else self.df.columns
        self.df.to_csv(outfile_path, sep='\t', columns=to_export, index=False)

        if do_zip:
            logger.info(f'Compressing dataset file... {outfile_path}')
            zipped = ZipFile(f'{outfile_path}.zip', 'w')
            zipped.write(outfile_path, os.path.basename(outfile_path), compress_type=ZIP_DEFLATED)
            zipped.close()
            os.remove(outfile_path)

    def encode_col(self, col, new_col, encoding):
        def map(row):
            sequence = row[col]
            encoded = Dataset.encode_sequence(sequence, encoding)
            row[new_col] = encoded
            return row

        self.df = self.df.apply(map, axis=1)
        return self

    @staticmethod
    def encode_branches(dataset, branches):
        old_values = '|' in dataset.df[branches[0]][0]
        values = []
        if old_values:  # TODO remove eventually
            for branch in branches:
                value = []
                for string in dataset.df[branch]:
                    value.append(dataset.sequence_from_string(string))
                values.append(np.array(value))
        else:
            for branch in branches:
                if branch in ['seq', 'fold']:
                    alphabet = seq.ALPHABET if branch == 'seq' else seq.FOLDING
                    branch_encoding = seq.onehot_encode_alphabet(alphabet)
                    dataset.encode_col(branch, f'{branch}_encoded', branch_encoding)
                    np_col = [np.array(value) for value in dataset.df[f'{branch}_encoded']]
                    values.append(np.array(np_col))
                elif branch == 'cons':
                    dataset.df[branch] = dataset.df.apply(lambda x: [float(n) for n in x[branch].split(',')], axis=1)
                    np_col = [np.array([np.array([float(v)]) for v in value]) for value in dataset.df[branch].to_list()]
                    values.append(np.array(np_col))
        # Do not return data in an extra array if there's only one branch
        if len(values) == 1:
            values = values[0]

        return values

    @staticmethod
    def read_in_fasta(fasta_file, evaluation=False):
        df = pd.DataFrame()

        with open(fasta_file, 'r') as file:
            header = None
            sequence = ""
            for line in file:
                if '>' in line:
                    # Save finished previous key value pair (unless it's the first iteration)
                    if header:
                        new_row = pd.DataFrame([[header, sequence, klass]]) if evaluation else pd.DataFrame([[header, sequence]])
                        df = df.append(new_row)
                    if evaluation:
                        parts = line.strip().strip('>').split()
                        klass = parts[-1]
                        header = ''.join(parts[0:-1])
                    else:
                        header = line.strip().strip('>')
                    sequence = ""
                else:
                    if header:
                        sequence += line.strip()
                    else:
                        raise UserInputError("Provided reference file does not start with '>' fasta identifier.")
            # Save the last key value pair
            if header and sequence:
                new_row = pd.DataFrame([[header, sequence, klass]]) if evaluation else pd.DataFrame([[header, sequence]])
                df = df.append(new_row)

        if evaluation:
            df.columns = ['header', 'seq', 'klass']
        else:
            df.columns = ['header', 'seq']
        return df

    @staticmethod
    def read_in_text(text):
        df = pd.DataFrame()
        seq_series = pd.Series(text.strip().split('\n'))
        df['seq'] = seq_series
        return df

    @staticmethod
    def map_to_fasta(df, branch, strand, fasta, predict):
        df[branch] = None
        tmp_dir = tempfile.gettempdir()
        tmp_df = df
        has_klass = 'klass' in df.columns

        # we need to preserve klass name in Preprocessing, while bedtools can only keep name column
        if has_klass: df['name'] = df['klass']

        if strand:
            key_cols = ['chrom_name', 'seq_start', 'seq_end', 'name', 'score', 'strand_sign']
            tmp_df['name'] = tmp_df['name'].replace(r'^\s*$', 'x', regex=True)
            tmp_df['strand_sign'] = tmp_df['strand_sign'].replace(r'^\s*$', '+', regex=True)
        else:
            key_cols = ['chrom_name', 'seq_start', 'seq_end']
        bed_file = Dataset.dataframe_to_bed(tmp_df, key_cols, tmp_dir, f'bed_{str(datetime.datetime.now().strftime("%Y%m%d-%H%M%S"))}')

        tmp_file = os.path.join(tmp_dir, f'mapped_{str(datetime.datetime.now().strftime("%Y%m%d-%H%M%S"))}')
        err_file = os.path.join(tmp_dir, f'err_{str(datetime.datetime.now().strftime("%Y%m%d-%H%M%S"))}')
        out_file = open(tmp_file, 'w+')
        out_err = open(err_file, 'w+')

        try:
            if strand:
                subprocess.run(['bedtools', 'getfasta', '-s', '-name', '-tab', '-fi', fasta, '-bed', bed_file],
                               stdout=out_file, stderr=out_err, check=True)
            else:
                subprocess.run(['bedtools', 'getfasta', '-name', '-tab', '-fi', fasta, '-bed', bed_file],
                               stdout=out_file, stderr=out_err, check=True)
        except Exception:
            raise ProcessError('There was an error during mapping intervals to the fasta reference. '
                               f'Please check bedtools error report: {err_file}.')

        bedtools_df = pd.read_csv(tmp_file, sep='\t', header=None)

        if len(bedtools_df) > len(df):
            # For some reason sometimes returns first sequence multiple times
            dif = len(bedtools_df) - len(df)
            bedtools_df = bedtools_df[dif:]

        if len(df) == len(bedtools_df):
            logger.info(f'Sequence: Mapped 100% of the intervals.')
            df[branch] = bedtools_df.iloc[:, 1]
            new_df = df
        else:
            logger.info(f'Sequence: Mapped {round((len(bedtools_df)/len(df)*100), 1)}% intervals ({len(bedtools_df)} out of {len(df)})')
            # It is too slow to cherrypick the mapped into to original df, so we replace it by the smaller one instead
            bedtools_df.columns = ['header', branch]
            bedtools_df['chrom_name'] = ''; bedtools_df['seq_start'] = np.nan; bedtools_df['seq_end'] = np.nan; bedtools_df['klass'] = None

            def parse_header(row):
                header = row['header']
                klass, rest = header.split('::')
                chr, rest = rest.split(':')
                row['chrom_name'] = chr
                row['seq_start'] = int(rest.split('-')[0])
                rest = rest.split('-')[1]
                row['seq_end'] = int(rest.split('(')[0])
                match = re.search(r'\((.)\)$', rest)
                strand = match[1] if match else ''
                row['strand_sign'] = strand
                if has_klass: row['klass'] = klass
                return row

            new_df = bedtools_df.apply(parse_header, axis=1)
            new_df = new_df.drop('header', 1)
            reordered_cols = ['chrom_name', 'seq_start', 'seq_end', 'strand_sign', 'klass', branch]
            new_df = new_df[reordered_cols]

        new_df.reset_index(inplace=True, drop=True)
        return new_df

    @staticmethod
    def encode_sequence(sequence, encoding):
        new_sequence = [seq.translate(item, encoding) for item in sequence]
        return new_sequence

    @staticmethod
    def sequence_from_string(string):
        # TODO ideally make more explicit, maybe split the method
        #  (currently expects floats everywhere, can ever be soemthing else?)
        parts = string.strip().split('|')
        sequence = []

        for part in parts:
            if ',' in part:  # expects one hot encoded sequence, thus branches seq and fold
                subparts = part.strip().split(',')
                new_part = []
                for subpart in subparts:
                    new_part.append(float(subpart))
                sequence.append(np.array(new_part))
            else:  # expects not encoded sequence, thus branch cons
                sequence.append(np.array([float(part)]))

        return np.array(sequence)

    @staticmethod
    def map_to_wig(branch, df, ref_folder):
        not_found_chrs = set()
        chrom_files = f.list_files_in_dir(ref_folder, 'wig')

        current_file = None
        zipped = None
        current_chr = None
        current_header = {}
        parsed_line = {}
        df_copy = df.copy()

        for i, row in df_copy.iterrows():
            score = []

            if row['chrom_name'] and row['chrom_name'] == current_chr:
                result = Dataset.map_datapoint_to_wig(
                    score, zipped, row['seq_start'], row['seq_end'], current_file, current_header, parsed_line)
                if result:
                    score, current_header, parsed_line = result
                else:
                    # Covering the case when we reach end of reference file while still having some samples with current_chr not mapped
                    not_found_chrs.add(current_chr)
                    continue
            elif row['chrom_name'] in not_found_chrs:
                continue
            else:
                # When reading from new reference file
                files = list(filter(lambda f: f"{row['chrom_name']}." in os.path.basename(f), chrom_files))
                if len(files) == 1:
                    if current_file:
                        current_file.close()
                    current_chr = row['chrom_name']

                    current_file, zipped = f.unzip_if_zipped(files[0])

                    line = f.read_decoded_line(current_file, zipped)
                    # Expecting first line of the file to be a header
                    if 'chrom' in line:
                        current_header = seq.parse_wig_header(line)
                    else:
                        raise UserInputError('File not starting with a proper wig header.')
                    result = Dataset.map_datapoint_to_wig(
                        score, zipped, row['seq_start'], row['seq_end'], current_file, current_header,
                        parsed_line)
                    if result:
                        score, current_header, parsed_line = result
                    else:  # should not happen here, as it's beginning of the file
                        not_found_chrs.add(current_chr)
                        continue
                else:
                    not_found_chrs.add(row['chrom_name'])
                    if len(files) == 0:
                        # TODO or rather raise an exception to let user fix it?
                        # Anyway, let the user know if none were found, thus the path given is wrong (currently it looks like it went through)
                        logger.warning(
                            f"Didn\'t find appropriate conservation file for {row['chrom_name']}, skipping the chromosome.")
                    else:  # len(files) > 1
                        logger.warning(f"Found multiple conservation files for {row['chrom_name']}, skipping the chromosome.")
                    continue

            if score and len(score) == (row['seq_end'] - row['seq_start']):
                # Score may be fully or partially missing if the coordinates are not part of the reference
                str_score = [str(num) for num in score]
                df.loc[i, branch] = ','.join(str_score)

        unmapped = df[branch].isna().sum()
        logger.info(f'Conservation score: mapped {round(((len(df)-unmapped)/len(df)*100), 1)}% of the intervals ({(len(df)-unmapped)} out of {len(df)}).')
        return df

    @staticmethod
    def map_datapoint_to_wig(score, zipped, dp_start, dp_end, current_file, current_header, parsed_line):
        # TODO check the logic of passing on the parsed line
        # FIXME not covered: when the end of ref file is reached while there are still some samples from that file
        # unmapped - it reads empty lines - break with first empty line read as expecting it to be EOF? and move to
        # next chromosome somehow

        line = f.read_decoded_line(current_file, zipped)
        new_score = []
        if not line: return None

        if 'chrom' in line:
            current_header = seq.parse_wig_header(line)
            result = Dataset.map_datapoint_to_wig(score, zipped, dp_start, dp_end, current_file, current_header, parsed_line)
            if result:
                new_score, current_header, parsed_line = result
            else:
                return None
        else:
            if dp_start < current_header['start']:
                # Missed beginning of the datapoint while reading through the reference (should not happen)
                pass
            else:
                current_header, parsed_line = seq.parse_wig_line(line, current_header)
                while dp_start not in parsed_line.keys():
                    line = f.read_decoded_line(current_file, zipped)
                    if not line: return None
                    if 'chrom' in line:
                        current_header = seq.parse_wig_header(line)
                    else:
                        current_header, parsed_line = seq.parse_wig_line(line, current_header)
                    if current_header['start'] > dp_end:
                        # Did not find datapoint coordinates in reference file, might happen
                        break
                else:
                    for i in range(0, (dp_end - dp_start)):
                        coord = dp_start + i
                        if coord in parsed_line.keys():
                            score.append(parsed_line[coord])
                        else:
                            line = f.read_decoded_line(current_file, zipped)
                            if not line: return None
                            if 'chrom' in line:
                                current_header = seq.parse_wig_header(line)
                                line = f.read_decoded_line(current_file, zipped)
                                if not line: return None
                            current_header, parsed_line = seq.parse_wig_line(line, current_header)
                            try:
                                score.append(parsed_line[coord])
                            except:
                                # Lost the score in the reference in the middle of a datapoint, might happen
                                break

        return [score + new_score, current_header, parsed_line]

    @staticmethod
    def fold_branch(df, key_cols, seq_branch=True, ncpu=1):
        tmp_dir = tempfile.gettempdir()
        original_length = df.shape[0]
        has_klass = 'klass' in df.columns
        if has_klass: key_cols += ['klass']
        fasta_file = Dataset.dataframe_to_fasta(df, 'fold', key_cols, tmp_dir, f'df_{str(datetime.datetime.now().strftime("%Y%m%d-%H%M%S"))}')

        out_path = os.path.join(tmp_dir, f'folded_df_{str(datetime.datetime.now().strftime("%Y%m%d-%H%M%S"))}')
        out_file = open(out_path, 'w+')

        subprocess.run(['RNAfold', '--verbose', '--noPS', f'--jobs={ncpu}', fasta_file], stdout=out_file, check=True)

        folded_df = pd.read_csv(out_path, header=None, sep='\t')
        folded_df.reset_index(inplace=True, drop=True)  # ensure ordered index

        if len(folded_df) > (original_length*3):
            # For some reason sometimes returns first sequence multiple times
            dif = len(folded_df) - (original_length*3)
            new = round(dif/3)*3
            folded_df = folded_df[new:]
            folded_df.reset_index(inplace=True, drop=True)

        folded_len = (len(folded_df) / 3)
        folded_df.columns = ['output']

        def parse_folding(row):
            folding = row['output']
            # line format: '.... (0.00)'
            value = folding.split(' ')[0].strip()
            row['fold'] = value
            return row

        def parse_all(row):
            header = row['header']
            header_parts = header.split('_')
            row['chrom_name'] = header_parts[0][1:]
            row['seq_start'] = header_parts[1]
            row['seq_end'] = header_parts[2]
            row['strand_sign'] = header_parts[3]
            if has_klass:
                klass = '_'.join(header_parts[4:])
                row['klass'] = klass

            row['seq'] = row['seq'].replace('U', 'T')

            folding = row['fold']
            # line format: '.... (0.00)'
            value = folding.split(' ')[0].strip()
            row['fold'] = value
            return row

        # The order should remain the same as long as --unordered is not set to True
        if folded_len == original_length:
            logger.info('Secondary structure: Folded 100% sequences.')
            # We're interested only in each third line in the output file (there are 3 lines per one input sequence)
            folded_df['fold'] = None
            folded_df = folded_df[(folded_df.index+1) % 3 == 0]
            folded_df = folded_df.apply(parse_folding, axis=1)
            folded_df.reset_index(inplace=True, drop=True)
            new_df = df
            new_df.reset_index(inplace=True, drop=True)
            new_df['fold'] = folded_df['fold']
        else:
            logger.info(f'Secondary structure: Folded {round(((len(folded_df)/3)/original_length*100), 1)}% of sequences ({len(folded_df)/3} out of {original_length}.')
            df1 = folded_df.iloc[::3].reset_index(drop=True)
            df2 = folded_df.iloc[1::3].reset_index(drop=True)
            df3 = folded_df.iloc[2::3].reset_index(drop=True)
            new_df = pd.concat([df1, df2, df3], ignore_index=True, axis=1)
            new_df.columns = ['header', 'seq', 'fold']

            new_df['chrom_name'] = ''; new_df['seq_start'] = np.nan; new_df['seq_end'] = np.nan; new_df['klass'] = None
            new_df = new_df.apply(parse_all, axis=1)
            new_df = new_df.drop('header', 1)
            reordered_cols = ['chrom_name', 'seq_start', 'seq_end', 'strand_sign', 'klass', 'seq', 'fold']
            new_df = new_df[reordered_cols]
            if not seq_branch:
                new_df = new_df.drop('seq', 1)
            new_df.reset_index(inplace=True, drop=True)
            new_df.sort_values(by=['chrom_name', 'seq_start'])

        return new_df

    @staticmethod
    def dataframe_to_bed(df, key_cols, path, name):
        path_to_bed = os.path.join(path, (name + ".bed"))
        content = []

        def to_bed(row):
            cols = ""
            for col in key_cols:
                cols += str(row[col])
                cols += '\t'
            line = cols.strip() + '\n'
            content.append(line)
            return row

        df.apply(to_bed, axis=1)
        f.write(path_to_bed, ''.join(content).strip())
        return path_to_bed

    @staticmethod
    def dataframe_to_fasta(df, branch, key_cols, path, name):
        path_to_fasta = os.path.join(path, (name + ".fa"))
        content = []

        def to_fasta(row):
            key = []
            for col in key_cols:
                key.append(str(row[col]))
            line1 = ">" + '_'.join(key) + "\n"
            line2 = str(row[branch]) + "\n"
            content.append(line1)
            content.append(line2)
            return row

        df.apply(to_fasta, axis=1)
        f.write(path_to_fasta, ''.join(content).strip())
        return path_to_fasta

    @staticmethod
    def apply_window(df, window_size, win_place='rand', type='bed'):
        def bed_window(row):
            length = row['seq_end'] - row['seq_start']
            if length > window_size:
                above = length - window_size
                if win_place == 'rand':
                    diff = np.random.randint(0, above)
                else:
                    diff = round(above/2)
                new_start = row['seq_start'] + diff
                new_end = row['seq_start'] + diff + window_size
            elif length < window_size:
                missing = window_size - length
                if win_place == 'rand':
                    diff = np.random.randint(0, missing)
                else:
                    diff = round(missing/2)
                new_start = row['seq_start'] - diff
                new_end = row['seq_end'] + (missing - diff)
            else:
                new_start = row['seq_start']
                new_end = row['seq_end']

            row['seq_start'] = int(new_start)
            row['seq_end'] = int(new_end)
            return row

        def fasta_window(row):
            sequence = row['seq']
            length = len(sequence)
            if length > window_size:
                above = length - window_size
                rand = np.random.randint(0, above)
                new_sequence = sequence[rand:(rand+window_size)]
            elif length < window_size:
                missing = window_size - length
                rand = np.random.randint(0, missing)
                new_sequence = ('N' * rand) + sequence + ('N' * (missing-rand))
            else:
                new_sequence = sequence

            row['seq'] = new_sequence
            return row

        if type == 'bed':
            df = df.apply(bed_window, axis=1)
        elif type == 'fasta':
            df = df.apply(fasta_window, axis=1)

        return df
```
