## Additional file 1 for "ENNGene: an Easy Neural Network model building tool for Genomics": eval_plots.py

```
import matplotlib.pyplot as plt
import numpy as np
import os
import pandas as pd
import seaborn
import tensorflow as tf

from sklearn.metrics import average_precision_score, auc, confusion_matrix, precision_recall_curve, roc_curve


# three functions that return evaluation plots for 2 and more classes
# evaluation confusion matrix, precision recall curve and ROC curve


def plot_eval_cfm(y_true, y_pred, labels_dict, output_dir_path):
    file_path = os.path.join(output_dir_path, 'confusion_matrix')
    klass_names = list(labels_dict.keys())
    model_cfm = confusion_matrix(y_true, y_pred, labels=list(labels_dict.values()))

    figsize = (36, 30)

    cfm_sum = np.sum(model_cfm, axis=1, keepdims=True)
    cfm_perc = model_cfm / cfm_sum.astype(float) * 100
    annot = np.empty_like(model_cfm).astype(str)
    nrows, ncols = model_cfm.shape
    for i in range(nrows):
        for j in range(ncols):
            c = model_cfm[i, j]
            p = cfm_perc[i, j]
            if i == j:
                s = cfm_sum[i]
                annot[i, j] = '%.1f%%\n%d/%d' % (p, c, s)
            elif c == 0:
                annot[i, j] = ''
            else:
                annot[i, j] = '%.1f%%\n%d' % (p, c)
                
    cfm = pd.DataFrame(model_cfm)
    cfm.index.name = 'True'
    cfm.columns.name = 'Predicted'
    fig, ax = plt.subplots(figsize=figsize)
    seaborn.heatmap(cfm, cmap='Reds', annot=annot, fmt='', ax=ax, annot_kws={"fontsize": 14})
    ax.set_xlabel('Highest scoring predicted class', fontsize=20)
    ax.xaxis.set_label_position('top') 
    ax.set_xticklabels(klass_names, fontsize=18)
    ax.xaxis.tick_top()
    ax.set_ylabel('True class', fontsize=20)
    ax.set_yticklabels(klass_names, fontsize=18)
    
    plt.savefig(file_path, format='png', dpi=300)
    plt.clf()
    
                                            
def plot_multiclass_prec_recall_curve(y_test, y_pred, labels_dict, output_dir_path):
    # y_pred = np.around(y_pred)
    file_path = os.path.join(output_dir_path, 'precision_recall')
    precision = dict()
    recall = dict()
    average_precision = dict()
    klass_labels = list(labels_dict.keys())
    n_classes = len(klass_labels)

    for i in range(n_classes):
        precision[i], recall[i], _ = precision_recall_curve(y_test[:, i], y_pred[:, i])
        average_precision[i] = average_precision_score(y_test[:, i], y_pred[:, i])
        to_export = {'precision': precision[i], 'recall': recall[i]}
        pr_rec_df = pd.DataFrame.from_dict(to_export)
        pr_rec_df.to_csv(os.path.join(output_dir_path, f'precision_recall_{klass_labels[i]}.tsv'), sep='\t', index=False)

    # A "micro-average": quantifying score on all classes jointly
    precision["micro"], recall["micro"], _ = precision_recall_curve(y_test.ravel(), y_pred.ravel())
    average_precision["micro"] = average_precision_score(y_test, y_pred, average="micro")

    # A macro-average # TODO
    # precision["macro"], recall["macro"], _ = precision_recall_curve(y_test.ravel(), y_score.ravel())
    # average_precision["macro"] = average_precision_score(y_test, y_score, average="macro")

    plt.figure(figsize=(12, 12))

    f_scores = np.linspace(0.2, 0.8, num=4)
    lines = []
    labels = []
    for f_score in f_scores:
        f_score = round(f_score, 1)  # for some reason 0.6 was showing as 0.6000000000001
        x = np.linspace(0.01, 1)
        y = f_score * x / (2 * x - f_score)
        l,  = plt.plot(x[y >= 0], y[y >= 0], color='gray', alpha=0.2)
        plt.annotate(f'f1 = {f_score}', xy=(0.9, y[45] + 0.02))

    lines.append(l)
    labels.append('iso-f1 curves')

    l, = plt.plot(recall["micro"], precision["micro"], color='gold', lw=3)
    lines.append(l)
    labels.append(f'Micro-average precision-recall (AP = {round(average_precision["micro"], 4)})')

    # l, = plt.plot(recall["macro"], precision["macro"], color='gold', lw=3)
    # lines.append(l)
    # labels.append(f'Macro-average precision-recall (auc = {average_precision["macro"]})')

    avg_precisions = {}
    for i in range(n_classes):
        l, = plt.plot(recall[i], precision[i], lw=2)
        lines.append(l)
        labels.append(f'Precision-recall for {klass_labels[i]} (AP = {round(average_precision[i], 4)})')
        avg_precisions.update({klass_labels[i]: round(average_precision[i], 4)})

    plt.xlim([-0.05, 1.0])
    plt.ylim([0.0, 1.05])
    plt.xlabel('Recall', fontsize=14)
    plt.ylabel('Precision', fontsize=14)
    plt.title('Precision-Recall curve', fontsize=16)
    plt.legend(lines, labels, loc='lower left', prop=dict(size=14))
    plt.savefig(file_path, format='png', dpi=300)
    plt.clf()

    return avg_precisions


def plot_multiclass_roc_curve(test_y, y_pred, labels_dict, output_dir_path):
    # Compute ROC curve and ROC area for each class
    # y_pred = np.around(y_pred)
    file_path = os.path.join(output_dir_path, 'roc')
    fpr = dict()
    tpr = dict()
    roc_auc = dict()
    roc_auc2 = dict()
    # keras_auc = tf.keras.metrics.AUC(num_thresholds=200, curve='ROC')
    klass_labels = list(labels_dict.keys())
    n_classes = len(klass_labels)

    for i in range(n_classes):
        fpr[i], tpr[i], _ = roc_curve(test_y[:, i], y_pred[:, i])
        roc_auc[i] = auc(fpr[i], tpr[i])
        # keras_auc.update_state(test_y[:, i], y_pred[:, i])
        # roc_auc2[i] = keras_auc.result().numpy()
        to_export = {'fpr': fpr[i], 'tpr': tpr[i]}
        pr_rec_df = pd.DataFrame.from_dict(to_export)
        pr_rec_df.to_csv(os.path.join(output_dir_path, f'fpr_tpr_{klass_labels[i]}.tsv'), sep='\t', index=False)

    # Compute micro-average ROC curve and ROC area
    fpr["micro"], tpr["micro"], _ = roc_curve(test_y.ravel(), y_pred.ravel())
    roc_auc["micro"] = auc(fpr["micro"], tpr["micro"])

    # Compute macro-average ROC and ROC curve
    # First aggregate all false positive rates
    all_fpr = np.unique(np.concatenate([fpr[i] for i in range(n_classes)]))

    # Then interpolate all ROC curves at this points
    mean_tpr = np.zeros_like(all_fpr)
    for i in range(n_classes):
        mean_tpr += np.interp(all_fpr, fpr[i], tpr[i])

    # Finally average it and compute AUC
    mean_tpr /= n_classes

    fpr["macro"] = all_fpr
    tpr["macro"] = mean_tpr
    roc_auc["macro"] = auc(fpr["macro"], tpr["macro"])

    # Plot all ROC curves
    plt.figure(figsize=(12, 12))
    plt.plot(fpr["micro"], tpr["micro"], label=f'Micro-average ROC curve (auc = {round(roc_auc["micro"], 4)})', linestyle=':')
    plt.plot(fpr["macro"], tpr["macro"], label=f'Macro-average ROC curve (auc = {round(roc_auc["macro"], 4)})', linestyle=':')

    aucs = {}
    for i in range(n_classes):
        plt.plot(fpr[i], tpr[i], lw=3, label=f'ROC curve of {klass_labels[i]} (auc = {round(roc_auc[i], 4)})')
                                             # f', keras auc = {round(roc_auc2[i], 4)})')
        aucs.update({klass_labels[i]: round(roc_auc[i], 4)})
        # aucs.update({f'{klass_labels[i]} KERAS': round(roc_auc2[i], 4)})

    plt.plot([0, 1], [0, 1], 'k--', lw=3, alpha=0.2)
    plt.xlim([-0.05, 1.0])
    plt.ylim([0.0, 1.05])
    plt.xlabel('False Positive Rate (1-specifity)', fontsize=14)
    plt.ylabel('True Positive Rate (sensitivity)', fontsize=14)
    plt.title('Multiclass Receiver operating characteristic', fontsize=16)
    plt.legend(loc="lower right", prop=dict(size=14))
    plt.savefig(file_path, format='png', dpi=300)
    plt.clf()

    return aucs
```
