## Additional file 1 for "ENNGene: an Easy Neural Network model building tool for Genomics": exceptions.py

```
class MyException(Exception):

    def __init__(self, *args):
        if args:
            self.message = args[0]
        else:
            self.message = None

    def __str__(self):
        if self.message:
            return self.message
        else:
            return 'There was an error in the app code.'


class UserInputError(MyException):

    def __str__(self):
        if self.message:
            return self.message
        else:
            return 'User gave wrong input.'


class ProcessError(MyException):

    def __str__(self):
        if self.message:
            return self.message
        else:
            return 'Failed to finish a part of app process.'
```
