## Additional file 1 for "ENNGene: an Easy Neural Network model building tool for Genomics": file_utils.py

```
# module containing methods for file handling
import gzip
import os
from zipfile import ZipFile


def list_files_in_dir(path, extension='*'):
    file_paths = []
    for root, _, files in os.walk(path):
        for file in files:
            if extension in file:
                file_paths.append(os.path.join(root, file))
    return file_paths


def write(path, content):
    file = open(path, 'w')
    file.write(content)
    file.close()


def unzip_if_zipped(zipped_file):
    if ".gz" in zipped_file:
        file = gzip.open(zipped_file, 'r')
        zipped = True
    elif ".zip" in zipped_file:
        file = ZipFile(zipped_file).extractall()
        zipped = True
    else:
        file = open(zipped_file)
        zipped = False
    return file, zipped


def read_decoded_line(opened_file, zipped):
    line = opened_file.readline().strip()
    if zipped:
        line = line.decode('utf-8')

    return line
```
