## Additional file 1 for "ENNGene: an Easy Neural Network model building tool for Genomics": ig.py

```
#@title Licensed under the Apache License, Version 2.0
#https://www.apache.org/licenses/LICENSE-2.0

from os import path
from numpy.lib.function_base import diff
import tensorflow as tf
import numpy as np
import matplotlib as mpl
import matplotlib.cm as cm

import logging
logger = logging.getLogger('ig')


def _expand_dims(x):
    return tf.expand_dims(x, axis=0)

def _get_path_calculator(alphas):
    def _calculate_path(baseline, inp):
        return baseline + alphas * (inp - baseline)
    return _calculate_path

def generate_path_inputs(baselines, inputs, alphas):
    """
    Generate interpolated 'images' along a linear path at alpha intervals between a baseline tensor

    baselines: 2D, shape: (win, width)
    input_seq: preprocessed sample, shape: (win, width)
    input_fold: preprocessed sample, shape: (win, width)

    return: shape [(alphas_len, win, width), (alphas_len, win, width2),]
    """
    
    baselines = map(_expand_dims, baselines)
    inputs = map(_expand_dims, inputs)
    
    path_calculator = _get_path_calculator(alphas)
    
    return [path_calculator(base, inp) for base, inp in zip(baselines, inputs)]


def compute_gradients(model, path_inputs, target_class=0):
    """
    compute dependency of each field on whole result, compared to interpolated 'images'

    :param model: trained model
    :param path_inputs: interpolated tensors, shape: (alphas_len, win, width)
    :return: shape: (alphas_len, win, width)
    """
    with tf.GradientTape() as tape:
        tape.watch(path_inputs)
        predictions = model(path_inputs)
        outputs = tf.convert_to_tensor(
            [envelope[target_class] for envelope in predictions], 
            dtype=tf.float32
        )

    gradients = tape.gradient(outputs, path_inputs)
    return gradients


def generate_alphas(m_steps=50, method='riemann_trapezoidal'):
    """
    Args:
    m_steps(Tensor): A 0D tensor of an int corresponding to the number of linear
    interpolation steps for computing an approximate integral. Default is 50.
    method(str): A string representing the integral approximation method. The
       following methods are implemented:
      - riemann_trapezoidal(default)
      - riemann_left
      - riemann_midpoint
      - riemann_right
    Returns:
      alphas(Tensor): A 1D tensor of uniformly spaced floats with the shape
      (m_steps,).
      """
    m_steps_float = tf.cast(m_steps, float)

    if method == 'riemann_trapezoidal':
        alphas = tf.linspace(0.0, 1.0, m_steps+1)
    elif method == 'riemann_left':
        alphas = tf.linspace(0.0, 1.0 - (1.0 / m_steps_float), m_steps)
    elif method == 'riemann_midpoint':
        alphas = tf.linspace(1.0 / (2.0 * m_steps_float), 1.0 - 1.0 / (2.0 * m_steps_float), m_steps)
    elif method == 'riemann_right':
        alphas = tf.linspace(1.0 / m_steps_float, 1.0, m_steps)
    else:
        raise AssertionError("Provided Riemann approximation method is not valid.")

    return alphas


def integral_approximation(gradients, method='riemann_trapezoidal'):
    """Compute numerical approximation of integral from gradients.
    Args:
    gradients(Tensor): A 4D tensor of floats with the shape
    (m_steps, img_height, img_width, 3).
    method(str): A string representing the integral approximation method. The
    following methods are implemented:
    - riemann_trapezoidal(default)
    - riemann_left
    - riemann_midpoint
    - riemann_right
    Returns:
    integrated_gradients(Tensor): A 3D tensor of floats with the shape
    (img_height, img_width, 3).
    """
    if method == 'riemann_trapezoidal':
        grads = (gradients[:-1] + gradients[1:]) / tf.constant(2.0)
    elif method == 'riemann_left':
        grads = gradients
    elif method == 'riemann_midpoint':
        grads = gradients
    elif method == 'riemann_right':
        grads = gradients
    else:
        raise AssertionError("Provided Riemann approximation method is not valid.")

    # Average integration approximation.
    integrated_gradients = tf.math.reduce_mean(grads, axis=0)

    return integrated_gradients


def integrated_gradients(model, baselines, inputs, target_class,
                         m_steps=50, method='riemann_trapezoidal', batch_size=50):
    """
    Args:
      model(keras.Model): A trained model to generate predictions and inspect.
      baselines(Tensor): List of 2D, shape: (win, width)
      inputs(Tensor): List of preprocessed samples, shape: (win, width)
      m_steps(Tensor): A 0D tensor of an integer corresponding to the number of
        linear interpolation steps for computing an approximate integral.
      method(str): A string representing the integral approximation method. The
        following methods are implemented:
        - riemann_trapezoidal(default)
        - riemann_left
        - riemann_midpoint
        - riemann_right
      batch_size(Tensor): A 0D tensor of an integer corresponding to a batch
        size for alpha to scale computation and prevent OOM errors. Note: needs to
        be tf.int64 and shoud be < m_steps. Default value is 32.
    Returns:
      integrated_gradients(Tensor): A 2D tensor of floats with the same
        shape as the input tensor.
    """
    alphas = generate_alphas(m_steps=m_steps, method=method)

    # Initialize TensorArray outside loop to collect gradients. Note: this data structure
    gradient_batches = [tf.TensorArray(tf.float32, size=m_steps + 1) for _ in inputs]

    # Iterate alphas range and batch computation for speed, memory efficiency, and scaling to larger m_steps.
    for alpha in tf.range(0, len(alphas), batch_size):
        from_ = alpha
        to = tf.minimum(from_ + batch_size, len(alphas))
        alpha_batch = alphas[from_:to]

        # 2. Generate interpolated inputs between baseline and input.
        interpolated_path_input_batch_list = generate_path_inputs(baselines, inputs, alpha_batch[:, tf.newaxis, tf.newaxis])
        

        # 3. Compute gradients between model outputs and interpolated inputs.
        new_batch_list = compute_gradients(
            model=model,
            path_inputs=interpolated_path_input_batch_list,
            target_class=target_class
        )
        

        # Write batch indices and gradients to TensorArray.
        gradient_batches = [branch.scatter(tf.range(from_, to), new_batch_to_branch) 
                            for branch, new_batch_to_branch in zip(gradient_batches, new_batch_list)]

    # Stack path gradients together row-wise into single tensor.
    gradient_batches = [branch.stack() for branch in gradient_batches]

    # 4. Integral approximation through averaging gradients.
    avg_gradients = [integral_approximation(gradients=gradients, method=method) for gradients in gradient_batches]


    # 5. Scale integrated gradients with respect to input.
    return [(input_ - baseline) * avg_gradient for input_, baseline, avg_gradient in zip(inputs, baselines, avg_gradients)]


def smoothgrad(model, baselines, inputs, target_class,
                         m_steps=50, method='riemann_trapezoidal', batch_size=50, 
                         stddev=0.15, smoothing_repetitions=20):
    
    results = []
    input_maxes = [np.max(_input) for _input in inputs]
    input_mins = [np.min(_input) for _input in inputs]
    gauss_bases = [max_ - min_ for max_, min_ in zip(input_maxes, input_mins)]    
    
    for _ in range(smoothing_repetitions):
        stddev_list = [np.random.normal(scale=stddev*(gauss_base), size=input_.shape) for input_, gauss_base in zip(inputs, gauss_bases)]
        inputs_plus_stddev = [sum(x) for x in zip(inputs, stddev_list)]
        results.append(integrated_gradients(model, baselines, inputs_plus_stddev, target_class, m_steps, method, batch_size))
        
    
    return [sum(i) / smoothing_repetitions for i in zip(*results)]

def _absmax(a, axis=1):
    amax = np.max(a, axis)
    amin = np.min(a, axis)
    return np.where(-amin > amax, amin, amax)

def choose_validation_points(integrated_gradients_list):
    """
    Args:
          integrated_gradients_list(Tensor): A list of 2D tensor of floats with shape (window_size, width_of_sequence_encoded).
          window_size: int, length of sequence, num of bases
          width: int, width of encoded base
    Return: List of attributes for highlighting DNA string sequence
    """
    return [_absmax(x) for x in integrated_gradients_list]


def visualize_token_attrs(sequence, attrs, _min, _max, cmap=cm.coolwarm):
    """
    Visualize attributions for given set of tokens.
    Args:
    - tokens: An array of tokens
    - attrs: An array of attributions, of same size as 'tokens',
      with attrs[i] being the attribution to tokens[i]

    Returns:
    - visualization: HTML text with colorful representation of DNA sequence
        build on model prediction
    """
    norm = mpl.colors.Normalize(vmin=_min, vmax=_max)
    m = cm.ScalarMappable(norm=norm, cmap=cmap)
    
    html_text = []
    
    for i, tok in enumerate(sequence):
        r, g, b, _ = m.to_rgba(attrs[i]) # ignore alpha
        r, g, b = 255*r, 255*g, 255*b # rescale
        html_text.append("<span style='font-weight:bold;background-color:rgb(%d,%d,%d)'>%s </span>" % (r, g, b, tok))


    html_text.append("</div><br>")
    
    return html_text
```
