## Additional file 1 for "ENNGene: an Easy Neural Network model building tool for Genomics": sequence.py

```
import numpy as np
import streamlit as st
import _io

from . import file_utils as f
from .exceptions import UserInputError

# TODO allow option custom, to be specified by text input
# TODO add amino acid alphabet - in that case disable cons and fold i guess
ALPHABET = {'A': 0, 'C': 1, 'G': 2, 'T': 3, 'U': 3}
FOLDING = {'.': 0, '|': 1, 'x': 2, '<': 3, '>': 4, '(': 5, ')': 6}


@st.cache(hash_funcs={_io.TextIOWrapper: lambda _: None}, suppress_st_warning=True)
def read_and_cache(fasta):
    with st.spinner('Parsing reference fasta file to infer the available chromosomes. May take up to few minutes...'):
        parsed = parse_fasta_reference(fasta)
    return parsed


def parse_fasta_reference(fasta_file):
    chromosomes = []

    key = None
    value = ""

    file, zipped = f.unzip_if_zipped(fasta_file)
    while True:
        line = f.read_decoded_line(file, zipped)
        if not line:
            break
    
        if '>' in line:
            # Save finished previous key value pair (unless it's the first iteration)
            if key:  # and is_valid_chr(key):
                # Save only sequence for chromosomes we are interested in (skip scaffolds etc.)
                chromosomes.append(key)

            key = line.strip().strip('>')
            value = ""
        else:
            if key:
                line = line.strip()
                value += line
                l = [char for char in line.upper()]
            else:
                raise UserInputError("Provided reference file does not start with '>' fasta identifier.")

    chromosomes.append(key)  # save the last one
    file.close()
    chromosomes.sort()

    return chromosomes


# def is_valid_chr(chromosome):
#     return not not re.search(r'^(chr)*((\d{1,3})|(M|m|MT|mt|x|X|y|Y))$', chromosome)


def parse_wig_header(line):
    # example: fixedStep chrom=chr22 start=10510001 step=1 # may also contain span (default = 1)
    header = {'span': 1}

    parts = line.split()
    file_type = parts.pop(0)
    header.update({'file_type': file_type})

    if file_type not in ['fixedStep', 'variableStep']:
        raise UserInputError(f'Unknown type of wig file provided: {file_type}. Only fixedStep or variableStep allowed.')

    for part in parts:
        key, value = part.split('=')
        if key == 'chrom':
            header.update({key: value})
        elif key == 'start':
            header.update({key: int(value) - 1})
        elif key in ['span', 'step']:
            header.update({key: int(value)})

    return header


def parse_wig_line(line, header):
    parsed_line = {}
    if header['file_type'] == 'variableStep':
        parts = line.split()
        start = int(parts[0]) - 1
        value = float(parts[1])
        for i in range(header['span']):
            coord = start + i
            parsed_line.update({coord: value})
        header['start'] = start + header['span']
    elif header['file_type'] == 'fixedStep':
        value = float(line)
        for i in range(header['span']):
            coord = header['start'] + i
            parsed_line.update({coord: value})
        header['start'] += header['step']

    return [header, parsed_line]


def complement(sequence_list, dictionary):
    return [dictionary[base] for base in sequence_list]


def onehot_encode_alphabet(alphabet):
    encoded_alphabet = {}
    for char, pos in alphabet.items():
        ohe = np.zeros(len(set(alphabet.values())))
        ohe[pos] = 1
        encoded_alphabet.update({char: ohe})
    if alphabet == ALPHABET:
        encoded_alphabet.update({'N': np.full(len(set(alphabet.values())), 0.25)})

    return encoded_alphabet


def translate(char, encoding):
    if not char:
        return None
    if not encoding or not (encoding.__class__.__name__ == 'dict'):
        raise ValueError('Encoding missing...')

    if char in encoding.keys():
        return encoding[char]
    elif char.upper() in encoding.keys():
        return encoding[char.upper()]
    else:
        raise UserInputError(f"Invalid character '{char}' found, given encoding {encoding}. "
                         "Provided encoding must contain all possible characters (case-insensitive).")
```
