## Additional file 1 for "ENNGene: an Easy Neural Network model building tool for Genomics": subcommand.py

```
from datetime import datetime
import logging
import numpy as np
import os
from pathlib import Path
import matplotlib.pyplot as plt
import seaborn as sns
import pandas as pd
import shutil
import streamlit as st
import streamlit.components.v1 as stcomponents
import tensorflow as tf
from tqdm import tqdm
import yaml

from . import eval_plots
from . import ig
from . import validators
from .exceptions import UserInputError

logger = logging.getLogger('root')


# noinspection PyAttributeOutsideInit
class Subcommand:

    BRANCHES = {'Sequence': 'seq',
                'Conservation score': 'cons',
                'Secondary structure': 'fold'}
    OPTIMIZERS = {'SGD': 'sgd',
                  'RMSprop': 'rmsprop',
                  'Adam': 'adam'}
    LR_OPTIMS = {'Fixed lr': 'fixed',
                 'LR scheduler': 'lr_scheduler',
                 # 'LR finder': 'lr_finder',
                 'One cycle policy': 'one_cycle'}
    WIN_PLACEMENT = {'Centered': 'center',
                     'Randomized': 'rand'}

    def general_options(self):
        self.params_loaded = False
        self.defaults = {}
        self.defaults.update(self.default_params())
        self.load_params = st.checkbox('Load parameters from a previous run', value=False)

        if self.load_params:
            folder = st.text_input('Folder from the previous run of the task (must contain the parameters.yaml file)')
            if folder:
                if os.path.isdir(folder):
                    param_files = [file for file in os.listdir(folder) if (file == 'parameters.yaml') and
                                   (os.path.isfile(os.path.join(folder, file)))]
                    if len(param_files) >= 1 and os.path.isfile(os.path.join(folder, param_files[0])):
                        param_file = os.path.join(folder, param_files[0])
                        with open(param_file, 'r') as file:
                            try:
                                user_params = yaml.safe_load(file)
                            except Exception as err:
                                logger.exception(f'{err.__class__.__name__}: {err}')
                                raise UserInputError('An error occurred while processing given yaml file.')
                            if self.__class__.__name__ in user_params.keys():
                                # For v <= 1.0.1 the numbers were saved separately in a dict TODO remove eventually
                                if self.__class__.__name__ == 'Train' and \
                                        'no_branches_layers' in user_params['Train'].keys() and \
                                        type(user_params['Train']['no_branches_layers']) == dict:
                                    user_params['Train']['no_branches_layers'] = max(list(user_params['Train']['no_branches_layers'].values()))
                                self.defaults.update(user_params[self.__class__.__name__])
                                self.params_loaded = True
                            else:
                                raise UserInputError('Given yaml file does not contain parameters for the currently selected task.')
                    else:
                        raise UserInputError('No yaml file was found in the given folder.')
                else:
                    raise UserInputError('Given folder does not exist.')
        self.params.update(self.defaults)

        self.params['output_folder'] = st.text_input(
            'Output folder (result files will be exported here; home directory used as default)',
            value=self.defaults['output_folder']
        )
        st.markdown('---')

    # Used by Evaluation and Prediction modules
    def model_options(self, blackbox=False, warning=None):
        missing_model = False
        missing_params = False
        model_types = {'Use a model trained by the ENNGene': 'from_app',
                       'Use a custom trained model': 'custom'}

        st.markdown(warning) if warning else None
        self.params['model_source'] = model_types[st.radio(
            'Select a source of the trained model:',
            list(model_types.keys()), index=self.get_dict_index(self.defaults['model_source'], model_types))]
        if self.params['model_source'] == 'from_app':
            self.params['model_folder'] = st.text_input('Training folder containing the model (hdf5 file)',
                                              value=self.defaults['model_folder'])
            if self.params['model_folder'] and os.path.isdir(self.params['model_folder']):
                model_files = [f for f in os.listdir(self.params['model_folder']) if f.endswith('.hdf5') and
                               os.path.isfile(os.path.join(self.params['model_folder'], f))]
                if len(model_files) == 0:
                    missing_model = True
                    st.markdown('#### Sorry, there is no hdf5 file in given folder.')
                elif len(model_files) == 1:
                    self.params['model_file'] = os.path.join(self.params['model_folder'], model_files[0])
                    st.markdown(f"###### Model file: {self.params['model_file']}")
                elif len(model_files) > 1:
                    missing_model = True
                    st.markdown(
                        '#### Sorry, there is too many hdf5 files in the given folder. Please specify the model file below.')

                if not blackbox:
                    previous_param_files = [f for f in os.listdir(self.params['model_folder']) if (f == 'parameters.yaml') and
                                   (os.path.isfile(os.path.join(self.params['model_folder'], f)))]
                    if len(previous_param_files) == 0:
                        missing_params = True
                        st.markdown('#### Sorry, could not find parameters.yaml file in the given folder. '
                                    'Check the folder or specify the parameters below.')
                    elif len(previous_param_files) == 1:
                        training_params = {'win': None, 'no_klasses': 0, 'klasses': [], 'branches': []}
                        self.previous_param_file = os.path.join(self.params['model_folder'], previous_param_files[0])
                        with open(self.previous_param_file, 'r') as file:
                            user_params = yaml.safe_load(file)
                        try:
                            training_params['win'] = user_params['Preprocess']['win']
                            klasses = user_params['Preprocess']['klasses']
                            training_params['no_klasses'] = len(klasses)
                            training_params['klasses'] = klasses
                            training_params['branches'] = user_params['Train']['branches']
                        except:
                            missing_params = True
                            st.markdown('#### Sorry, could not read the parameters from given folder. '
                                        'Check the folder or specify the parameters below.')
                        if not training_params['win'] \
                                or training_params['no_klasses'] == 0 or len(training_params['klasses']) == 0 \
                                or len(training_params['klasses']) != training_params['no_klasses'] \
                                or len(training_params['branches']) == 0:
                            missing_params = True
                            st.markdown('#### Sorry, could not read the parameters from given folder. '
                                        'Check the folder or specify the parameters below.')
                        else:
                            st.markdown('###### Parameters read from given folder:\n'
                                        f"* Window size: {training_params['win']}\n"
                                        f"* No. of classes: {training_params['no_klasses']}\n"
                                        f"* Class labels: {', '.join(training_params['klasses'])}\n"
                                        f"* Branches: {', '.join([self.get_dict_key(b, self.BRANCHES) for b in training_params['branches']])}")
                            self.params.update(training_params)
                    if len(previous_param_files) > 1:
                        missing_params = True
                        st.markdown('#### Sorry, there is too many parameters.yaml files in the given folder. '
                                    'Check the folder or specify the parameters below.')

        if self.params['model_source'] == 'custom' or missing_params or missing_model:
            if not missing_params:
                self.params['model_file'] = st.text_input('Path to the trained model (hdf5 file)',
                                                          value=self.defaults['model_file'])
            if (self.params['model_source'] == 'custom' or missing_params) and not blackbox:
                st.markdown('##### **WARNING:** Parameters window size, branches, and number of classes must be the same as when used for training the given model.')
                self.params['win'] = int(
                    st.number_input('Window size used for training', min_value=3, value=self.defaults['win']))
                self.params['no_klasses'] = int(st.number_input('Number of classes used for training', min_value=2,
                                                                value=self.defaults['no_klasses']))
                st.markdown('##### **WARNING:** Make sure the class order is the same as when training the model.')
                for i in range(self.params['no_klasses']):
                    if len(self.params['klasses']) >= i + 1:
                        value = self.params['klasses'][i]
                    else:
                        value = str(i)
                        self.params['klasses'].append(value)
                    self.params['klasses'][i] = st.text_input(f'Class {i} label:', value=value)
                self.params['branches'] = list(map(lambda name: self.BRANCHES[name],
                                                   st.multiselect('Branches',
                                                                  list(self.BRANCHES.keys()))))

        self.validation_hash['is_model_file'].append(self.params['model_file'])

    def sequence_options(self, seq_types, evaluation):
        if 'cons' in self.params['branches']:
            if evaluation:
                seq_types = {'BED file': 'bed', 'Blackbox dataset': 'blackbox'}
                self.params['seq_type'] = seq_types[st.radio(
                    'Select a source of the sequences:',
                    list(seq_types.keys()), index=self.get_dict_index(self.defaults['seq_type'], seq_types))]
            else:
                # to map to the conservation files we need the coordinates
                self.params['seq_type'] = 'bed'
                st.markdown(
                    '###### Note: Only BED files allowed when Conservation score branch is applied (the coordinates are necessary).')
        else:
            self.params['seq_type'] = seq_types[st.radio(
                'Select a source of the sequences:',
                list(seq_types.keys()), index=self.get_dict_index(self.defaults['seq_type'], seq_types))]
        self.params['win_place'] = self.WIN_PLACEMENT[st.radio(
            'Choose a way to place the window upon the sequence:',
            list(self.WIN_PLACEMENT.keys()), index=self.get_dict_index(self.defaults['win_place'], self.WIN_PLACEMENT))]
        self.references = {}
        if self.params['seq_type'] == 'bed':
            self.params['seq_source'] = st.text_input(
                'Path to the BED file containing intervals to be classified', value=self.defaults['seq_source'])
            if evaluation:
                st.markdown(
                    '###### Note: The first (extra) column of the file must contain the name of the class per each sequence. '
                    'Class names must correspond to those used when training the model.')
            self.validation_hash['is_bed'].append({'file': self.params['seq_source'], 'evaluation': evaluation})
            self.params['strand'] = st.checkbox('Apply strand', self.defaults['strand'])

            if 'seq' in self.params['branches'] or 'fold' in self.params['branches']:
                self.params['fasta_ref'] = st.text_input('Path to the reference fasta file',
                                                         value=self.defaults['fasta_ref'])
                self.references.update({'seq': self.params['fasta_ref'], 'fold': self.params['fasta_ref']})
                self.validation_hash['is_fasta'].append(self.params['fasta_ref'])
            if 'cons' in self.params['branches']:
                self.params['cons_dir'] = st.text_input('Path to folder containing reference conservation files',
                                                        value=self.defaults['cons_dir'])
                self.references.update({'cons': self.params['cons_dir']})
                self.validation_hash['is_wig_dir'].append(self.params['cons_dir'])

        elif self.params['seq_type'] == 'fasta' or self.params['seq_type'] == 'text':
            st.markdown('###### WARNING: Sequences shorter than the window size will be padded with Ns (may affect '
                        'the prediction accuracy). Longer sequences will be cut to the length of the window.')
            if self.params['seq_type'] == 'fasta':
                self.params['seq_source'] = st.text_input(
                    'Path to FASTA file containing sequences to be classified', value=self.defaults['seq_source'])
                if evaluation:
                    st.markdown(
                        "###### Note: The class name must be provided as a last part of the header, separated by a space. E.g. '>chr16:655478-655578 FUS_positives'. "
                        'Class names must correspond to those used when training the model.')
                self.validation_hash['is_fasta'].append(self.params['seq_source'])

            elif self.params['seq_type'] == 'text':
                self.params['seq_source'] = st.text_area(
                    'One or more sequences to be classified (each sequence on a new line)',
                    value=self.defaults['seq_source'])
                self.validation_hash['is_multiline_text'].append(self.params['seq_source'])
        elif self.params['seq_type'] == 'blackbox':
            self.params['seq_source'] = st.text_input(
                'Path to the Blackbox dataset file exported from the Preprocess module', value=self.defaults['seq_source'])
            st.markdown(
                '###### Note: Dataset should come from the same data as those used for training the model, '
                'or the parameters must match at least (e.g. class names, window size, branches...).')

            self.validation_hash['is_blackbox'].append(self.params['seq_source'])

        if 'fold' in self.params['branches']:
            # currently used only as an option for RNAfold
            max_cpu = os.cpu_count() or 1
            self.ncpu = st.slider('Number of CPUs to be used for folding (max = all available CPUs on the machine).',
                                  min_value=1, max_value=max_cpu, value=max_cpu)
        else:
            self.ncpu = 1

    def evaluate_model(self, encoded_labels, model, test_x, test_y, params, out_dir):
        test_results = model.evaluate(
            test_x,
            test_y,
            verbose=1,
            sample_weight=None)
        y_pred = model.predict(test_x, verbose=1)

        self.log_eval_metrics(test_results, params)

        # Plot evaluation metrics
        # categorical_labels = {key: i for i, (key, _) in enumerate(encoded_labels.items())}
        aucs = eval_plots.plot_multiclass_roc_curve(test_y, y_pred, encoded_labels, out_dir)
        avg_precisions = eval_plots.plot_multiclass_prec_recall_curve(test_y, y_pred, encoded_labels, out_dir)
        # FIXME
        # eval_plots.plot_eval_cfm(np.argmax(test_y, axis=1), np.argmax(y_pred, axis=1), categorical_labels, out_dir)
        self.log_plotted_metrics(aucs, avg_precisions, params)

        return y_pred

    @staticmethod
    def get_klass(predicted, klasses):
        #TODO give the user choice of the tresshold value? - would have to specify per each class, if the highest scoring class would be above its threshold, then we would call it, otherwise uncertain

        # treshold = 0.98
        # chosen = []
        # for probs in predicted:
        #     max_prob = np.amax(probs)
        #     if max_prob > treshold:
        #         not considering an option there would be two exactly same highest probabilities
        # index = np.where(probs == max_prob)[0][0]
        # value = klasses[index]
        # else:
        #     value = 'UNCERTAIN'
        # chosen.append(value)

        chosen = [klasses[np.argmax(probs)] for probs in predicted]

        return chosen

    @staticmethod
    def log_eval_metrics(test_results, params):
        params['eval_loss'] = str(round(test_results[0], 4))
        params['eval_acc'] = str(round(test_results[1], 4))

        logger.info('Evaluation loss: ' + params['eval_loss'])
        logger.info('Evaluation acc: ' + params['eval_acc'])

        st.text(f"Evaluation loss: {params['eval_loss']} \n"
                f"Evaluation accuracy: {params['eval_acc']} \n")

    @staticmethod
    def log_plotted_metrics(aucs, avg_precisions, params):
        auc_cell = ''
        for klass, auc in aucs.items():
            auc_cell += f'{klass}: {auc}, '
        params['auc'] = auc_cell.strip(', ')
        ap_cell = ''
        for klass, ap in avg_precisions.items():
            ap_cell += f'{klass}: {ap}, '
        params['avg_precision'] = ap_cell.strip(', ')

        logger.info('AUC: ' + params['auc'])
        logger.info('Average precision: ' + params['avg_precision'])

        auc_rows = ''
        for klass, auc in aucs.items():
            auc_rows += f'{klass}: {auc}\n'
        st.text(f'AUC \n{auc_rows}')

        ap_rows = ''
        for klass, ap in avg_precisions.items():
            ap_rows += f'{klass}: {ap}\n'
        st.text(f'Average precision \n{ap_rows}')

    def validate_and_run(self, validation_hash):
        st.markdown('---')
        if st.button('Run'):
            warnings = self.validate_input(validation_hash)
            if len(warnings) == 0:
                logger.info('\n'.join(['%s: %s' % (key, value) for (key, value) in self.params.items()]))
                self.run()
            else:
                st.warning('  \n'.join(warnings))

    @staticmethod
    def validate_input(validation_hash):
        warnings = []
        for validator, items in validation_hash.items():
            for item in items:
                if type(item) == dict:
                    warnings.append(getattr(validators, validator)(**item))
                else:
                    warnings.append(getattr(validators, validator)(item))

        return list(filter(None, warnings))

    @staticmethod
    def spent_time(time1):
        time2 = datetime.now()
        t = time2 - time1
        return t

    @staticmethod
    def ensure_dir(dir_path):
        if not os.path.exists(dir_path):
            try:
                os.makedirs(dir_path)
            except Exception:
                raise UserInputError(f"Failed to create output folder at given path: {dir_path}.")

    @staticmethod
    def get_dict_index(value, dictionary):
        return list(dictionary.values()).index(value)

    @staticmethod
    def get_dict_key(value, dictionary):
        index = Subcommand.get_dict_index(value, dictionary)
        return list(dictionary.keys())[index]


    @staticmethod
    def cons_to_symbol(cons):
        cons = np.round((cons / 0.5) + 5)
        if cons > 9:
            return "+ "
        elif cons < 0:
            return "- "
        else:
            return str(int(cons)) + " "
    
    
    @staticmethod
    def visualize_specifier(branches):
        LETTER_HEIGHT = 20
               
        def visualize(row):
            sub_viz = []
            _max = -np.inf
            _min = np.inf
            
            for branch in branches:
                _max = max(_max, row[branch + "_ig"].max()) 
                _min = min(_max, row[branch + "_ig"].min())
                
            
            visualisation = {
                branch: ig.visualize_token_attrs(row[branch], row[branch+"_ig"], _min, _max) for branch in branches
            }
            
            sequence_len = len(row[branches[0]]) # just get sequence length, should be equal across all branches
            max_text_len = len(str(sequence_len)) + 1
        
            display_rows_count = sequence_len//25
            
            
            for sequence_pos in range(display_rows_count):
                start = str(sequence_pos*25) 
                end = str((sequence_pos+1)*25) 
                
                sub_viz.append("<div style='font-family: monospace, monospace'>")
                
                # funky sort to ensure seq -> fold -> cons order
                for i_branch, branch in enumerate(sorted(branches)[::-1]):
                    if i_branch == 0:
                        sub_viz.append(start + '<span style="opacity:0;">' + (max_text_len-len(start))*"_" +'</span>')
                        sub_viz.extend(visualisation[branch][sequence_pos*25:(sequence_pos+1)*25])
                        sub_viz.append( '<span style="opacity:0;">' + (max_text_len-len(end))*"_" +'</span>' + end)
                    else:
                        sub_viz.append('<span style="opacity:0;">' + max_text_len*"_" +'</span>')
                        sub_viz.extend(visualisation[branch][sequence_pos*25:(sequence_pos+1)*25])
                        sub_viz.append('<span style="opacity:0;">' + max_text_len*"_" +'</span>')
                        
                    sub_viz.append(" " + branch)                    
                    sub_viz.append("<br>")
                
                sub_viz.append("</div>")
                sub_viz.append("<br>")
                
            
            sub_viz.append("<hr>")
            stcomponents.html("".join(sub_viz), height=LETTER_HEIGHT*(len(branches)+1)*(display_rows_count))
            sub_viz = []
                
        
        return visualize
    
    @staticmethod
    def calculate_ig(dataset, model, predict_x, klasses, branches, use_smoothgrad=False):
        if not isinstance(predict_x, list):
            logger.info("predict_x is not a list, wrapping in an array")
            predict_x = (np.array(predict_x),)
                        
        # baseline of zeros in equal shape as inputs
        baselines = [tf.zeros(shape=x[0].shape) for x in predict_x]
        top_klasses = [klasses.index(klass) for klass in dataset.df['highest scoring class']]

        ig_per_branch = { branch: [] for branch in branches }
      

        # take each prediction, unprocessed data and count IG
        for inputs, target_class in tqdm(zip(zip(*predict_x), top_klasses), total=len(predict_x[0])): 
            # zip(*predict_x) decompress a list of n lists into a single list of tuples
            # e. g. [[a,b], [c,d]] becomes [(a,c), (b,d)] ; [[a, b, c]] becomes [(a,), (b,), (c,)]
            
            # return tensor of shape: (window width(sequence length), encoded base shape)
            inputs = [tf.convert_to_tensor(_input, dtype=tf.float32) for _input in inputs]
            
            # contain significance of each base in sequence
            if use_smoothgrad:
                ig_method = ig.smoothgrad
            else:
                ig_method = ig.integrated_gradients
            
            ig_atributions = ig_method(model, baselines, inputs, target_class)

            # choose attribution for specific encoded base
            selected_ig_atributions = ig.choose_validation_points(ig_atributions)
            
            for branch, ig_atribution in zip(branches, selected_ig_atributions):
                ig_per_branch[branch].append(ig_atribution)

        for branch in branches:
            dataset.df[branch + "_ig"] = ig_per_branch[branch]
            
        # Show ten best predictions per class in the application window
        st.markdown('---')
        st.markdown('### Integrated Gradients Visualisation')
        st.markdown('Below are ten sequences with highest predicted score per each class. \n'
                    'You can find html visualisation code for all the sequences in the results.tsv file.\n\n'
                    'The higher is the attribution of the sequence to the prediction, the more pronounced is its red color. '
                    'On the other hand, the blue color means low level of attribution.')
        best = dataset.df[klasses + [branch+'_ig' for branch in branches] + branches]

        visualize = Subcommand.visualize_specifier(branches)
        
        for klass in klasses:
            st.markdown(f'#### {klass}')
            best_ten = best.sort_values(by=klass, ascending=False, inplace=False)[:10]
            
            for _, row in best_ten.iterrows():    
                if 'cons' in branches:
                    row['cons'] = [Subcommand.cons_to_symbol(cons_score) for cons_score in  row['cons']] 
                visualize(row)

    @staticmethod
    def finalize_run(logger, out_dir, user_params, csv_header, csv_row, placeholder=None, previous_param_file=None):
        place = placeholder or st.empty()
        place.text(f'You can find your results at {out_dir}')
        params = user_params.copy()
        task = params.pop('task')
        params = {task: user_params}
        if previous_param_file:
            with open(previous_param_file, 'r') as file:
                previous_params = yaml.safe_load(file)
            params.update(previous_params)

        with open(os.path.join(out_dir, 'parameters.yaml'), 'w') as file:
            yaml.dump(params, file)

        parent_dir = Path(out_dir).parent
        table_file = os.path.join(parent_dir, 'parameters.tsv')
        if table_file:
            write_header = not os.path.isfile(table_file)
            with open(table_file, 'a') as file:
                file.write(csv_header) if write_header else None
                file.write(csv_row)
                pass

        file_handler = [handler for handler in logger.handlers if type(handler) == logging.FileHandler]
        if file_handler:
            logfile_path = file_handler[0].baseFilename
            logfile_name = os.path.basename(logfile_path)
            shutil.move(logfile_path, os.path.join(out_dir, logfile_name))

    @staticmethod
    def preprocess_header():
        return 'Preprocess directory\t' \
               'Preprocess branches\t' \
               'Strand\t' \
               'Window\t' \
               'Window placement\t' \
               'Split\t' \
               'Split ratio\t' \
               'Chromosomes\t' \
               'Reduced classes\t' \
               'Reduce ratio\t' \
               'Use mapped\t' \
               'Input files\t' \
               'Classes\t' \
               'Full_dataset_file\t' \
               'Fasta ref\t' \
               'Conservation ref\t'

    def preprocess_row(self, params):
        return f"{os.path.basename(params['datasets_dir'])}\t" \
               f"{[self.get_dict_key(b, self.BRANCHES) for b in params['branches']]}\t" \
               f"{'Yes' if (params['strand'] and 'seq' in params['branches']) else ('No' if 'seq' in params['branches'] else '-')}\t" \
               f"{params['win']}\t" \
               f"{self.get_dict_key(params['win_place'], self.WIN_PLACEMENT)}\t" \
               f"{'Random' if params['split'] == 'rand' else 'By chromosomes'}\t" \
               f"{params['split_ratio'] if params['split'] == 'rand' else '-'}\t" \
               f"{params['chromosomes'] if params['split'] == 'by_chr' else '-'}\t" \
               f"{params['reducelist'] if len(params['reducelist']) != 0 else '-'}\t" \
               f"{params['reduceratio'] if len(params['reducelist']) != 0 else '-'}\t" \
               f"{'Yes' if params['use_mapped'] else 'No'}\t" \
               f"{params['input_files']}\t" \
               f"{params['klasses']}\t" \
               f"{params['full_dataset_file'] if params['use_mapped'] else '-'}\t" \
               f"{params['fasta'] if params['fasta'] else '-'}\t" \
               f"{params['cons_dir'] if params['cons_dir'] else '-'}\t"

    @staticmethod
    def train_header():
        return 'Training directory\t' \
               'Evaluation loss\t' \
               'Evaluation accuracy\t' \
               'AUC\t' \
               'Average precision\t' \
               'Training loss\t' \
               'Training accuracy\t' \
               'Validation loss\t' \
               'Validation accuracy\t' \
               'Training branches\t' \
               'Batch size\t' \
               'Optimizer\t' \
               'Learning rate\t' \
               'LR optimizer\t' \
               'Epochs\t' \
               'No. branches layers\t' \
               'Branches layers\t' \
               'No. common layers\t' \
               'Common layers\t'

    def train_row(self, params):
        return f"{os.path.basename(params['train_dir'])}\t" \
               f"{params['eval_loss']}\t" \
               f"{params['eval_acc']}\t" \
               f"{params['auc']}\t" \
               f"{params['avg_precision']}\t" \
               f"{params['train_loss']}\t" \
               f"{params['train_acc']}\t" \
               f"{params['val_loss']}\t" \
               f"{params['val_acc']}\t" \
               f"{[self.get_dict_key(b, self.BRANCHES) for b in params['branches']]}\t" \
               f"{params['batch_size']}\t" \
               f"{self.get_dict_key(params['optimizer'], self.OPTIMIZERS)}\t" \
               f"{params['lr']}\t" \
               f"{self.get_dict_key(params['lr_optim'], self.LR_OPTIMS) if params['lr_optim'] else '-'}\t" \
               f"{params['epochs']}\t" \
               f"{params['no_branches_layers']}\t" \
               f"{params['branches_layers']}\t" \
               f"{params['no_common_layers']}\t" \
               f"{params['common_layers']}\t"

    @staticmethod
    def eval_header():
        return 'Evaluation directory\t' \
               'Evaluation loss\t' \
               'Evaluation accuracy\t' \
               'AUC\t' \
               'Average precision\t' \
               'Model file\t' \
               'Evaluation branches\t' \
               'Window\t' \
               'Window placement\t' \
               'No. classes\t' \
               'Classes\t' \
               'Sequence source type\t' \
               'Sequence source\t' \
               'Strand\t' \
               'Fasta ref.\t' \
               'Conservation ref\t'

    def eval_row(self, params):
        return f"{os.path.basename(params['eval_dir'])}\t" \
               f"{params['eval_loss']}\t" \
               f"{params['eval_acc']}\t" \
               f"{params['auc']}\t" \
               f"{params['avg_precision']}\t" \
               f"{params['model_file']}\t" \
               f"{params['branches']}\t" \
               f"{params['win']}\t" \
               f"{self.get_dict_key(params['win_place'], self.WIN_PLACEMENT)}\t" \
               f"{params['no_klasses']}\t" \
               f"{params['klasses']}\t" \
               f"{self.get_dict_key(params['seq_type'], self.SEQ_TYPES)}\t" \
               f"{params['seq_source']}\t" \
               f"{params['strand']}\t" \
               f"{params['fasta_ref']}\t" \
               f"{params['cons_dir']}\t"

    @staticmethod
    def predict_header():
        return 'Predict directory\t' \
               'Model file\t' \
               'Predict branches\t' \
               'Window\t' \
               'Window placement\t' \
               'No. classes\t' \
               'Classes\t' \
               'Sequence source type\t' \
               'Sequence source\t' \
               'Strand\t' \
               'Fasta ref.\t' \
               'Conservation ref\t'

    def predict_row(self, params):
        return f"{os.path.basename(params['predict_dir'])}\t" \
               f"{params['model_file']}\t" \
               f"{params['branches']}\t" \
               f"{params['win']}\t" \
               f"{self.get_dict_key(params['win_place'], self.WIN_PLACEMENT)}\t" \
               f"{params['no_klasses']}\t" \
               f"{params['klasses']}\t" \
               f"{self.get_dict_key(params['seq_type'], self.SEQ_TYPES)}\t" \
               f"{params['seq_source']}\t" \
               f"{params['strand']}\t" \
               f"{params['fasta_ref']}\t" \
               f"{params['cons_dir']}\t"
```
