## Additional file 1 for "ENNGene: an Easy Neural Network model building tool for Genomics": validators.py

```
import h5py
import os
import subprocess

from .dataset import Dataset
from . import file_utils as f

BRANCHES_REV = {'seq': 'Sequence',
                'cons': 'Conservation score',
                'fold': 'Secondary structure'}


def not_empty_branches(branches):
    warning = 'You must select at least one branch.'
    return warning if len(branches) == 0 else None


def min_two_files(input_files):
    files = [f for f in input_files if not None]
    warning = 'You must provide at least two input files for a classification problem.'
    return warning if len(files) < 2 else None


def uniq_files(input_files):
    files = [f for f in input_files if not None]
    warning = f"The input files must be unique. Currently: {files}."
    return warning if any(files.count(element) > 1 for element in files) else None


def uniq_klasses(klasses):
    warning = f"The class names must be unique. Currently: {klasses}."
    return warning if any(klasses.count(element) > 1 for element in klasses) else None


def is_bed(file, evaluation):
    invalid = False

    if len(file) == 0:
        invalid = True
        warning = 'You must provide the BED file.'
    else:
        if os.path.isfile(file):
            with open(file) as f:
                try:
                    line = f.readline()
                except Exception:
                    invalid = True
                cells = line.split('\t')
                if evaluation:
                    klass = cells.pop(0)
                if len(cells) >= 3:
                    try:
                        int(cells[1])
                        int(cells[2])
                    except Exception:
                        invalid = True
                else:
                    invalid = True
            warning = f"File {file} does not look like valid BED file (tab separated)."
        else:
            invalid = True
            warning = f'Given BED file {file} does not exist.'

    return warning if invalid else None


def is_fasta(file):
    invalid = False

    if len(file) == 0:
        invalid = True
        warning = 'You must provide the FASTA file.'
    else:
        if os.path.isfile(file):
            fasta, zipped = f.unzip_if_zipped(file)
            # the fasta must be unzipped for the bedtools
            if zipped:
                invalid = True
                warning = 'The fasta file must be extracted first. Can not accept compressed file.'
            else:
                try:
                    line1 = f.read_decoded_line(fasta, zipped)
                    line2 = f.read_decoded_line(fasta, zipped)
                    if not line1 or not ('>' in line1) or not line2:
                        invalid = True
                except Exception:
                    invalid = True
                warning = f"File {file} does not look like valid FASTA file."
        else:
            invalid = True
            warning = f'Given FASTA file {file} does not exist.'

    return warning if invalid else None


def is_blackbox(file_path):
    invalid = False
    try:
        dataset = Dataset.load_from_file(file_path)
        # category = dataset.category
        base_columns = ['chrom_name', 'seq_start', 'seq_end', 'strand_sign', 'klass']
        if not all(col in dataset.df.columns for col in base_columns):  # or category != 'blackbox'
            invalid = True
            warning = 'Given file does not seem like valid blackbox dataset. Please check the file.'
        if int(subprocess.check_output(['wc', '-l', file_path]).split()[0]) <= 1:
            invalid = True
            warning = 'Given blackbox dataset file seems to be empty.'
    except Exception:
        invalid = True
        warning = 'Sorry, could not parse given dataset file. Please check the file.'

    return warning if invalid else None


def is_wig_dir(folder):
    # Checks just one random (first found) wig file
    invalid = False

    if len(folder) == 0:
        invalid = True
        warning = 'You must provide the conservation reference directory.'
    else:
        if os.path.isdir(folder):
            files = f.list_files_in_dir(folder, 'wig')
            one_wig = next((file for file in files if 'wig' in file), None)
            if one_wig:
                try:
                    wig_file, zipped = f.unzip_if_zipped(one_wig)
                    line1 = f.read_decoded_line(wig_file, zipped)
                    if not ('fixedStep' in line1 or 'variableStep' in line1) or not ('chrom' in line1):
                        invalid = True
                        warning = f"Provided wig file {one_wig} starts with unknown header."
                    line2 = f.read_decoded_line(wig_file, zipped)
                    float(line2)
                except Exception:
                    invalid = True
                    warning = f"Tried to look at a provided wig file: {one_wig} and failed to properly read it. Please check the format."
            else:
                invalid = True
                warning = "I don't see any WIG file in conservation reference directory."
        else:
            invalid = True
            warning = 'Provided conservation reference directory does not exist.'

    return warning if invalid else None


def is_ratio(string):
    invalid = False
    if len(string) == 0:
        invalid = True
        warning = 'You must provide the split ratio.'
    else:
        if ':' in string:
            parts = string.split(':')
            if len(parts) == 4:
                try:
                    numbers = [float(part) for part in parts]
                    for i, number in enumerate(numbers):
                        warning = 'All numbers in the split ratio must be bigger than zero (only blackbox dataset can be zero).'
                        if (i < 3) & (number <= 0):
                            invalid = True
                        if (i == 3) & (number < 0):
                            # blackbox can be zero but not negative
                            invalid = True
                except Exception:
                    invalid = True
                    warning = 'All parts of the split ratio must be numbers (required format: train:validation:test:blackbox).'
            else:
                invalid = True
                warning = 'Invalid format of the split ratio (required format: train:validation:test:blackbox).'
        else:
            invalid = True
            warning = 'Invalid format of the split ratio (required format: train:validation:test:blackbox).'

    return warning if invalid else None


def is_dataset_dir(folder):
    invalid = False

    if len(folder) == 0:
        invalid = True
        warning = 'You must provide a folder with the preprocessed files.'
    else:
        if os.path.isdir(folder):
            param_files = [file for file in os.listdir(folder) if (file == 'parameters.yaml') and
                           (os.path.isfile(os.path.join(folder, file)))]
            if len(param_files) == 1:
                files = f.list_files_in_dir(folder, 'zip')
                dataset_files = {'train': [], 'validation': [], 'test': [], 'blackbox': []}
                for file in files:
                    category = next((category for category in dataset_files.keys() if category in os.path.basename(file)), None)
                    if category: dataset_files[category].append(file)
                for category, files in dataset_files.items():
                    if category == 'blackbox': continue  # the blackbox dataset is optional
                    if len(files) != 1:
                        invalid = True
                        warning = 'Each category (train, test, validation) must be represented by exactly one preprocessed file in the given folder.'
                    else:
                        if files[0] != f'{category}.zip':
                            invalid = True
                            warning = ''
            else:
                invalid = True
                warning = 'Sorry, there is no parameters.yaml file in the given folder. Make sure to provide the whole ' \
                          'datasets folder (not just the one with final datasets).'
        else:
            invalid = True
            warning = 'Given folder with preprocessed files does not exist.'

    return warning if invalid else None


def is_full_dataset(file_path, branches):
    invalid = False

    if len(file_path) == 0:
        invalid = True
        warning = 'You must provide the mapped file.'
    else:
        try:
            dataset = Dataset.load_from_file(file_path)
            category = dataset.category
            base_columns = ['chrom_name', 'seq_start', 'seq_end', 'strand_sign', 'klass']
            if category or not all(col in dataset.df.columns for col in base_columns):
                invalid = True
                warning = 'Given mapped file does not contain necessary data. Please check the file.'
            if not all(col in dataset.df.columns for col in branches):
                invalid = True
                warning = 'Given mapped file does not contain selected branches.'
        except Exception:
            invalid = True
            warning = 'Sorry, could not parse given mapped file. Please check the file.'

    return warning if invalid else None


def not_empty_chromosomes(chromosomes_list):
    invalid = False
    for category, chromosomes in chromosomes_list:
        if category == 'blackbox': continue
        if not chromosomes:
            invalid = True
            warning = 'You must select at least one chromosome per each category (only blackbox dataset can remain empty).'

    return warning if invalid else None


def is_model_file(file_path):
    invalid = False
    if len(file_path) == 0:
        invalid = True
        warning = 'You must provide the hdf5 file with a trained model.'
    else:
        if os.path.isfile(file_path):
            if not h5py.is_hdf5(file_path):
                invalid = True
                warning = 'Given file does not seem to be a valid model (requires hdf5 format).'
        else:
            invalid = True
            warning = 'Given file with model does not exist.'

    return warning if invalid else None


def is_multiline_text(text):
    invalid = False
    sequences = text.strip().split('\n')
    if len(text) == 0 or len(sequences) == 0:
        invalid = True
        warning = 'You must provide at least one sequence to be classified.'

    return warning if invalid else None
```
