## Additional file 1 for "ENNGene: an Easy Neural Network model building tool for Genomics": ENNGene_installation.py

```
from pathlib import Path 
from subprocess import SubprocessError, check_output
import subprocess
import sys


# Check anaconda and install if needed
print('Checking Anaconda \n')
conda_check = subprocess.run('conda --version', shell=True, capture_output=True)
if len(conda_check.stdout) > 0:
    version = conda_check.stdout.decode('ascii')
    print(f'Anaconda is already installed! Version: {version}')
else:
    print('Cannot continue without Anaconda installed in your system. Read the Docs. Stopping installation. \n', 'red')
    sys.exit(0)


# Install and import python package termcolor
print('Installing python package termcolor to conda env, that\'s necessary for this installation script. \n')
try:
    termcolor_pkg_install = subprocess.run('conda install termcolor', shell=True, check=True)
    print('\nPackage successfully installed. \n')
    from termcolor import colored
except subprocess.CalledProcessError:
    print('\nOups, something went wrong, check the message above!')
    sys.exit(0)


def yes_or_no(question, color='blue'):
    yes = {'yes','y', 'ye'}
    no = {'no','n'}
    answer = input(colored(f'{question}', f'{color}')).lower()
    if answer in yes:
        return True
    elif answer in no:
        return False
    else:
        print(colored('Please respond with "yes/no" or "y/n". \n', 'yellow'))
        yes_or_no(question)


def perform_and_check(cmd, color='red', answer='Oups, something went wrong, check the message above! \n'):
    try:
        subprocess.run(cmd, shell=True, check=True)
        return True
    except subprocess.CalledProcessError:
        print(colored(f'{answer}', color))
        return False


def empty_directory(directory_path):
    try:
        empty_dir = any(directory_path.iterdir())
        if empty_dir is True:
            return True
        else:
            return False
    except FileNotFoundError:
        return False


def valid_directory(question, default_path):
    user_path = input(colored(f'{question}', 'blue'))
    user_dir = Path(user_path)
    if user_dir.exists():
        if empty_directory(user_dir) is False:
            return user_dir
        else:
            valid_directory('Chosen directory is not empty. Please, choose another directory: ', default_path)
    else:
        non_existing_directory = yes_or_no(f'Provided directory ({user_dir}) doesn\'t exist. Do you want to create it? (y/n) If you don\'t, default directory will be used instead: ')
        if non_existing_directory is True:
            try:
                user_dir.mkdir(parents=True, exist_ok=True)
                if user_dir.exists():
                    return user_dir
            except PermissionError:
                print(colored('You don\'t have permission to write to this directory. Check your path and provide it again.', 'yellow'))
                valid_directory(question, default_path)
            else:
                print(colored('Invalid path. Please, fill in correct absolute path to the directory.', 'yellow'))
                valid_directory(question, default_path)
        else:
            user_dir = default_path
            return user_dir


# Set up $HOME directory 
home = Path.home()


# Update conda to the latest version
print(colored('Updating Anaconda \n', 'blue'))
update_conda = perform_and_check('conda update --yes conda && conda update --yes anaconda')
if update_conda is True:
    print(colored('Update complete! \n', 'green'))
else:
    print(colored('Updating conda has failed. \n\
You can update conda manually later with following command: conda update --yes conda && conda update --yes anaconda \n\
Installation continues.. \n', 'yellow'))


# Extract anaconda directory
conda_dir = subprocess.run('which conda', shell=True, capture_output=True, check=True).stdout.decode('ascii')[:-6]


# Check and install git, if necessary
print(colored('Checking Git \n', 'blue'))
git_check = subprocess.run('git --version', shell=True, capture_output=True)
if len(git_check.stdout) > 0:
    version = git_check.stdout.decode('ascii')
    print(colored(f'Git is already installed! Version: {version}', 'green'))
else:
    git_status = yes_or_no('Git is not installed on your system and it is necessary to clone enngene.\n\
Do you agree with installing Git to your system? (y/n) ')
    if git_status is True:
        print(colored('Installing Git \n', 'blue'))
        git_installer = perform_and_check('conda install -c anaconda git')
        if git_installer is True:
            print(colored('\nGit installation has successfully finished! \n', 'green'))
        else:
            print(colored('Git installation has failed. Cannot continue without installing Git. Stopping instalation.. \n', 'red'))
            sys.exit(0)
    else:
        print(colored('Cannot continue without installing Git. You can install Git manually using following command: conda install -c anaconda git \n\
Stopping instalation. \n', 'red'))
        sys.exit(0)


# Download ENNgene to HOME (default) or users directory (requires username and password to Gitlab)
enngene_dir_status = yes_or_no(f'ENNGene will be downloaded to your /home directory: {home}/enngene \n\
Do you want to change the directory? (y/n) ')
if enngene_dir_status is True:
    user_dir = valid_directory(f'Your /home directory is: {home} \n\
Specify absolute path to the existing directory, where ENNGene should be placed (e.g. {home}/Documents/): \n', home)
    enngene_dir = user_dir / 'enngene'
    try:
        empty_dir = any(enngene_dir.iterdir())
        if empty_dir is True:
            non_empty_default_dir_status = yes_or_no('Default directory is not empty. Do you want to delete its content (necessary for successful cloning)? (y/n): ', color='yellow')
            if non_empty_default_dir_status is True:
                delete_directory_content = perform_and_check(f'rm -rf {enngene_dir}')
                if delete_directory_content is True:
                    print(colored('\nDirectory has been successfully removed. \n', 'green'))
                else:
                    print(colored('Cannot set up directory for cloning ENNGene inside. Stopping installation.. \n', 'red'))
                    sys.exit(0)
            else:
                print(colored('Cannot set up directory for cloning ENNGene inside. Stopping installation.. \n', 'red'))
                sys.exit(0)
    except FileNotFoundError:
        pass
    else:
        pass
else:
    enngene_dir = home / 'enngene'
    try:
        empty_dir = any(enngene_dir.iterdir())
        if empty_dir is True:
            non_empty_default_dir_status = yes_or_no('Default directory is not empty. Do you want to delete its content (necessary for successful cloning)? (y/n): ', color='yellow')
            if non_empty_default_dir_status is True:
                delete_directory_content = perform_and_check(f'rm -rf {enngene_dir}')
                if delete_directory_content is True:
                    print(colored('\nDirectory has been successfully removed. \n', 'green'))
                else:
                    print(colored('Cannot set up directory for cloning ENNGene inside. Stopping installation.. \n', 'red'))
                    sys.exit(0)
            else:
                print(colored('Cannot set up directory for cloning ENNGene inside. Stopping installation.. \n', 'red'))
                sys.exit(0)
    except FileNotFoundError:
        pass
    else:
        pass


print(colored(f'Downloading ENNGene from the repository to {enngene_dir} \n', 'blue'))
try:
    subprocess.run(f'git clone --single-branch --branch master https://github.com/ML-Bioinfo-CEITEC/ENNGene {enngene_dir}', shell=True, check=True)
    print(colored(f'\nCloning ENNGene into the chosen directory has successfully finished! \n\
You may find ENNGene in {enngene_dir} \n', 'green'))
except subprocess.CalledProcessError as err:
    print(colored('Cannot clone ENNGene from GitHub. You can check the problem and repeat the installation process. Stopping installation.. \n', 'red'))
    sys.exit(0)
else:    
    pass


# Create conda env and install all dependencies
print(colored('Creating conda environment: enngene; installing all dependencies \n', 'blue'))
conda_env = perform_and_check(cmd=f'conda env create -f {enngene_dir}/environment.yml', 
                              answer='Conda environment named "enngene" already exists. Updating the env..\n',
                              color='yellow')
if conda_env is True:
    print(colored('Complete! \nYou can activate env using cmd: conda activate enngene \n', 'green'))
else:
    conda_env_update = perform_and_check(cmd=f'conda env update --name enngene -f {enngene_dir}/environment.yml',
                                        color='yellow')
    if conda_env_update is True:
        print(colored('Complete! \nYou can activate env using cmd: conda activate enngene \n', 'green'))
    else:
        conda_env_remove_status = yes_or_no(f'There is problem with updating existing enngene environment. Do you agree with removing old environment \
and replacing it with a new one? (y/n) ')
        if conda_env_remove_status is True:
            conda_remove_env = perform_and_check(cmd=f'conda env remove --name enngene')
            if conda_remove_env is True:
                print(colored('Complete! \nOld enngene environment was successfully removed.\n', 'green'))
                print(colored('Creating conda environment: enngene; installing all dependencies \n', 'blue'))
                new_conda_env = perform_and_check(cmd=f'conda env create -f {enngene_dir}/environment.yml',
                                                  color='red')
                if new_conda_env is True:
                    print(colored('Complete! \nYou can activate env using cmd: conda activate enngene \n', 'green'))
                else:
                    print(colored(f'Environment set up has failed. \n\
Use the following command to create environment manually later: conda env create -f {enngene_dir}/environment.yml \n\
If you only need to update enngene environment, use following command: conda env update --name engene -f {enngene_dir}/environment.yml \n\
Installation continues.. \n', 'yellow'))
        else:
            print(colored(f'Cannot continue without removing existing enngene environment.. \n\
Use the following commands to remove and re-create environment manually later: \n\
\t\t\t\t conda env remove --name enngene \n\
\t\t\t\t conda env create -f {enngene_dir}/environment.yml \n', 'yellow'))
        

# Create and activate ENNGene launcher
print(colored('Creating ENNGene launcher \n', 'blue'))

tmp_launcher_path = enngene_dir / 'launcher.txt'
final_launcher_path = enngene_dir / 'launcher.desktop'

with tmp_launcher_path.open('w', encoding='utf-8') as launcher:
    launcher.write(
        f'#!/bin/bash\n\
        [Desktop Entry]\n\
        Name=ENNGene\n\
        Exec=bash -c "source {conda_dir}activate enngene && streamlit run {enngene_dir}/enngene/enngene.py; read"\n\
        Icon=utilities-terminal\n\
        Terminal=true\n\
        Type=Application\n\
        Categories=Application;')

make_launcher_executable_1 = perform_and_check(f'mv {tmp_launcher_path} {final_launcher_path}')
make_launcher_executable_2 = perform_and_check(f'chmod a+x {final_launcher_path}')
if make_launcher_executable_1 and make_launcher_executable_2 is True:
    print(colored(f'Complete! You can find the launcher at {final_launcher_path} \n', 'green'))
elif make_launcher_executable_1 is True and make_launcher_executable_2 is False:
    print(colored(f'Cannot set up privileges for the launcher. \n\
Please, check your permissions and set up privileges of the launcher later with the following command: chmod a+x {final_launcher_path} \n', 'yellow'))
else:
    print(colored('Creating the launcher has failed. Installation has finished, BUT without functional launcher. \n', 'red'))
    print(colored(f'You can check the problem and repeat the installation or run the ENNGene using following commands: \n\
\tconda activate enngene \n\
\tstreamlit run {enngene_dir}/enngene/enngene.py \n', 'yellow'))
    sys.exit(0)


# Ask user whether he wants to create a desktop launcher; create it if so
desktop_launcher_status = yes_or_no('Do you want to make a copy of a launcher to your desktop? (y/n) ')
if desktop_launcher_status is True:
    print(colored('Making copy of launcher to your Desktop \n', 'blue'))
    make_desktop_launcher_executable_1 = perform_and_check(f'cp {enngene_dir}/launcher.desktop {home}/Desktop/')
    make_desktop_launcher_executable_2 = perform_and_check(f'chmod a+x {home}/Desktop/launcher.desktop')
    if make_desktop_launcher_executable_1 and make_desktop_launcher_executable_2 is True:
        print(colored(f'Complete! You can find the desktop launcher at {home}/Desktop/launcher.desktop \n', 'green'))
    elif make_launcher_executable_1 is True and make_launcher_executable_2 is False:
        print(colored(f'Cannot set up privileges for the desktop launcher. \n\
Please, check your permissions and set up privileges of the launcher later with the following command: chmod a+x {home}/Desktop/launcher.desktop \n', 'yellow'))
    else:
        print(colored('Creating the desktop launcher has failed. Installation has finished, BUT without functional desktop launcher. \n', 'red'))
        print(colored(f'You can check the problem and repeat the installation or run the ENNGene using launcher in the ENNGene folder or by running following commands: \n\
    \tconda activate enngene \n\
    \tstreamlit run {enngene_dir}/enngene/enngene.py \n', 'yellow'))
        sys.exit(0)
else:
    print(colored(f'\nSkipping desktop launcher. \n\
You can create a desktop launcher later by using following commands: \n\
\t cp {enngene_dir}/launcher.desktop {home}/Desktop/ \n\
\t chmod a+x {home}/Desktop/launcher.desktop \n', 'green'))
    pass


# Finish the installation
print(colored('Installation has finished! \n', 'green'))
```
