## Additional file 1 for "ENNGene: an Easy Neural Network model building tool for Genomics": environment.yml

```
name: enngene
channels:
  - anaconda
  - bioconda
  - defaults
dependencies:
  - _libgcc_mutex=0.1=main
  - altair=3.2.0=py36_0
  - attrs=19.3.0=py_0
  - bedtools=2.29.2=hc088bd4_0
  - blas=1.0=mkl
  - bzip2=1.0.8=h7b6447c_0
  - ca-certificates=2020.12.8=h06a4308_0
  - cairo=1.14.12=h8948797_3
  - certifi=2020.12.5=py36h06a4308_0
  - cycler=0.10.0=py36_0
  - dbus=1.13.12=h746ee38_0
  - expat=2.2.6=he6710b0_0
  - fontconfig=2.13.0=h9420a91_0
  - freetype=2.9.1=h8a8886c_1
  - fribidi=1.0.5=h7b6447c_0
  - glib=2.56.2=hd408876_0
  - graphite2=1.3.13=h23475e2_0
  - graphviz=2.40.1=h21bd128_2
  - gst-plugins-base=1.14.0=hbbd80ab_1
  - gstreamer=1.14.0=hb453b48_1
  - h5py=2.10.0=py36h7918eee_0
  - harfbuzz=1.8.8=hffaf4a1_0
  - hdf5=1.10.4=hb1b8bf9_0
  - icu=58.2=h9c2bf20_1
  - importlib_metadata=1.5.0=py36_0
  - intel-openmp=2019.4=243
  - joblib=0.17.0=py_0
  - jpeg=9b=h024ee3a_2
  - jsonschema=3.2.0=py36_0
  - kiwisolver=1.1.0=py36he6710b0_0
  - libedit=3.1.20181209=hc058e9b_0
  - libffi=3.2.1=hd88cf55_4
  - libgcc-ng=9.1.0=hdf63c60_0
  - libgfortran-ng=7.3.0=hdf63c60_0
  - libpng=1.6.37=hbc83047_0
  - libstdcxx-ng=9.1.0=hdf63c60_0
  - libtiff=4.1.0=h2733197_0
  - libuuid=1.0.3=h1bed415_2
  - libxcb=1.13=h1bed415_1
  - libxml2=2.9.9=hea5a465_1
  - markupsafe=1.1.1=py36h7b6447c_0
  - matplotlib=3.1.1=py36h5429711_0
  - mkl=2019.4=243
  - mkl-service=2.3.0=py36he904b0f_0
  - mkl_fft=1.0.15=py36ha843d7b_0
  - mkl_random=1.1.0=py36hd6b4f25_0
  - ncurses=6.1=he6710b0_1
  - numpy-base=1.18.1=py36hde5b4d6_1
  - openssl=1.1.1i=h27cfd23_0
  - pango=1.42.4=h049681c_0
  - pcre=8.43=he6710b0_0
  - pixman=0.38.0=h7b6447c_0
  - pydot=1.3.0=py36_0
  - pyparsing=2.4.6=py_0
  - pyqt=5.9.2=py36h05f1152_2
  - python=3.6.8=h0371630_0
  - pytz=2019.3=py_0
  - qt=5.9.7=h5867ecd_1
  - readline=7.0=h7b6447c_5
  - scikit-learn=0.23.2=py36h0573a6f_0
  - scipy=1.5.2=py36h0b6359f_0
  - seaborn=0.11.0=py_0
  - setuptools=45.1.0=py36_0
  - sip=4.19.8=py36hf484d3e_0
  - six=1.14.0=py36_0
  - sqlite=3.30.1=h7b6447c_0
  - threadpoolctl=2.1.0=pyh5ca1d4c_0
  - tk=8.6.8=hbc83047_0
  - toolz=0.10.0=py_0
  - viennarna=2.4.14=py36h8b12597_0
  - wheel=0.34.1=py36_0
  - xz=5.2.4=h14c3975_4
  - yaml=0.1.7=h96e3832_1
  - zlib=1.2.11=h7b6447c_3
  - zstd=1.3.7=h0b5b093_0
  - pip:
    - absl-py==0.8.1
    - argh==0.26.2
    - astor==0.8.0
    - backcall==0.2.0
    - base58==1.0.3
    - bleach==3.1.5
    - blinker==1.4
    - boto3==1.14.31
    - botocore==1.17.31
    - cachetools==4.1.1
    - chardet==3.0.4
    - click==7.0
    - decorator==4.4.1
    - defusedxml==0.6.0
    - docutils==0.15.2
    - entrypoints==0.3
    - enum-compat==0.0.3
    - future==0.18.2
    - gast==0.2.2
    - gitdb==4.0.5
    - gitpython==3.1.11
    - google-auth==1.20.0
    - google-auth-oauthlib==0.4.1
    - google-pasta==0.1.8
    - grpcio==1.25.0
    - idna==2.8
    - importlib-metadata==0.23
    - ipykernel==5.3.4
    - ipython==7.16.1
    - ipython-genutils==0.2.0
    - ipywidgets==7.5.1
    - jedi==0.17.2
    - jinja2==2.10.3
    - jmespath==0.9.4
    - jupyter-client==6.1.6
    - jupyter-core==4.6.3
    - keras-applications==1.0.8
    - keras-preprocessing==1.1.0
    - markdown==3.1.1
    - mistune==0.8.4
    - more-itertools==7.2.0
    - nbconvert==5.6.1
    - nbformat==5.0.7
    - notebook==6.0.3
    - numpy==1.17.4
    - oauthlib==3.1.0
    - opt-einsum==3.1.0
    - packaging==20.4
    - pandas==1.1.1
    - pandocfilters==1.4.2
    - parso==0.7.1
    - pathtools==0.1.2
    - pexpect==4.8.0
    - pickleshare==0.7.5
    - pillow==6.2.1
    - pip==20.3.3
    - prometheus-client==0.8.0
    - prompt-toolkit==3.0.5
    - protobuf==3.12.4
    - ptyprocess==0.6.0
    - pyarrow==1.0.0
    - pyasn1==0.4.8
    - pyasn1-modules==0.2.7
    - pydeck==0.4.0
    - pygments==2.6.1
    - pyrsistent==0.15.5
    - python-dateutil==2.8.0
    - pyyaml==5.1.2
    - pyzmq==19.0.1
    - requests==2.22.0
    - requests-oauthlib==1.3.0
    - rsa==4.0
    - s3transfer==0.3.3
    - send2trash==1.5.0
    - smmap==3.0.4
    - streamlit==0.72.0
    - style==1.1.0
    - tensorboard==2.0.1
    - tensorflow-estimator==2.0.1
    - tensorflow-gpu==2.0.0
    - termcolor==1.1.0
    - terminado==0.8.3
    - testpath==0.4.4
    - toml==0.10.0
    - tornado==5.1.1
    - tqdm==4.61.1
    - traitlets==4.3.3
    - tzlocal==2.0.0
    - update==0.0.1
    - urllib3==1.25.7
    - validators==0.14.0
    - watchdog==0.9.0
    - wcwidth==0.2.5
    - webencodings==0.5.1
    - werkzeug==0.16.0
    - widgetsnbextension==3.5.1
    - wrapt==1.11.2
    - zipp==0.6.0
```
