## Additional file 1 for "ENNGene: an Easy Neural Network model building tool for Genomics": README.md

## ENNGene 
ENNGene is an application that simplifies the local training of custom Convolutional Neural Network models on Genomic data 
via an easy to use Graphical User Interface. ENNGene allows multiple streams of input information, including sequence, 
evolutionary conservation, and secondary structure, and includes utilities that can perform needed preprocessing steps, 
allowing simplified input such as genomic coordinates. 
ENNGene deals with all steps of training and evaluation of Convolutional Neural Networks for genomics, 
empowering researchers that may not be experts in machine learning to perform this powerful type of analysis.

> We are preparing a video tutorial for an even easier start with the ENNGene application!

### Installation
To install ENNGene, you need the following prerequisites:
- [Python 3](https://www.python.org/) - Python language is usually already available with the OS
- [Anaconda](https://www.anaconda.com/products/individual) - Python package manager for safe installation 
- Web browser

Application and installation scripts were tested on the following systems:
- Ubuntu 18.04.5 LTS, 64-bit, GNOME Version 3.28.2, with Google Chrome browser (Version 86.0.4240.198 (Official Build) (64-bit))
- Ubuntu 20.04.1 LTS, 64-bit, GNOME Version 3.36.8, with Google Chrome browser (Version 87.0.4280.88 (Official Build) (64-bit))
- Ubuntu 20.04.1 LTS, 64-bit, GNOME version 3.36.8, with Firefox Browser 84.0 64-bit.

#### "One-click" installation
For 'one-click' installation, please copy the following lines into the terminal:

`curl -fsSL https://raw.githubusercontent.com/ML-Bioinfo-CEITEC/ENNGene/master/installation.py > ENNGene_installation.py`

`python3 ENNGene_installation.py`

This will download the repository, and create a new conda environment where all the necessary packages will be installed.

User has the opportunity to create a desktop launcher for the app during the "one-click" installation.

**Note** that the launcher will work only on the Ubuntu OS. 
For the Ubuntu 18.04 LTS, there can be two fully functional launchers (first in the app folder, second on the desktop) right after the installation. 
If you have the OS Ubuntu 20.04 LTS, only the desktop launcher will work due to the nature of the system, and the launcher needs to be activated. 
To activate the launcher, right-click on the launcher and check the option `Allow launching`.

If the installation does not work for you, please follow the steps for the manual installation below.

#### Manual installation
If you do not have [Anaconda](https://www.anaconda.com/distribution/) installed on your computer, please do so first. 
- Download the latest release from [the repository](https://github.com/ML-Bioinfo-CEITEC/ENNGene/releases)
- Unzip the directory `tar -xf enngene.tar.gz`
- Go to the project directory `cd enngene`
- Recreate environment from yml file `conda env create -f environment.yml`
- Activate the environment`conda activate enngene`
- Run the app `cd enngene` and `streamlit run enngene.py`

### Implementation
ENNGene is built atop TensorFlow, one of the most popular Deep Learning frameworks. 
It can run on either CPU or GPU, offering a considerable speed-up when training larger CNNs.
ENNGene accepts BED genomic interval files corresponding to a user-determined genome or transcriptome reference.
Classifiers can be built for two or more classes.
Full reproducibility of the process is ensured by logging all the user’s input and exporting it as a yaml file that can be loaded in ENNGene for future runs.
The application consists of three consecutive modules, each module performing a specific task.
 
To select a task browse the select box in the side panel. 

`Output folder` There is a subfolder created for each task in the given output folder (thus you can use the same path for all the tasks you run). 
After running a task, several files will be exported to the task subfolder.
 * Parameters.yaml file - contains all the parameters set by the user.
 The yaml file can be imported in a future run to reproduce the set up.
 * Log file - contains logged user input, warnings, errors etc. 
 * Parameters.tsv file - a tsv table with one row per run, shared across the task.
   You can easily manage and compare results across the specific task with different parameters' setup.
 * Other task-specific files.

The ENNGene application uses the [Streamlit framework](https://www.streamlit.io/) that is still in its early stages of development.
Currently, all input files (or folders) must be defined by an absolute path.

> **Due to the nature of the Streamlit framework, it is strongly recommended to fill-in the input fields top to bottom to avoid resetting already specified options.**

Hopefully, this behavior will be removed with the framework update.

##### Error handling
All the user input is verified by the application for its correctness, to the possible extent. 
When an incorrect input is given (e.g. file with a wrong format, non-existent folder etc.), the application gives a warning and exits before starting the task itself.
This way, you can save a lot of time, instead of debugging the process yourself. 

If you see an unspecified warning 'Unexpected error occurred in the application.', please check the provided log file for more details.
Sometimes this happens as a result of a glitch in the Streamlit framework, and simple reloading the page with the application solves the issue.

> **If you get an 'Unexpected error', try first reloading the page with the application. 
>This sometimes happens due to a glitch in the Streamlit framework, and reloading the page solves the issue.** 

#### 1 Preprocessing
In the first module, data is preprocessed into a format convenient for CNN input.

`Use already preprocessed file` Check this option to save preprocessing time if you already have files prepared from the previous run. 
Using a mapped file, you can still change the datasets' size, or redistribute data across categories.

`Branches` You may select one or more input types engineered from the given interval files.
Each input type later corresponds to a branch in the neural network.
 * Sequence – one-hot encoded RNA or DNA sequence. Requires reference genome/transcriptome in a fasta file.
 * Secondary structure – computed by [ViennaRNA](https://www.tbi.univie.ac.at/RNA/) package, one-hot encoded. (Also requires the reference genome in fasta file).
 * Conservation score – counted based on the user provided reference file/s. This option is the most time-consuming, we advise to use it judiciously.

`Apply strand` Choose to apply (if available) or ignore strand information.

`Window size` All the samples prepared for the neural network must be of the same length, defined by the window size.
Longer sequences get shortened, while short sequences are completed based on the reference.
Both is done by randomly placing a window of selected size on the original sequence.

`Window placement` Choose a way to place the window upon the sequence:
 * Randomized 
 * Centered

`Path to the reference fasta file` File containing reference genome/transcriptome. 
Required when Sequence or Secondary structure branch is selected.

`Path to folder containing reference conservation files` Required when Conservation score branch is selected.'Path to folder containing reference conservation files'

`Number of CPUs` You might assign multiple CPUs for the computation of the secondary structure.

##### Input Coordinate Files
`Number of input files` There can be an arbitrary number of input files in BED format (two at minimum).
Each input file corresponds to one class for the classification. Class name is based on the file name.

`File no. 1, File no. 2, ...` Enter an absolute path for each interval file separately.

##### Dataset Size Reduction
`Classes to be reduced` Number of samples in the selected class/es can be reduced to save the computing resources when training larger networks.
 
`Target dataset size` Define a target size per each dataset selected to be reduced.
Input a decimal number if you want to reduce the sample size by a ratio (e.g. 0.1 to get 10%), or an integer if you wish to select final dataset size (e.g. 5000 if you want exactly 5000 samples).
A hint showing a number of rows in the original input file is displayed at the end. 

*Note: Samples are randomly shuffled across the chromosomes before the size reduction. 
If you split the dataset by chromosomes after reducing its size, make sure all the classes in all the categories (train, test, etc.) 
contain at least some data, as some small chromosomes might get fully removed.*

##### Data Split
Supplied data must be split into training, validation and testing datasets.

*Note: The testing dataset is used for a direct evaluation of the trained model. 
If you train multiple models on the same datasets, you might want to keep a 'blackbox' dataset for a final evaluation.*

`Random` Data are split into the categories randomly across the chromosomes, based on the given ratio.

`Target ratio` Defines the ratio of the number of samples between the categories. Required format: train:validation:test:blackbox.

`By chromosomes` Specific chromosomes might be selected for each category.
To use this option, a fasta file with reference genome/transcriptome must be provided (the same one required for the sequence and secondary structure branches).
List of available chromosomes is then inferred from the provided reference (scaffolds are ignored) - that may take up to few minutes.

*Note: When selecting the chromosomes for the categories Streamlit will issue a warning 'Running read_and_cache(...).'. 
You may disregard that and continue selecting the chromosomes. 
Although, if your machine cannot handle it, and the process gets stuck, your input might get nullified. 
In that case, you will want to wait until the warning disappears.*

`Run` After all the parameters are set and selected, press the run button. 
Depending on the amount of data, selected options, and the hardware available, the preprocessing might take several minutes to hours. 

Files with preprocessed datasets are exported to the 'datasets' subfolder at the `output folder` defined at the beginning.  

#### 2 Training
In the second module, neural network architecture as well as the hyperparameters are set, and the model is trained using the data preprocessed in the first module.   

`Datasets folder` Define a path to the folder containing datasets created by the Preprocessing module. 

`Branches` Select the branches you want the model to be composed of. 
You might choose only from the branches preprocessed in the first module.

`Output TensorBoard log files` For more information see the [official site](https://www.tensorflow.org/tensorboard).

##### Training Options

`Batch size` Number of training samples utilized in one iteration. 

`No. of training epochs` An epoch is one complete pass through the training data. There can be an arbitrary number of training epochs.

`Apply early stopping` A regularization technique to avoid overfitting when training for too many epochs. 
The model will stop training if the validation loss does not decrease for more than 0.01 (min_delta) during 10 training epochs (patience). 

`Optimizer` Select an optimizer. Available options: 
 * Stochastic Gradient Descent ([SGD](https://www.tensorflow.org/api_docs/python/tf/keras/optimizers/SGD)) - 
 parameters are set as follows: momentum = 0.9, [nesterov](http://proceedings.mlr.press/v28/sutskever13.pdf) = True.
 * [RMSprop](https://www.tensorflow.org/api_docs/python/tf/keras/optimizers/RMSprop) - parameters are set to default.
 * [Adam](https://www.tensorflow.org/api_docs/python/tf/keras/optimizers/Adam) - parameters are set to default. Implements [Adam algorithm](https://arxiv.org/abs/1412.6980).

`Learning rate options` Applies only for the SGD optimizer. Available options:
 * Use fixed learning rate - applies the same learning rate value throughout the whole training.
 * Use [learning rate scheduler](https://www.tensorflow.org/api_docs/python/tf/keras/callbacks/LearningRateScheduler) - 
 gradually decreases learning rate from the given value.
 * Apply [one cycle policy](https://arxiv.org/abs/1506.01186) - uses the learning rate value as a maximum.
 The implementation for Keras is taken from [here](https://github.com/titu1994/keras-one-cycle), originally ported from the [fast.ai project](https://github.com/fastai/fastai).

`Learning rate` Corresponds to the step size during the gradient descent.

`Metric` Choose a metric. Available options: accuracy.

`Loss function` Choose a loss function. Available options: categorical crossentropy.

##### Network Architecture

The last section determines the network architecture.
You may define architecture for each of the selected branches separately, as well as for the common part of the network following the branches' concatenation.

`Number of layers` First set a number of layers per each part (branches or common part of the network).

`Layer type` Types available for the branches: 
 * [Convolution layer](https://www.tensorflow.org/api_docs/python/tf/keras/layers/Conv1D)
 <!-- * [Locally connected 1D layer](https://www.tensorflow.org/api_docs/python/tf/keras/layers/LocallyConnected1D)--> 
 
 Types available for the connected part of the neural network:
 * [Convolution layer](https://www.tensorflow.org/api_docs/python/tf/keras/layers/Conv1D)
 * [LSTM](https://www.tensorflow.org/api_docs/python/tf/keras/layers/LSTM)
 * [GRU](https://www.tensorflow.org/api_docs/python/tf/keras/layers/GRU)  
 * [Dense layer](https://www.tensorflow.org/api_docs/python/tf/keras/layers/Dense)
<!--* [Locally connected 1D layer](https://www.tensorflow.org/api_docs/python/tf/keras/layers/LocallyConnected1D)-->

Different types of layers are not combinable. 
E.g. when LSTM layer is chosen, it can be followed either by another LSTM layer, or a Dense layer only.
However, multiple layers of the same type can be stacked. 

`Show advanced options` If checked, you may set options specific per layer type. If not, the defaults will apply.

*Note: Due to the nature of the Streamlit framework, it is necessary to keep the checkbox checked for the options to be applied.
If it is unchecked, the options get reset.*

Common options are:
 * `Batch normalization` Applies [batch normalization](https://arxiv.org/abs/1502.03167) for the layer if checked.
 * `Dropout rate` Select a [dropout](https://jmlr.org/papers/v15/srivastava14a.html) rate. 

Options available for Convolutional layers:  <!-- and Locally connected 1D layers:-->
* `Number of filters` The number of output filters in the convolution.
* `Kernel size` Specifies the length of the 1D convolutional window.

Options available for Dense layer, GRU and LSTM:
* `Number of units` Dimensionality of the output space.

Option available for GRU and LSTM:
* `Bidirectional` Apply a bidirectional wrapper on a recurrent layer.

*Note: The softmax activation function is used for the last layer.*

`Run` When all is set, press the run button to start training the model.
The training time depends on many variables (dataset size, network architecture, number of epochs, hardware available, etc.).
You can monitor the progress on the chart indicating metric and loss function values.

Resulting model and other files are exported to the 'training' subfolder in the selected `output folder`. 

#### 3 Evaluation & Prediction
In the last two modules, a trained model can be evaluated on sequences with a know class or used to classify novel, unseen data.
As the parameters for the two modules are mostly overlapping, they will be covered in this section together.
Sequences provided to be classified are preprocessed similar to the first module for the purpose of the CNN. 

##### Model
You can either use a model trained with the ENNGene application, or any custom trained model.

`Use a model trained by the ENNGene application` Preferred option. When selected this option, you must provide:
 * `Training folder containing the model (hdf5 file)` Except the hdf5 file with the trained model, the folder must contain the parameters.yaml file logged when training the model. 
 Form that the parameters necessary for sequence preprocessing are read, and displayed below the field after that. 

`Use a custom trained model` When using model trained otherwise than through the application, necessary parameters must be provided separately.
When selected this option, you must provide:
 * `Trained model (hdf5 file)` Path to the hdf5 file with the trained model.
 * `Window size` The size of the window must be the same as when used ofr the training the given model.
 * `Number of classes` Number must be the same as the number of classes used for training the given model.
 * `Class labels` Provide names of the classes for better results interpretation. 
 The order of the classes must be the same as when encoding them for training the given model.
 * `Branches` Define branches used in the given model's architecture. Available options are: Sequence, Secondary structure, Conservation score. 
 
##### Sequences

You can provide the input sequences you wish to classify in following formats:
 * `BED file` - When used for Evaluation, a column containing class information must be inserted at the beginning of the file.
   I.e. the first column of the file must contain the name of the class per each sequence. Class names must correspond to those used when training the model.
 * `FASTA file` - When used for Evaluation, a klass name must be provided as a last part of the header, separated by a space. E.g. '>chr16:655478-655578 FUS_positives'.  
 * `Text input` - Available for Prediction. Paste one sequence per line.
 * `Blackbox dataset` - Available for Evaluation. Provide a path to the blackbox dataset file exported by the Preprocess module.
Dataset should come from the same data as those used for training the model, or the parameters must match at least (e.g. class names, window size, branches...).  

*Note: If the Conservation score branch is applied, only files in BED format are accepted, as the coordinates are necessary to get the score.*

`Window placement` Choose a way to place the window upon the sequence:
* Randomized
* Centered

`BED file` When providing the sequences via an interval file, following must be specified:
 * `Path to the BED file containing intervals to be classified`
 * `Apply strand` Choose to apply (if available) or ignore strand information.
 * `Path to the reference fasta file` File containing reference genome/transcriptome.
 Required when Sequence or Secondary structure branch is selected.
 * `Path to folder containing reference conservation files` Required when Conservation score branch is selected.'Path to folder containing reference conservation files' 

*Note: When providing the sequences via FASTA file or text input, sequences shorter than the window size will be padded with Ns 
(might affect the prediction accuracy). Longer sequences will be cut to the length of the window.*

`Calculate Integrated Gradients` [Integrated Gradients](https://arxiv.org/abs/1703.01365) are available for calculation.
Ten highest scoring sequences per each class are printed at the bottom of the application.
The gradient values for each sequence are also exported for future use as last columns of the results.tsv file.
Note that calculating the integrated gradients is a time-consuming process, it may take several minutes up to few hours (depending on the number of sequences).

Sequence visualization can be used for auxiliary evaluation and debugging of the model.  
The technique is based on maximizing the difference between a baseline, and an input sequence.
That dependency is the core for the sequence attribution, and is expressed as color enhancement of each nucleobase.
The higher is the attribution of the sequence to the prediction, the more pronounced is its red color.  
On the other hand, the blue color means low level of attribution.

`Run` After all the parameters are set and selected, press the run button. 
Calculating the predictions might take minutes to hours, depending on the number of sequences, branches, hardware available etc.

Results are exported to the 'prediction' subfolder in the selected `output folder`. Information about the input sequences
are preserved in the result file (e.g. fasta header or coordinates from a bed file), while there are multiple columns with the results appended.
First, there is one column per each klass showing predicted probability of the sequence belonging to the given class.
Last result column shows the highest scoring class (do not confuse with predicted class - that is based on the user's choice of the threshold for each class).

<!--
### Development
For now, if you wish to work with the app, test or develop the code, please contact me at Slack (@Eliska), and we can discuss the details.

#### How to contribute
Following the [shared repository model](https://help.github.com/en/github/collaborating-with-issues-and-pull-requests/about-collaborative-development-models),
the process is roughly as follows:
* Download the repository using `git clone git@gitlab.com:RBP_Bioinformatics/deepnet.git`
* Depending on the situation, create your own branch or use existing feature branch for development
    * Make sure you have the latest commits `git pull` (or `git fetch` first)
    * Use existing remote branch:
        * E.g. `git checkout -b streamlit_feature`
        * When finished, push your changes `git push` 
        (again, first make sure you have all the latest commits from remote branch)
    * Or create your own branch:
        * Checkout the branch you want to fork, e.g. `git checkout development`
        * Create and checkout the new branch, e.g. `git checkout -b your_name_development`
        * When making first push from this branch, set upstream tracking 
        `git push --set-upstream origin your_name_development`
* When you want to push a group of commits from a side branch to master or another protected branch, open a pull request
    * You might add a summary for the pull request, if it helps the clarity
    * Changes in the pull request can be reviewed and discussed by other collaborators
    * After that the pull request will be merged by an admin into the chosen base branch

#### To be done
See dedicated [Trello board](https://trello.com/b/me9e2k1e/rbp-binding) for a list of current tasks, issues, and additional information.

#### Tests 
(*Not yet available*)
-->
