## Supplementary figures and images for "ENNGene: an Easy Neural Network model building tool for Genomics"

### CEITEC_logo_K-0.png

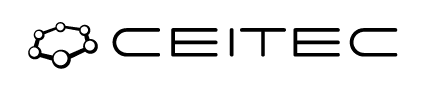

### muni-lg-eng-rgb.png

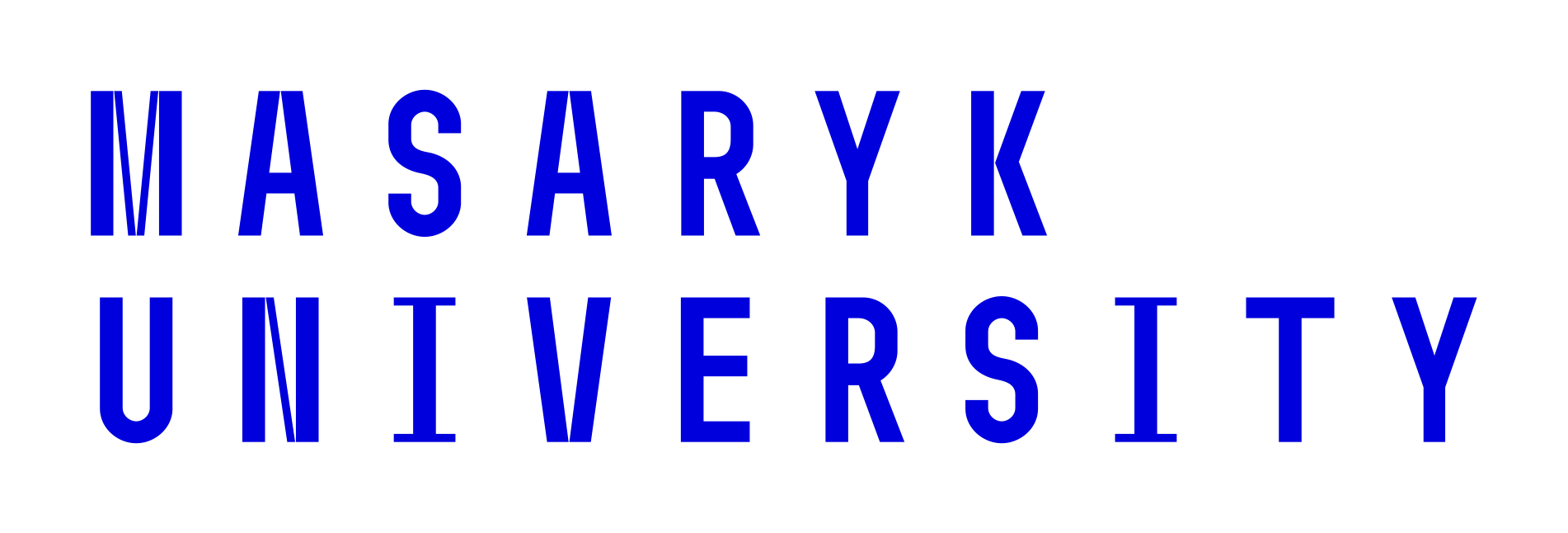

### muni-lg-rgb.png

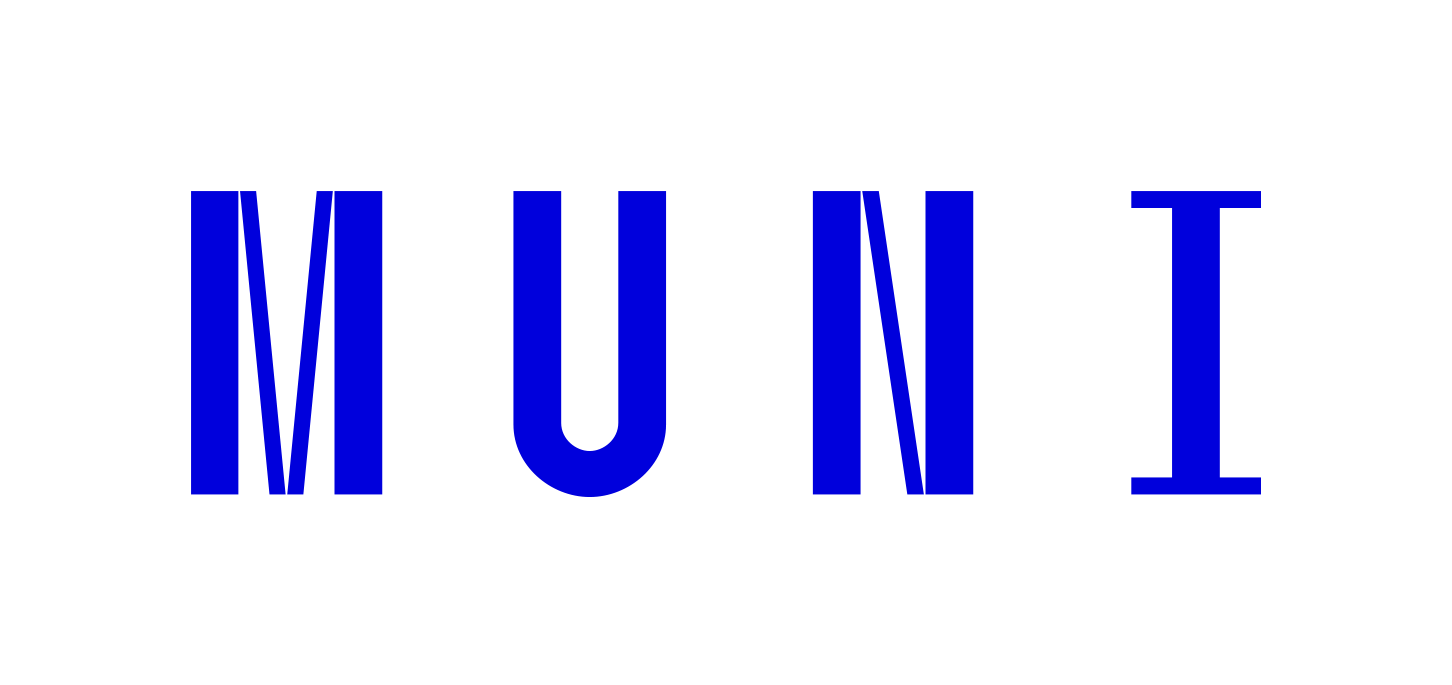
